# Structural basis of eIF4E recognition of synthetic mRNA 5′ cap analogues links cap architecture to affinity and biological activity

**DOI:** 10.64898/2026.10.05.756672

**Authors:** Sabine Schneider, Sarah Moreno, Moritz Thran, Rainer Joachim Schwarz

## Abstract

Eukaryotic mRNAs carry a 5′-terminal 7-methylguanosine (m^7^G) cap that is recognized by multiple cellular proteins and is essential for mRNA self-discrimination, stability, and translation. The eukaryotic translation initiation factor 4E (eIF4E) binds the 5′ cap and promotes recruitment of mRNAs to the ribosomal translation initiation complex, a critical step in cap-dependent translation. Beyond naturally occurring cap modifications that modulate mRNA interactions and fate, non-natural 5′ cap modifications can be used to finely tune diverse aspects of mRNA biology. This has gained particular relevance with the emergence of in vitro transcribed mRNAs as a powerful therapeutic modality for protein replacement therapies and vaccines. Here, we report binding data for a series of synthetic 5′ cap analogues interacting with eIF4E, together with X-ray crystal structures of eIF4E-cap complexes, providing detailed insights into the structural determinants of cap recognition by eIF4E. Furthermore, analysis of gene expression from mRNAs carrying different synthetic cap analogues in human cell lines and mice revealed a relationship between cap architecture and protein expression. Taken together, our structural, biochemical, and functional data establish structure-activity relationships for synthetic cap analogues and identify structural features associated with enhanced translational performance.

**GRAPHICAL ABSTRACT:** 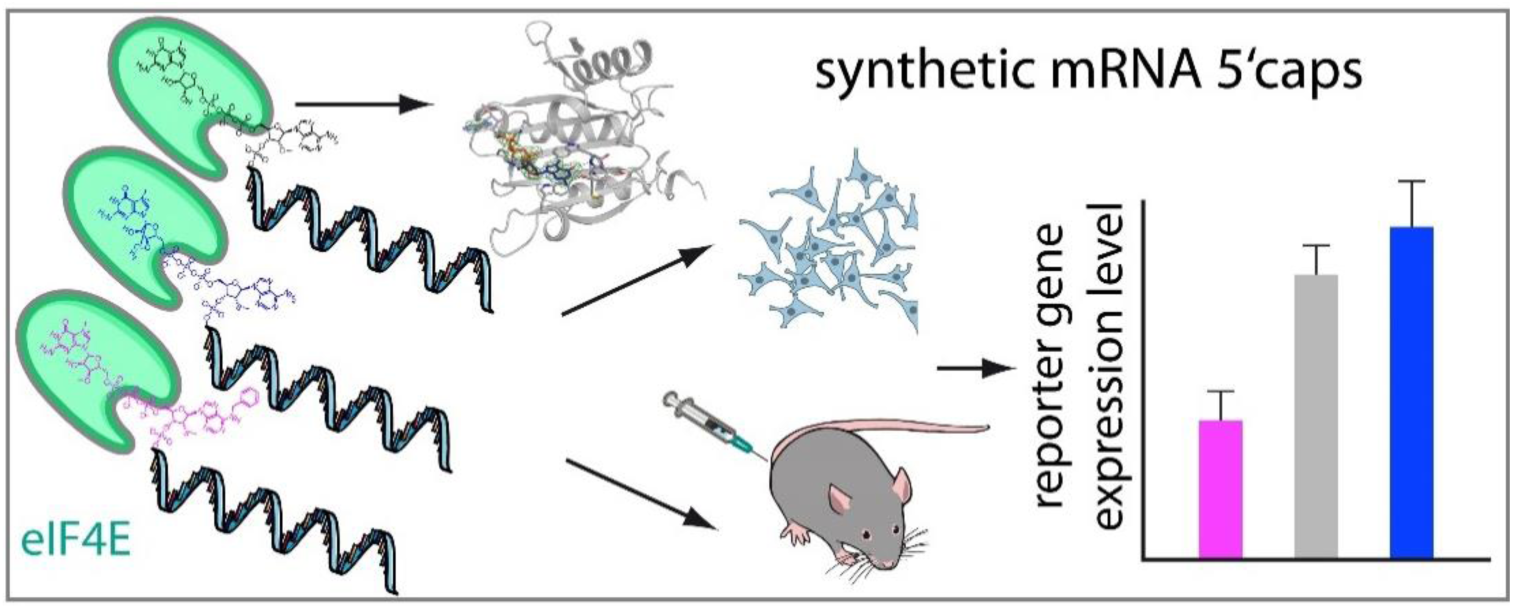

RNA icon by Servier https://smart.servier.com/ is licensed under CC-BY 3.0 Unported https://creativecommons.org/licenses/by/3.0/

NIAID Visual & Medical Arts. (10/7/2024). Lab Mouse. NIAID NIH BIOART Source. bioart.niaid.nih.gov/bioart/281

NIAID Visual & Medical Arts. (10/7/2024). Syringe. NIAID NIH BIOART Source. bioart.niaid.nih.gov/bioart/505

NIAID Visual & Medical Arts. (10/7/2024). HEK 293 Cells. NIAID NIH BIOART Source. bioart.niaid.nih.gov/bioart/198

NIAID Visual & Medical Arts. (10/7/2024). Cap Binding Protein. NIAID NIH BIOART Source. bioart.niaid.nih.gov/bioart/66

RNA CC-BY 3.0 unported

## INTRODUCTION

Messenger RNA (mRNA)-based therapeutics have emerged as a powerful platform for vaccination and therapeutic protein expression. A key advantage of this technology is its rapid adaptability to new antigens and emerging pathogen variants. However, the success of mRNA-based therapeutics critically depends on efficient, high-level, and prolonged expression of the encoded antigen or therapeutic gene. In this context, mRNA stability as well as translational efficiency and duration of expression are key determinants of therapeutic performance. These properties are modulated by the length and composition of the 3’ poly(A) tail (1,2), the 5’ and 3’ untranslated regions (UTRs) (3,4), nucleobase modifications that reduce unwanted innate immune activation, such as *N*^1^-methylpseudouridine (m^1^Ψ)(5,6) and *N*^4^-acetylcytidine (ac^4^C)(7), as well as the 5’-terminal 7-methylguanosine (m^7^G) cap structure (m^7^GpppN) (8,9).

In eukaryotes, mRNAs carry a 5′-terminal m⁷GpppN cap (where N is any nucleotide), in which the m^7^G moiety is linked to the first transcribed nucleoside via a 5′-5′ triphosphate bridge. This cap structure is essential for mRNA identity and self/non-self discrimination, protects transcripts from exonucleolytic degradation (10), and supports efficient translation (11). Cap 0 can be further modified by 2′-O-methylation (2′-OMe/N_m_) of the first transcribed nucleotide to yield Cap 1 (m^7^GpppN_m_), whereas the presence of additional 2′-OMe on the second nucleotide defines Cap 2 (m^7^GpppN_m_N_m_; Supporting Figure 1).

These 2′-OMe groups contribute to self-recognition, dampen activation of innate immune sensors, and help prevent translational shutdown. In addition, Cap 2 has been reported to enhance resistance to decapping by enzymes such as Dcp1–Dcp2 (12). In mammalian mRNAs, when the first transcribed nucleotide is adenosine, the *N*⁶ position can also be methylated to form m^6^A_m_ (m^7^Gppp^m6^A_m_N_m_) (13,14) (Supporting Figure 1), which has been reported to positively correlate with expression rates in a cell-line- and tissue-specific manner (15,16).

Recognition of the mRNA cap by the eukaryotic translation initiation factor 4E (eIF4E), which recruits capped mRNAs to the translation initiation complex, is essential for cap-dependent translation (11,17). While eIF4E also participates in additional aspects of endogenous mRNA metabolism (18), its interaction with the 5’cap is a key determinant of translation efficiency for synthetic therapeutic mRNAs (18). Thus, optimisation of the 5′ cap structure is an important strategy to improve the stability and translational performance of in vitro-transcribed mRNAs for therapeutic applications. Synthetic cap structures can be incorporated co-transcriptionally during in vitro transcription or added enzymatically to the transcribed RNA. For example, mRNA COVID-19 vaccines have used Cap 1-type structures, including enzymatically generated Cap 1 (m^7^GpppA_m_pG) used by Moderna (19) and co-transcriptionally incorporated Cap 1 3′-OMe-type analogues (m^7^G_m_pppA_m_pG) used by BioNTech (20). Several synthetic cap analogues have been reported to improve mRNA stability and gene expression, including the *N*⁶-benzylated trinucleotide cap analogue m⁷Gpppᴮⁿ⁶AₘpG, termed AvantCap (21) and cap analogues containing a 1,2-dithiodiphosphate modification (22).

To investigate how 5′ cap modifications affect eIF4E binding and influence gene expression in human cell lines and mice, we synthesized a series of di- and trinucleotide cap analogues based on eukaryotic Cap 0 and Cap 1 structures (Scheme 1 and Supporting Figure 1) and compared them with Cap 1 (m^7^GpppA_m_pG) and Cap 1-3′-OMe (m^7^G_(3′-OMe)_pppA_m_pG). The modifications included replacement of the m^7^G ribose by a diethylene glycol linker (CV-Cap1; m7G_DiEG_pppG and CV-Cap2; m7G_DiEG_pppA_m_G), substitution of the naturally occurring 2′-OMe group at the +1 nucleotide of Cap 1 by a thiomethyl group (2’SMe; CV-Cap3; m^7^GpppA_sm_pG), and rigidification of the m^7^G ribose through incorporation of a bicyclic locked nucleic acid (LNA) scaffold (m^7^G_LNA_pppA_sm_G; CV-Cap4). In addition, inspired by the previously reported AvantCap analogue, in which *N^6^*-benzylation of the first transcribed adenosine enhances mRNA stability and expression (21), we synthesized CV-Cap5 containing both an *N^6^*-benzyl substituent and a 2′SMe group (m^7^G_m_ppp^bn6^A_m_pA; Scheme 1). Using biochemical, structural, and functional analyses, we show that cap flexibility influences eIF4E binding affinity and is associated with distinct water networks in eIF4E-cap complexes. Furthermore, we identify a previously undescribed binding groove that accommodates the exocyclic benzyl group of CV-Cap5. Upon transfection of immortalized murine and human cell lines, as well as primary human cell models, with luciferase-encoding mRNAs carrying these different synthetic 5′ cap structures, clear cap-dependent differences in reporter expression were observed. Finally, the translation activity was assessed in vivo by injecting lipid nanoparticle (LNP) formulated reporter mRNAs into the *Musculus tibialis* (*M. tibialis*) of BALB/c mice and the results support the notion that reduced conformational flexibility and increased structural preorganization of the cap promote efficient translation. In summary, the structural analysis of cap recognition by eIF4E provided here, together with the in vitro and in vivo data on translation efficiency, demonstrates a clear relationship between cap architecture and translational performance. This suggests that optimization of cap conformation represents an effective strategy for improving the performance of therapeutic mRNAs.

**Scheme 1:**
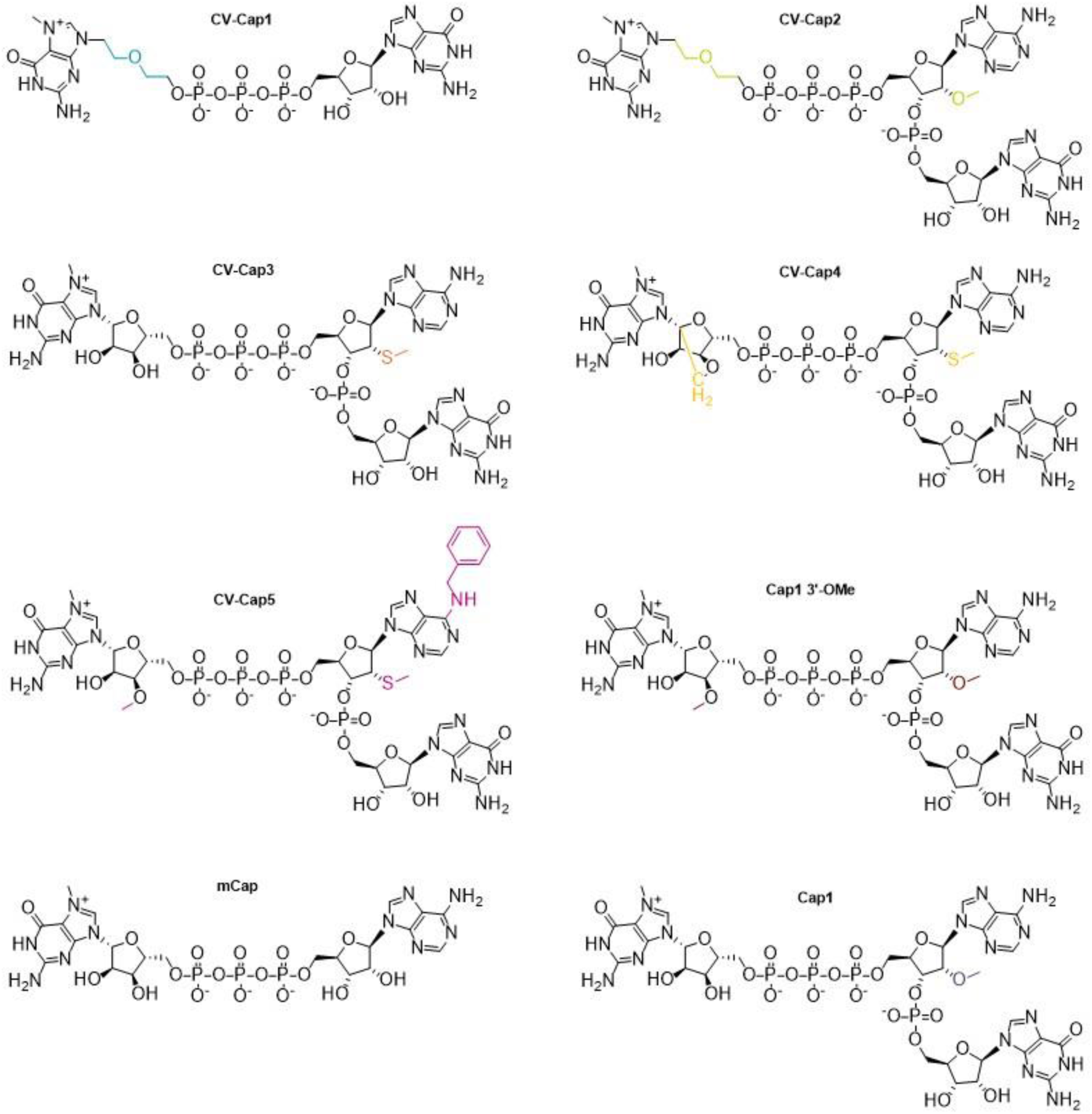
Chemical structures of cap-analogues used in this study.

## MATERIAL AND METHODS

### Protein expression and purification

#### Cloning and expression of Homo sapiens eIF4EΔ1-27

The synthetic sequence (Eurofins Genomics GmbH) encoding *H. sapiens* eIF4EΔ1-27 (Uniprot No. P06730) in frame with a N-terminal His-tag and precision protease cleavage site (Supporting Table 1), was cloned in pET28 (linearized with NcoI and HindIII) using the Hifi Assembly Kit (New England Biolabs (NEB)). Identity of the sequence was confirmed by sequencing and for protein expression the plasmid was transformed in *Escherichia coli* (*E. coli*) T7express cells (NEB). For protein expression cells obtained from an overnight culture (LB medium supplemented with 50 μg/ml Kanamycin) were diluted 1:100 in 2YT medium (50 μg/ml kanamycin) and incubated at 37 °C, 180 rpm until an OD 600 of 0.8 was reached. Protein expression was induced with 0.5 mM isopropyl-β-D-1-thiogalactopyranoside (IPTG) and cultures incubated overnight at 18°C. Cells were subsequently harvested by centrifugation (4000 × g; 20 min, RT) and cell pellets were stored at -20°C.

#### Purification of Homo sapiens eIF4EΔ1-27 for binding assays

The cell pellet obtained from 4 or 6 litre expression culture was resuspended in 40 or 60 mL lysis buffer (50 mM Tris-HCl, 500 mM NaCl, 10 mM imidazole, 2 mM tris(2-chlorehyl)phosphate (TCEP), pH 8), respectively, supplemented with DNase I (AppliChem), cOmplete TM protease inhibitors (Roche) and lysozyme (Carl Roth). Cells were lysed by homogenization using an EmulsiFlexC5 (Avestin Inc.) and cleared lysate was obtained by centrifugation (22,000 × g; 30 min; 4°C). Nucleic acids were removed from the lysate by adding dropwise polyethylenimine (8% w/v; Merck Millipore) to a final concentration of 0.5 % w/v, followed by a second centrifugation step (22,000 × g; 20 min; 4°C). The cleared lysate was filtered and incubated with 3 mL Ni-NTA Superflow beads (Cytiva) for 30°min at 4°C slowly rotating. The slurry was loaded onto a gravity flow column and the resin washed with 10 column volumes (CV) wash buffer (50 mM Tris-HCl, 500 mM NaCl, 20 mM imidazole, 2 mM TCEP pH 8). The protein was eluted with 5 CV elution buffer (50 mM Tris-HCl, 100 mM NaCl, 250 mM imidazole, 2 mM TCEP pH 8). The elution fraction was diluted 1:5 with 25 mM Hepes-KOH, pH 7.6, 10% w/v glycerol and the protein directly loaded on a ResourceS column (1 mL, Cytiva), equilibrated with 25 mM Hepes-KOH, 100 mM KCl, 2 mM DTT, 10 % w/v glycerol, pH 7.6 on an ÄktaGo FPLC system (Cytiva) at 8°C. The protein was eluted with a linear gradient (20 CV) of 25 mM Hepes-KOH, 1 M KCl, 2 mM DTT, 10% w/v glycerol, pH 7.6. Protein-containing fractions were pooled, concentrated with an Amicon Ultracell Centrifugal Filter Unit (MWCO 10 kDa, Merck Millipore), the aggregates were removed by centrifugation (15 min, 4°C, 15,000g) and the protein was loaded on a Superdex75 10/300 increase (multiple runs). Alternatively, the protein-containing fractions from the IEX column purification were pooled but not concentrated, the aggregates were removed by centrifugation (15 min, 4°C, 15,000 g) and the protein was loaded onto Superdex200 16/60 column (Cytiva). Size exclusion chromatography columns were equilibrated with 25 mM Hepes-KOH, 100 mM KCl, 2 mM DTT, 10% w/v glycerol, pH 7.6. The protein-containing fractions were pooled, and the concentration adjusted to 0.4-0.65 mg/mL, aliquoted in PCR-tubes (30 μg/aliquot), flash-frozen in liquid nitrogen and stored at −80°C. Quantification was done by UV/Vis spectroscopy with a Denovix DS-11 (Biozym) and N60 (Implen) nanophotometer, using a millimolar extinction coefficient of ε_280nm_ = 2.4 mL x mg^-1^ x cm^-1^. Protein yield was ∼1 mg/L expression culture. Since eIF4E tends to aggregate, glycerol was added to all buffers after the His-tag purification step and the protein was not refrozen.

For complex crystallization of eIF4E, cell lysis and His-tag purification were carried out as described above. However, for crystallization the His-tag was removed. Therefore, the protein-containing elution fractions from the Ni-NTA column were pooled and the buffer was exchanged to 25 mM Hepes, 100 mM KCl, 2 mM DTT, pH 7.6 using a HighPrep 26/10 desalting column (Cytiva) attached to an ÄktaGo FPLC system at 8°C. For tag removal Precission protease (lab made) was added (1:100) to the protein solution and incubated over night at 4°C. At the next day, the protein was loaded on a ResourceS column (1 ml, Cytiva) equilibrated with 25 mM Hepes-KOH, 100 mM KCl, 2 mM DTT, pH 7.6. The protein was eluted with a linear gradient (20 CV) of 25 mM Hepes-KOH, 1 M KCl, 2 mM DTT, pH 7.6. Protein-containing fractions were pooled, concentrated with an Amicon Ultracell Centrifugal Filter Unit (MWCO 10 kDa) and aggregates were removed by centrifugation (15 min, 4°C, 15,000 g). The protein was loaded on a Superdex75 10/300 increase column (Cytiva), equilibrated with 25 mM Hepes-KOH, 100°mM KCl, 2°mM DTT, pH 7.6. The pure protein was concentrated to ∼1 mg/mL and directly used for complex formation and crystallization.

### Affinity assay

For binding studies no differences between His-eIF4EΔ1-27 or tag-less eIF4EΔ1-27 were observed (data not shown). Thus His-eIF4EΔ1-27 was used for all subsequent assays.

Dissociation constants of cap analogues for the translation initiation factor eIF4E were determined using a fluorescent competitive binding assay (Py-FLINT-B) as described by Kasprzyk et al. (23) with Cap 1 serving as the reference ligand. Measurements were carried out in black, non-binding 96-well plates on a Tecan Spark plate reader. The assay buffer consisted of 50 mM HEPES-KOH (pH 7.2), 100 mM KCl, and 0.5 mM ethylenediaminetetraacetic acid (EDTA). Final assay mixtures (200 µL per well) contained pyrene probe (10 nM), eIF4EΔ1-27 (75 nM), and the respective cap analogue at the indicated concentration. Negative-control wells contained pyrene probe in assay buffer in the absence of eIF4EΔ1-27 (10 nM final pyrene concentration). Primary cap stock solutions were prepared by diluting 2 µL of a nominally 100 mM cap solution with 1579 µL of assay buffer to give a nominal concentration of 63.25 µM. The actual concentration of each stock solution was determined in triplicate by UV/Vis spectroscopy (NanoDrop) at 255 nm using the corresponding molar extinction coefficient. These experimentally determined concentrations were used to calculate correction factors for subsequent dilution steps. Working solutions for the dilution series were prepared in assay buffer from the quantified primary stock solutions. A freshly prepared mastermix containing eIF4EΔ1-27 and pyrene probe was used for each independent experiment. Prior to mastermix preparation, eIF4EΔ1-27 was diluted with 50 µL of protein buffer (20 mM HEPES-KOH pH 7.2, 2 mM DTT, 100 mM KCl, 10% v/v glycerol) and subjected to centrifugal filtration through a 0.45 µm PVDF Durapore filter. An additional 50 µL of protein buffer was applied to the filter and centrifuged. The concentration of the combined eluate was determined spectrophotometrically at 280 nm, and the eluate was then diluted with assay buffer and supplemented with pyrene probe to give a mastermix containing 150 nM eIF4EΔ1-27 and 20 nM pyrene probe. For the assay, 100 µL of each ligand dilution was transferred to the corresponding wells, followed by addition of 100 µL of the eIF4E/pyrene-probe mastermix, affording a final volume of 200 µL per well. Plates were incubated at 30 °C with orbital shaking (360 rpm, 900 s). Fluorescence was recorded at 30 °C using an excitation wavelength of 345 nm and an emission wavelength of 378 nm (10 nm bandwidth each), with the detector gain set to 150, 40 flashes per well, an integration time of 40 µs, and a z-position of 20,000.

Concentration-response curves were fitted in GraphPad Prism to obtain EC₅₀ values according to Equation 1, where *X* is the ligand concentration, EC₅₀ is the concentration resulting in half-maximal response, *Bottom* represents the signal of the free probe (negative control), and *Top* corresponds to the fully bound probe-protein complex.

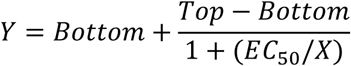

### *Homo sapiens* eIF4EΔ1-27 complex crystallization X-ray data collection, processing and structure refinement

Cap-analogues were added to a solution of eIF4EΔ1-27 protein (1 mg/ml ∼ 30 μM protein) to a final concentration of 300 μM (∼1:10 equivalents protein:cap). The protein-cap complex was then concentrated to 4-5 mg/mL using centrifugal filter devices, centrifuged (10 min, 4°C at 15,000 g) and high-throughput crystallizations were set up using the Oryx8 crystallization robot (Douglas Instruments) in different protein to precipitant ratios (1:2; 1:1; 2:1). The crystallization plates were incubated either at 18°C or 8°C and crystals usually appeared after ∼1-2 weeks. Crystallisation conditions for the different complexes are listed in Supporting Table 2. Diffraction data were collected at the European Synchrotron Radiation Facility (ESRF, ID30A-3) and Deutsches Elektronen Synchrotron (DESY, P13 and P14). The data were processed with XDS (24) and the resolution cut-off at was chosen according to the correlation coefficient of random half-data sets (CC 1/2) at about 50 % (25–27). To increase data multiplicity for the Cap1 and CV-Cap2 complexes, data sets collected on two crystals were combined. The structure was solved using the coordinates of the *H. sapiens* eIF4E structure (PDB code 1IPB) as molecular replacement model in PHASER 2.8.3 (28). The model was completed by iterative cycles of manual model building in COOT 0.9.8.96 (29) and restraint refinement in REFMAC5.8.0431 (30). Structure optimisation was carried out using the PDBredo server (31). For data processing and structure refinement statistics see Supporting Table 3. All structural figures were prepared with Open Source PyMol (Schodinger, LLC). Crystallographic data of the eIF4E complex structures have been deposited in the Protein Data Bank (https://www.ebi.ac.uk/pdbe/) under the PDB accession codes: 34AK (Cap 1); 34CV (CV-Cap1); 34CO (CV-Cap2, P1); 34DD (CV-Cap2, P2_1_), 34CY (CV-Cap4), 34DH (CV-Cap5).

### mRNA design and synthesis

A DNA sequence comprising the HSD17B4 5′-UTR, the photinus pyralis luciferase coding sequence, the PSMB3 3′-UTR, a histone stem-loop structure and a 100-nt poly(A) tail was cloned into a modified pUC19-derived vector (Supporting Table 4). The plasmid DNA was propagated in bacteria using common protocols and plasmid DNA was extracted, purified, and enzymatically linearized using a restriction enzyme. mRNAs were transcribed from linearized plasmid DNA (50 µg/mL) at 37°C for 3 h using T7-based in vitro transcription systems in the presence of the respective cap analog. Transcription conditions were optimized for each cap analog and are summarized in Supporting Table 5. Following transcription, template DNA was digested with Pulmozyme (2500 U/mL) in the presence of 3.2 mM CaCl₂ for 30 min at 37°C. mRNAs were purified by reversed-phase chromatography using a PLRP-S stationary phase and an acetonitrile gradient in triethylammonium acetate buffer (32).

### Cell cultivation, transfection and Luminescence measurements

#### Cell lines

Cells were cultured at 37 °C and 5% CO₂ and seeded in 96-well plates. 10,000 HDF cells (Promocell) were seeded in fibroblast growth medium (Promocell), 20,000 HepG2 cells (DSMZ) and 50,000 J774A.1 cells (ATCC) were seeded in DMEM (ATCC). All media contained antibiotics throughout culture and transfection. Cells were seeded 24 h prior to transfection in a total volume of 200 µL per well. On the day of transfection, the complete medium was replaced with serum-free Opti-MEM (Gibco).

mRNAs were complexed with Lipofectamine2000 or Lipofectamine MessengerMAX according to the manufacturer’s instructions in Opti-MEM and incubated for 20 min at room temperature prior to addition to the cells. Lipocomplexes were added to cells at a dose of 5 ng or 50 ng mRNA per well in a total volume of 200 µL. Cells were incubated with the complexes for 90 min, after which the transfection mixture was completely removed and replaced with 200 µL fresh complete medium per well. Cells were then further incubated at 37 °C and 5% CO₂ until harvesting. 100 µL of 1x passive lysis buffer (Biotium) was added to each well. Cells were shaken for 15 min at room temperature and then stored at -80°C for at least 1 h. After thawing, 20 µL of lysates were used to detect and measure luciferase activity via chemiluminescence using adenosine triphosphate (ATP) and D-luciferin in a Beetlejuice buffer system (p.j.k.). To this end, plates were placed in a plate reader (Tecan Spark) with injection device for Beetle-juice containing substrate for firefly luciferase. Per well, 50 µL of beetle-juice were added. Raw data containing relative light units were used to plot differences between mRNAs derived from cap analogues.

#### Primary cells

Primary human hepatocytes (pHH) and human skeletal muscle cells (HSkM) were incubated at 37 °C and 5% CO₂ and seeded in 96-well plates. HSkM cells were seeded at 48,000 cells per well in 200 µL differentiation medium (DMEM supplemented with 2% horse serum) 48 h prior to transfection. pHH cells were seeded at 40,000 cells per well in 200 µL plating medium (Williams’ E medium supplemented with hepatocyte plating supplements) 24 h prior to transfection on collagen-coated plates. On the day of transfection, culture medium was carefully removed and replaced with serum-free Opti-MEM. mRNAs were diluted in water and complexed with Lipofectamine MessengerMAX in Opti-MEM according to the manufacturer’s instructions. Complex formation was performed by sequential dilution of mRNA and transfection reagent in Opti-MEM followed by mixing and incubation prior to addition to cells. Transfections were performed using 50 ng mRNA per well. Lipocomplexes were added dropwise to the cells and incubated for 90 min at 37 °C and 5 % CO₂. After incubation, the transfection mixture was completely removed and replaced with 200 µL of fresh culture medium per well. Cells were further incubated under standard conditions until harvesting. At 24 and 48 h post transfection, cells were lysed by addition of 100 µL of 1× passive lysis buffer per well, followed by incubation at room temperature with shaking for 15 min and storage at −80 °C prior to analysis. Luciferase activity was quantified after thawing according to the protocol of cultured cell lines.

### Formulation

mRNA was encapsulated into lipid nanoparticles (LNP) through a self-assembly process where an ethanolic lipid solution of an ionizable cationic lipid, 1,2-distearoyl-sn-glycero-3-phophocholine, cholesterol, and a polyethylene glycol-lipid was rapidly mixed with an aqueous solution of mRNA at acidic pH.

For formulations containing lipids from Acuitas Therapeutics (Vancouver, Canada), the ionizable cationic lipid (proprietary to Acuitas) and LNP composition are described in patent applications WO 2017/075531A1WO and WO 2017/004143, respectively. (Formulations were prepared by CureVac.)

For formulations containing SM-102 as ionizable cationic lipid, the LNP composition was SM-102: DSPC: Cholesterol: PEG2000-DMG (50: 10: 38.5: 1.5). (Formulations were prepared by Merck (Darmstadt, Germany).)

### Animal experiments

All animal experiments were performed at Preclinics Gesellschaft für präklinische Forschung mbH (Preclinics, Potsdam, Germany) in accordance with the German Animal Welfare Act. Female BALB/c mice were obtained from Charles River Laboratories, acclimatized for at least one week, and entered the studies at 6-8 weeks of age. Mice were assigned to treatment groups on a cage-wise basis and distributed into two independent cohorts that were terminated 6 h or 24 h after administration. Each cohort comprised six animals. Both the left and right *M. tibialis* muscles were injected with the same formulation and analyzed as individual samples. No animals or samples were excluded from the analysis. Blinding was not performed. mRNA-LNP formulations were thawed to room temperature, diluted in 0.9% NaCl to the target dose, and mixed by gentle pipetting prior to administration. Mice were anesthetized with isoflurane before intramuscular injection. For the experiments shown in Figure 4A, mRNAs were formulated into an ALC-315-based LNP formulation (derived from Acuitas) and administered at a dose of 1 μg mRNA in 20 μl per *M. tibialis* muscle on each side. For the experiments shown in Figures 4B and 4C, mRNAs were formulated in SM-102 LNPs (Merck) and administered at doses of 0.2 μg or 1 μg mRNA in 20 μL per *M. tibialis* muscle on each side, respectively. Intramuscular injections were performed into the left and right *M. tibialis* muscles. At 6 h or 24 h post-administration, animals were euthanized by cervical dislocation. Muscles were harvested immediately and snap-frozen in liquid nitrogen.

For luciferase analysis, frozen muscle samples were lysed with a steel bead in 2 mL safe-lock tubes using a Tissue Lyser II (Qiagen) for 3 min at full speed. Passive lysis buffer (1×, 500 µL) was added to each sample, followed by an additional tissue-lysis step for 6 min at full speed. Samples were centrifuged at 13,500 rpm for 10 min at 4 °C, and the supernatants were transferred to 96-well plates and stored at −80°C until measurement. Lysates were thawed under a sterile bench and immediately placed on ice. PpLuc standards were prepared from a 100 ng/mL aliquot by serial 10-fold dilution in 1× lysis buffer down to 0.01 ng/mL. Lysates and standards were transferred in 20 µL aliquots to white LIA plates and measured on a Tecan Spark plate reader using a standard PpLuc protocol comprising injection of 50 µL Beetlejuice reagent containing D-luciferin and ATP per well, a 0.5 s delay, and a 2 s measurement. Raw luminescence values were normalized to individual muscle weights.

### Chemical synthesis

#### General

Unless otherwise specified, all starting materials are obtained from commercial suppliers in the highest purity available. If not stated otherwise, all reactions are conducted at room temperature, ambient atmosphere and as indicated in the individual procedures.

The temperatures are expressed as °C. Solvent removal was carried out using rotary evaporation at 40°C. Mass spectrometry (MS) was conducted using an Exploris 240 (Thermo Fisher) with HESI ion source and orbitrap analyser or an ISQ-EM Quadrupol mass detector (Thermo Fisher). ESI-MS using both MS systems was carried out in positive and negative mode using 5 mM ammonium acetate + 90% methanol, 50 mM ammonium acetate + acetonitrile and 5 mM ammonium acetate + acetonitrile. NMR spectra were recorded using a Bruker Avance III HDX 400 with a 5 mm BBFO sample head, a Bruker Avance II – Ultrashield 400 MHz, a Bruker Avance Neo – Ascend 400 MHz or an Magritek Ultra 80 MHz spectrometer. Chemical shifts (δ) are reported in ppm relative to the residual solvent signal. ^1^H-NMR data are reported as follows: chemical shift (multiplicity, number of hydrogens). Multiplicity is abbreviated as follows: s (singlet), d (doublet), t (triplet), q (quartet), p (quintet) m (multiplet), br (broad signal). For chiral compounds the sum of the integrals for both isomers is reported. Additionally, splitting of peaks for chiral compounds may be observed which is mostly reported as multiplet. All cap analogues were quantified using a Nanodrop 2000c or a NanodropOne UV/Vis spectrophotometer (Thermo Fisher) in UV/Vis mode with automatic path length option. Cap analogues were dissolved in 1 mL of water. A 1 µL aliquot was diluted 1:10 and 1:100 and measured at 255 nm for Cap 1 and Cap 2 analogues. Concentration and substance amount was calculated according to lambert-beers law using the respective extinction coefficient stated in the individual procedures. If no specific extinction coefficient is mentioned, then 30539 L/mol*cm for Cap 1 and 21000 L/mol*cm for Cap 0 is used.

For detailed protocols for the chemical synthesis and NMR spectra see the Supporting Information.

## RESULTS

### Synthesis of 5’cap analogues and expression and purification of eIF4E

To investigate the impact of 5’ cap modifications on the binding to eIF4E as well as gene expression in human cell lines and mice, we synthesised different di- and tri-nucleotide cap analogues of the eukaryotic Cap 0 and Cap 1 (Scheme 1, Supporting Figure 1 and Supporting Information) and compared them to Cap 1 and Cap 1-3′-OMe. In the CV-Cap1 and CV-Cap2 structures we replaced the m^7^G ribose with a diethylene linker which has the potential to alter interactions with decapping enzymes. In addition, we explored the impact of the 2’ modification by replacing the naturally occurring 2′-OMe moiety at the first transcribed nucleotide in Cap 1 by a 2’SMe group (CV-Cap3), as well as rigidifying the m^7^G ribose with a bicyclic LNA structure (CV-Cap4) additionally, which has been previously described beneficial for gene expression (33,34). Further, it was previously shown that bulky-adduct modification at the exocyclic N^6^ of the first transcribed adenine nucleotide improve mRNA stability and gene expression (21). Therefore, we synthesized CV-Cap5, which contains both a *N^6^*-benzyl substituent and a 2′SMe group (Scheme 1).

For the biochemical and structural analysis of eIF4E binding to different cap analogues, we cloned, expressed and purified human eIF4E without the first 27 N-terminal residues (eIF4EΔ1-27), equipped with a N-terminal His_6_-tag and HRV 3C protease cleavage site (Supporting Information and Supporting Table 1). Previously, either full-length mouse or human eIF4E, as well as N-terminal truncated versions, were used for biochemical and structural studies, showing that the N-terminus is disordered in crystal structures (35–41) and important for interaction with binding partners such as eIF4G and eIF4E binding protein 1 (4E-BP1) (42–44). Human and mouse eIF4E share an overall sequence identity of 98.2% and differ only in 4 amino acids, 3 of which are located in the first 17 N-terminal residues (Supporting Figure 2).

### Binding affinity of human eIF4E to cap analogues

In order to analyse the binding of the various synthetic cap analogues to eIF4E, their dissociation constant from purified human eIF4E was determined using a fluorescent competitive binding assay (Py-FLINT-B)(23). For comparison the commercially available Cap 1, Cap 1-3′-OMe as used in the SARS-CoV2 mRNA vaccine Comirnaty (BioNTech), as well as the dinucleotide mCap (m^7^GpppG), a dinucleotide Cap 0 analogue, were used (Scheme 1, Figure1A and Supporting Figure 3). Overall, the dinucleotide caps mCap and CV-Cap1 showed the lowest binding affinity (∼3.5-5 fold less), which may reflect the reduced interaction surface of dinucleotide and compared to trinucleotide cap analogues. Moreover, the Cap 1-3′-OMe possess an approximately two-fold lower binding affinity than the reference Cap 1. Among the trinucleotide cap-analogues CV-Cap2, with its ethylene glycol linker, displays the lowest affinity (∼3.5 fold lower), which is likely due to the higher flexibility of ethylene glycol compared to cyclic ribose and the associated higher entropic cost upon binding. CV-Cap3 and CV-Cap5 showed comparable binding affinities to Cap 1. While CV-Cap3 carries a 2’SMe group at the second adenine nucleotide, Cap 1 has here a 2′-OMe group. In addition, to the 2’SMe group of the A, CV-Cap5, carries a benzyl moiety at the N*^6^* of A as well as 3′-OMe group at the m^7^G. The highest binding affinity was observed for CV-Cap4 (m^7^G_LNA_pppA_sm_pG), which is characterised by the LNA modified ribose at the m^7^G which likely reduces the entropic cost due to its added rigidity.

### Crystal structures of eIF4E in complex with cap-analogues

The crystal structures of human and murine eIF4E in complex with m^7^G nucleotides and analogues and peptide fragments of binding partners, such as eIF4G and 4E-BP, as well as inhibitors have been reported (35–40,42–46). However, only one structure of the human eIF4E with a dinucleotide cap, m^7^GpppA has been described (PDB code 1IPB (36)).

In order to elucidate the molecular binding mode of the di- and trinucleotide cap-analogues investigated here to eIF4E, we crystallised and determined the X-ray crystal structures of human eIF4EΔ1-27 in complex with Cap 1, CV-Cap1, CV-Cap2, CV-Cap4 and CV-Cap5 to resolutions ranging from 1.75-2.1 Å. In all complex structures eIF4E can be superimposed with an r.m.s.d of about 0.15-0.3Å.

First, eIF4E was crystallized in complex with the trinucleotide Cap 1 and its structure was determined to 2.1 Å resolution in the space group P1. The asymmetric unit of the crystal contained 4 protein complexes, with two of the four molecules having higher intrinsic flexibility and thus lower local resolution, as indicated by the B-factors (Supporting Figure 4). In the two eIF4E-molecules with overall lower B-factors the m^7^GTP-moiety of the trinucleotide cap is well defined in the electron density, while the first and second transcribed nucleotide are flexible (Figure 1B and Supporting Figure 5). In the complex structure of human eIF4E and m^7^GpppA dinucleotide (PDB code 1IPB, space group P4_3_), the A is defined in the electron density due to its interaction with the C-terminal loop residue 204-210 that is stabilized by crystal contacts providing the molecular basis for the preference of an adenine base as the first transcribed nucleotide position (36). In the here reported complex of eIF4E and Cap 1 in the P1 space group this loop does not form crystal contacts and is therefore flexible (Supporting Figure 5).

**Figure 1:**
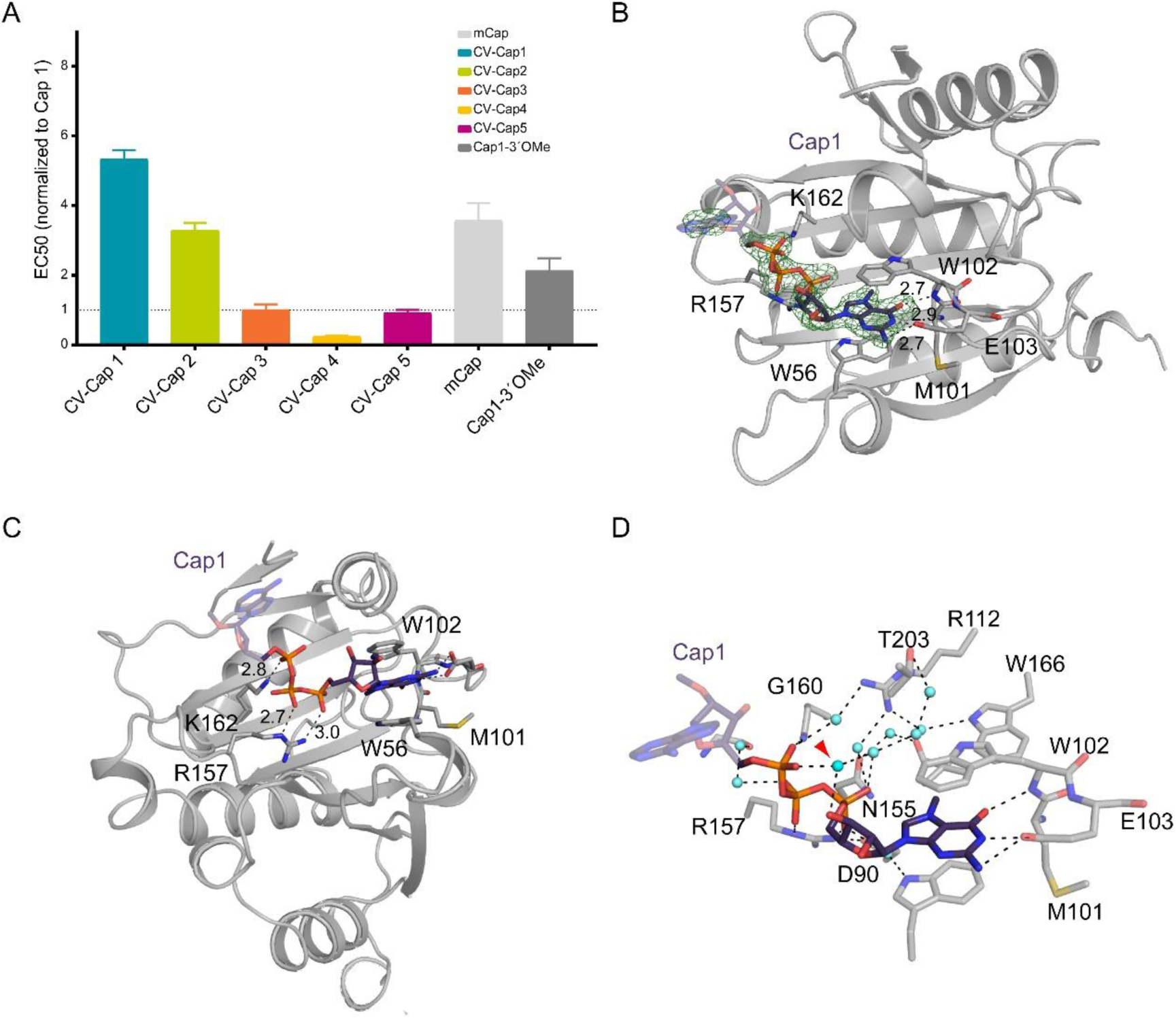
Binding affinity of eIF4E to cap-analogues and structure of eIF4E in complex with Cap 1. A) EC50 values of cap analog binding to eIF4E determined by the Py-FLINT-B competitive fluorescence assay and normalized to Cap 1. Data are shown as mean ± standard deviation from three independent experiments (n = 3). B+C) Binding mode of Cap 1 (purple stick model) to eIF4E (grey ribbon) with residues forming direct interactions shown as stick model. The simulated annealing omit-electron density of Cap 1 contoured at 3σ is shown as green mesh. D) Water-mediated contacts between eIF4E and Cap 1, with the water bridging the m^7^G-3’OH and the γ-phosphate highlighted with a red arrow.

The m^7^G moiety of the cap is sandwiched between Trp 56 and Trp 102, with the Watson-Crick-Franklin face forming hydrogen bonds with the carboxylic acid group of Glu 103 and the backbone amide of Trp 102 (Figure 1B), which is common to all m^7^G-eIF4E complexes. The α- and β-phosphates of the m^7^G-triphosphate bridge form hydrogen bonds with HN_ε_ and HN_η2_ in the guanidino group of Arg 157, while Lys 162 interacts with the γ-phosphate (Figure 1C). in addition, a dense network of water mediated contacts, including a water-bridged interaction between the m^7^G 3’ OH and α-phosphate can be observed (Figure 1D and Supporting Figure 5). The importance of water mediated as well as conserved water position in eIF4E crystal structures has been previously discussed (41).

The complex structures of eIF4E with the diethylene glycol di- and trinucleotide cap analogues CV-Cap1 and CV-Cap2 were also determined in the space group P1 to 1.75Å and 2.1Å resolution, respectively (Figure 2A). The crystal packing, with four protein complexes in the asymmetric unit of the crystal is comparable to the eIF4E-Cap 1 complex, with two complexes having higher intrinsic flexibility and lower local resolution (Supporting Figure 6). Again, the C-terminal loop 204-210 is flexible and the first (and second) transcribed nucleotides are not defined by the electron density, with an equivalent binding mode of CV-Cap1 and CV-Cap2 (Supporting Figure 7). However, the diethylene and triphosphate linker are well defined by the electron density (Supporting Figure 7). While the m^7^G forms the same interactions common to all eIF4E m^7^G complexes as described above, the positioning of the phosphate linker is shifted, likely due to the higher flexibility of the ethylene glycol compared to the ribose ring (Figure 2A). Moreover, the complex network of water mediated contacts is also altered. This reduction of binding interactions, in addition to the higher entropic cost, explains the observed lower affinity of CV-Cap1 and CV-Cap2 compared to Cap 1 and Cap 1-3′-OMe.

**Figure 2:**
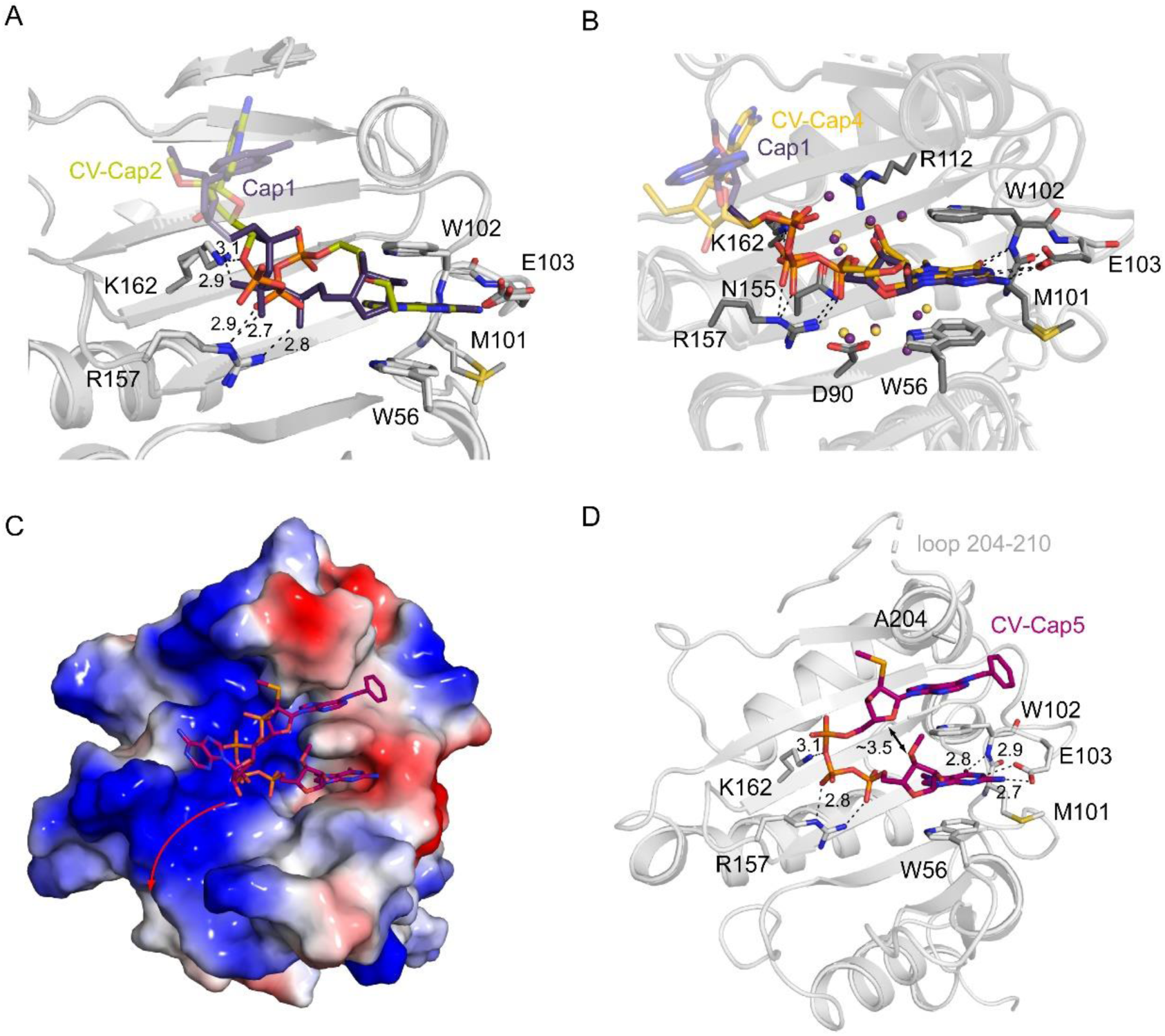
Structure of eIF4E in complex with CV-Cap2, CV-Cap4 and CV-Cap5. A) Binding mode of CV-Cap2 (light green stick model) to eIF4E (light grey ribbon) in the crystal space group P1. Residues forming direct interactions are shown as stick model. Superimposed is the eIF4E/Cap 1 complex (dark grey/purple), highlighting the shifted position of the triphosphate bridge and the reduced number of direct contacts. B) Superposition of the eIF4E complexes with CV-Cap4 (light grey/golden) and Cap 1 (dark grey/purple). The water molecules at structurally equivalent positions are shown as yellow (CV-Cap4 complex) and purple (Cap 1 complex) spheres, respectively. C) Surface representation of eIF4E in complex with CV-Cap5. pink stick-model), with the electrostatic potential mapped onto the surface of the protein. The red arrow indicates the likely position of the next nucleotides in a mRNA. D) Interactions between eIF4E (grey ribbon representation) and CV-Cap5, with the *N^6^*-butyl-2’SMe-A sandwiched between Trp 102 and the C-terminal loop 204-210. Protein residues directly interacting with the cap are shown as grey stick model.

In addition, the structure of eIF4E-CV-Cap2 complex could also be determined in the space group P2_1_ to 1.75 Å resolution, with four protein complexes in the asymmetric unit of the crystal. Binding of the m7G moiety is analogous to all other reported structures. However, here crystal packing contacts stabilize the C-terminal loop 204-210 as well as the first transcribed nucleotides of two complexes by stacking interactions (Supporting Figure 7). Furthermore, the second transcribed nucleotide is also defined in the electron density (Supporting Figure 7). Due to this stabilization of the first transcribed nucleotide, the triphosphate linker in this eIF4E CV-Cap2 complex structure in the P2_1_ symmetry occupies the equivalent position as in the Cap 1 or m^7^GpppA complexes, but again fewer water-mediated contacts can be observed (Supporting Figure 7).

Complex crystals of eIF4E and CV-Cap3 diffracted X-rays to about 2.9 Å resolution and belonged to the P1 symmetry. The only difference between CV-Cap3 and Cap1 is the thiomethyl-instead of a methoxy group at the 2’ position of the first transcribed nucleotide, which is flexible in the P1 symmetry and thus not defined in the electron density. Therefore, no model was built and refined for this complex.

Next, the crystal structure of eIF4E in complex with CV-Cap4, was determined to 2 Å resolution The crystals belonged again to the space group P1, with four complexes in the asymmetric unit of the crystal lattice. As observed for all here obtained crystal structures in this space group, two of the protein complexes show higher B-factors, with lower local resolution. The m^7^G and the triphosphate linker as well as the water network resemble the Cap 1- and m^7^GpppA-complex structures, with the locked ribose of the m^7^G aligning with the ribose in the Clean Cap or m^7^GpppA (Figure 2B and Supporting Figure 8). The high binding affinity of this cap-analogue is likely due to the dense interaction network as observed for the natural cap structure in combination with a reduced entropic cost due the locked ribose.

Crystals of eIF4E in complex with CV-Cap5 belonged to the space group P2_1_ and the coordinates of the structure were refined to 2 Å resolution. The two complexes in the asymmetric unit of the crystal lattice can be superimposed with an r.m.s.d of 0.08Å^2^. The trinucleotide cap-structure, as well as the protein N-terminus are fully defined in the electron density (Supporting Figure 9). Residues of the C-terminal loop 204-210 are partly ordered and the protein N-terminus and second transcribed nucleotide is involved in crystal packing interactions (Supporting Figure 9). Again, the m^7^Gm and the triphosphate linker are held equivalently to the Cap 1 and m^7^GpppA complexes. However, the *N*^6^-butyl-2’SMe-A, at the position of the first transcribed nucleotide, is bound in a novel binding position and packs between Trp 102 and the α-helix formed by residues 200-204 of the C-terminal loop (Figure 2C+D). In addition, the 3′-OMe of the m^7^G_m_ relatively close (∼3.5 Å) to the ribose of the *N^6^*-butyl-2’SMe-A (Figure 2D) and fewer water-mediated contacts can be observed (Supporting Figure 9). The entropic cost associated with the displacement of water molecules and the proximity of the 3′-OMe and ribose in the binding position, might explain that the binding affinity of this cap-analogue is comparable to Cap 1.

### Translation efficiency in various cells

Next, we investigated the impact of the cap-structure on protein levels in various immortalized murine and human cell lines, as well as primary human cell models. Upon transfection with luciferase-encoding mRNAs carrying different synthetic 5′ cap structures, clear cap-dependent differences in reporter expression were observed (Figure 3). Consistent with their reduced affinity for eIF4E, mRNAs capped with analogues containing flexible diethylene glycol linkers (CV-Cap1+2) exhibited lower luciferase expression than mRNAs capped with Cap 1 or Cap 1 3′-OMe. In contrast, replacement of the natural 2′-OMe group at the first transcribed nucleotide with a 2′SMe substituent (CV-Cap3) supported reporter expression that was, depending on the cellular model, comparable to that obtained with the commercial reference caps (Figure 3). Addition of the exocyclic benzyl-moiety to the first transcribed adenine nucleotide (CV-Cap5) showed, depending on the cells used, lower or similar expression levels than Cap 1 and Cap 1-3′-OMe (Figure 3B, C). Notably, the LNA-modified cap analogue (CV-Cap4) showed a tendency toward higher reporter expression (Figure 3B, C), suggesting that conformational restriction of the m⁷G ribose may enhance translational output in cellular systems.

**Figure 3:**
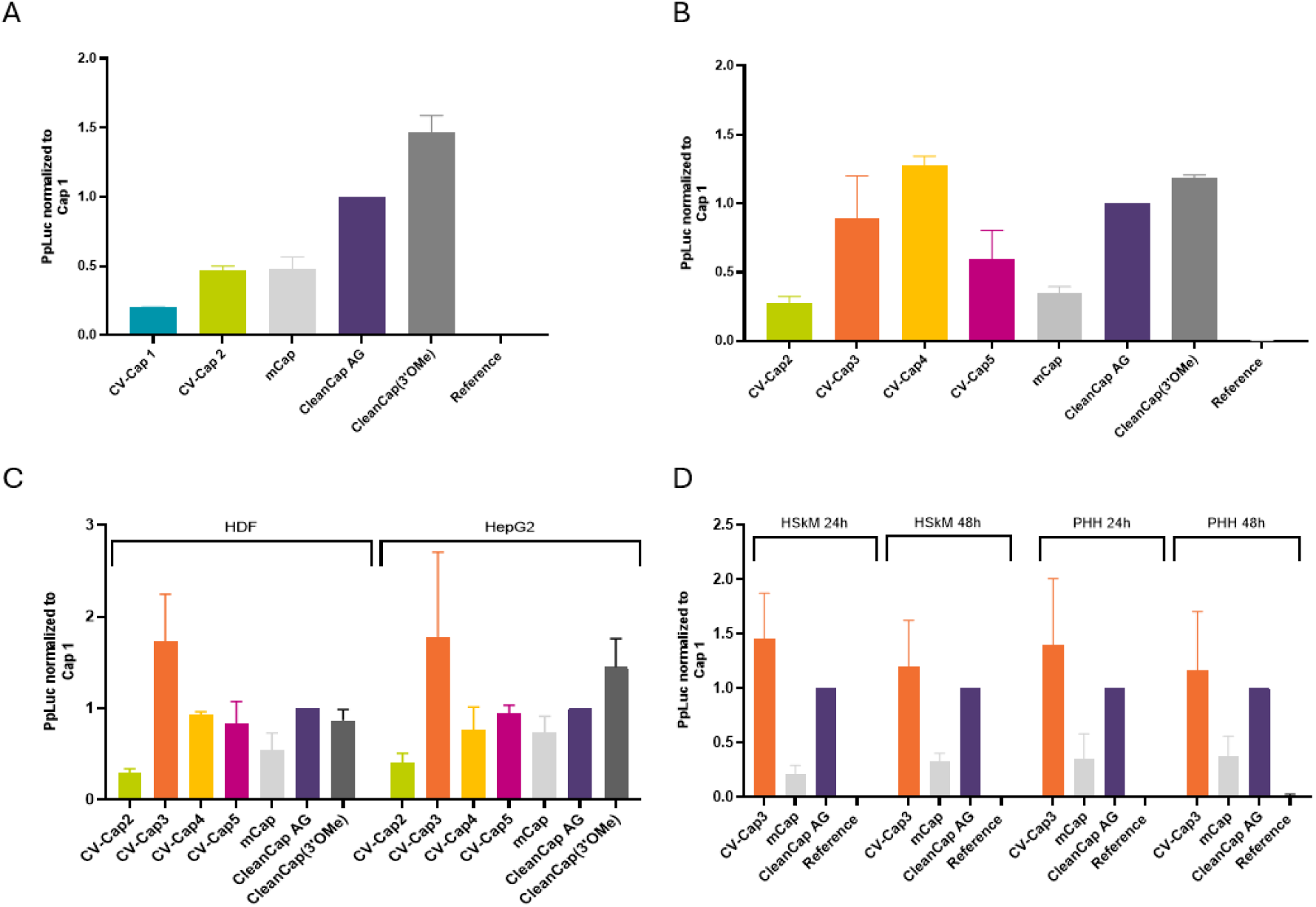
Cell-based translation activity of capped m^1^Ψ-modified mRNAs. PpLuc activity was determined after transfection of m^1^Ψ-modified mRNAs formulated with Lipofectamine 2000 (A) or Lipofectamine MessengerMAX (B–D) and normalized to Cap 1. A) J774A.1 cells, 50 ng mRNA, mRNAs carrying CV-Cap1 or CV-Cap2 versus commercial cap analogues. B) J774A.1 cells, 5 ng mRNA, mRNAs carrying CV-Cap2-4 versus commercial cap analogues. C) HDF and HepG2 cells, 5 ng mRNA, mRNAs carrying CV-Cap2–5 versus commercial cap analogues. D) Primary HSkM and PHH cells, 50 ng mRNA, mRNAs carrying CV-Cap3 versus commercial cap analogues. For panels A–D, data are shown as mean ± standard deviation (n = 3).

### Translation efficiency mouse model

To determine whether these effects translate to an in vivo setting, translation activity was assessed following intramuscular administration of LNP-formulated reporter mRNAs into the *M. tibialis* of BALB/c mice. Although the Cap 1 analogue containing the flexible diethylene glycol linker produced initial expression levels comparable to those of Cap 1, reporter activity declined more rapidly over time and remained clearly below that achieved with Cap 1-3′-OMe (Figure 4A). Incorporation of the 2′SMe modification improved in vivo expression relative to Cap 1 and yielded activity approaching that of Cap 1-3′-OMe (Figure 4B). Further conformational restriction through the LNA-modified cap analogue increased expression to levels comparable to those obtained with Cap 1 3′-OMe (Figure 4C). Together, these findings support the notion that reduced conformational flexibility and increased structural preorganization of the cap promote efficient translation in vivo. Although CV-Cap5 exhibited favorable properties in structural, biochemical, and cellular assays, its low synthetic yield currently precluded in vivo evaluation and highlights the need for further optimization of the synthetic route.

**Figure 4:**
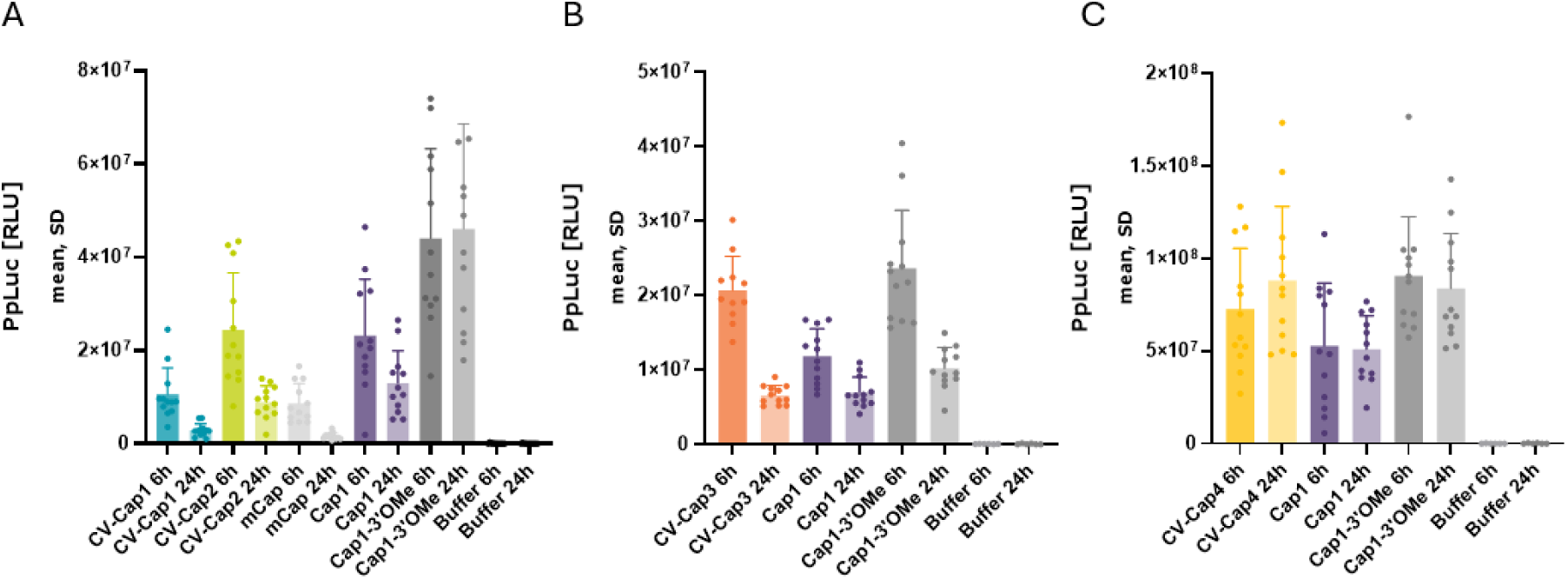
In vivo translation activity of capped m^1^Ψ-modified mRNAs after intramuscular administration in BALB/c mice. PpLuc activity was determined in muscle lysates after intramuscular injection of LNP-formulated mRNAs into the *M. tibialis*. A) ALC-315 LNPs, mRNAs carrying CV-Cap1 versus commercial cap analogues, analysed at 6 h and 24 h after injection. B) SM-102 LNPs, mRNAs carrying CV-Cap3 versus commercial cap analogues, analysed at 6 h and 24 h after injection. C) SM-102 LNPs, mRNAs carrying CV-Cap4 versus commercial cap analogues, analysed at 6 h and 24 h after injection. For panels A-C, data are shown as mean ± standard deviation (n = 12).

## DISCUSSION

In recent years mRNA has emerged as a powerful therapeutic modality for protein replacement therapies and vaccines. Therefore, modulation of factors that impact mRNA-encoded gene expression, such as mRNA stability, immunity and translation efficiency are of high importance. The translation initiation factor eIF4E binds to the 5’ m^7^GpppN mRNA cap structure and is not only essential for cap-dependent mRNA translation, but is implicated in all aspects of mRNA metabolism (18). In this context, modulation of the 5′ cap structure, its recognition by eIF4E, and the resulting effects on gene expression are of particular interest for therapeutic applications of mRNA technology.

In the present study we synthesised different 5’ cap analogues and investigated their binding to eIF4E to as well as their influence on gene expression levels in human cells and mice. Crystal structure analysis of the different eIF4E-cap complexes shows, that recognition of the m^7^G base of the non-canonical cap analogues is preserved in all eIF4E complex structures determined in this work and available in the PDB (Figure 1+2 and Supporting Information). Moreover, an intricate network of water mediated contacts is formed between eIF4E and the triphosphate linking unit (Figure 1 and Supporting Figure 5), the importance has been previously reported for cap recognition by eIF4E (41). Increasing the flexibility of the cap structure by replacing the m^7^G-ribose with a diethylene glycol linker (CV-Cap1, 2), results in perturbation of this water network (Supporting Figure 7), lower binding affinity (Figure 1A) and reporter gene expression in cell lines and mice (Figure 3 and 4). Albeit the N^6^-benzyl moiety of the CV-Cap5 occupies a novel binding pocket and forming a dense interaction network, its binding affinity is comparable to Cap 1 and reporter gene expression levels are partly even below the Cap 1 and Cap 1 3′-OMe (Figure 3). This is likely due to the entropic cost associated with the displacement of water molecules, again emphasising the importance of the water-mediated contacts, and the proximity of the 3′-OMe and ribose in the binding position (Figure 2). Rigidifying the m^7^G ribose by introducing an LNA-modification (CV-Cap4) does not perturb the water network (Supporting Figure 8) and leads not only to higher binding affinity (Figure 1A), but also to prolonged gene expression *in vivo* (Figure 4), albeit reporter gene expression levels seem to be dependent on the cellular context (Figure 3).

In summary, our structural, biochemical, and functional data show a relationship between cap architecture and translational output and provide a framework for the rational optimization of mRNA therapeutics. While increase in conformational flexibility of the cap structure, as introduced by the diethylene glycol linker, resulted in reduced reporter expression, rigidification by 2’SMe and LNA substitution was associated with enhanced translational output. Moreover, preservation of the intricate network of water-mediated contacts is of importance. This suggests that optimization of cap conformation represents an effective strategy for improving the performance of therapeutic mRNAs. The structural observations reported here point to previously unexplored opportunities for cap modification. Future studies may explore whether optimization of interactions within the newly identified binding pocket, together with maintenance of the associated water-mediated contacts, can be exploited to further enhance cap-dependent translation.

## ACKNOWLEDGEMENTS

We thank the beamline scientist of the beamlines David von Stetten, Sihyun Sung, Michael Agthe and Yehudi Bloch (P13 and P14 at DESY) and Philippe Carpentier (MASSIF-3 at ESRF) for their support during data collection. We also thank technical staff members Aline Trapp, Stephan Boshof, Robert Techel and Vanessa Schade for chemical synthesis and Laura Sauer, Franziska Polzer, Luca Hüttenmeister, Mirjam Rauch, Julia Herold, Rabea Haberer and Melina Klinger for biological testing at CureVac.

## AUTHOR CONTRIBUTIONS

S. S.: Conceptualization, Formal analysis, Methodology, Validation, Writing -original draft, review & editing. R.J. S.: Conceptualization, Formal analysis, Visualization, Writing – original draft, review & editing. S. M.: Formal analysis, Methodology. M. T.: Methodology, Formal analysis, Writing - review & editing.

## SUPPLEMENTARY DATA

Supplementary Data are available at NAR online.

## DATA AVAILABILIT

Diffraction data and model coordinates were deposited in the protein data base (PDBe) https://www.ebi.ac.uk/pdbe/, with the accession codes: eIF4E Cap1: PDB code 34AK, eIF4E CV-Cap1: PDB code 34CV, eIF4E CV-Cap2: PDB code 34CO and 34DD, eIF4E CV-Cap4: PDB code 34CY, eIF4E CV-Cap5: PDB code 34DH.

