## Supporting Information for "Structural basis of eIF4E recognition of synthetic mRNA 5′ cap analogues links cap architecture to affinity and biological activity"

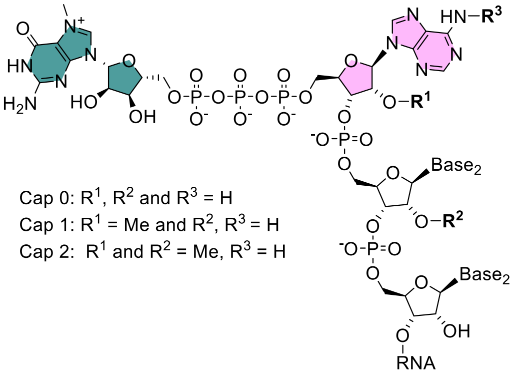

**Supporting Fig. 1.** Eukaryotic mRNA-cap structures.

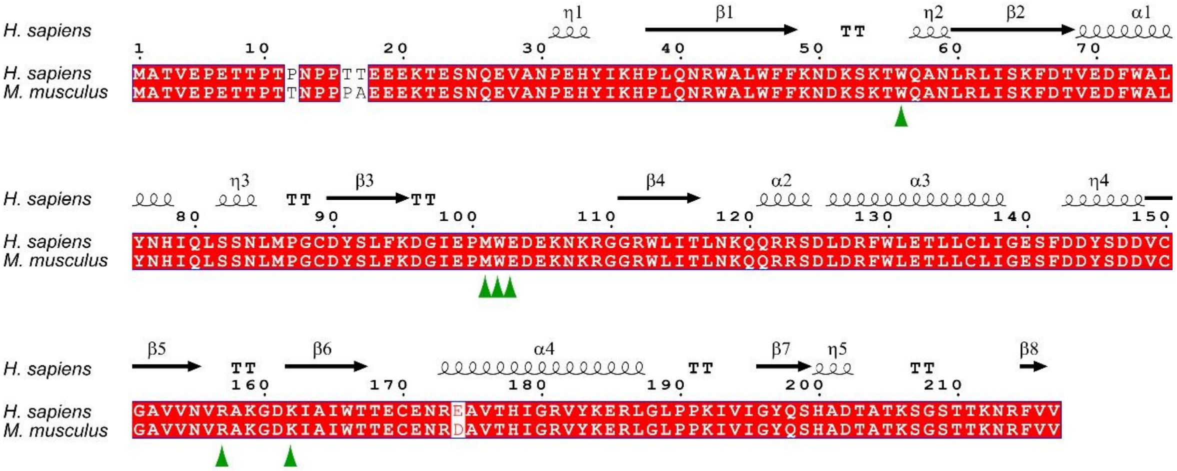

**Supporting Figure 2**. Sequence alignment of human (Uniprot No. P06730) and mouse (Uniprot P63073) eIF4E. Residues forming direct interactions with the bound cap are highlighted with green arrows. The alignment was carried out by ClustalX (1) and annotated with ESPript (2).

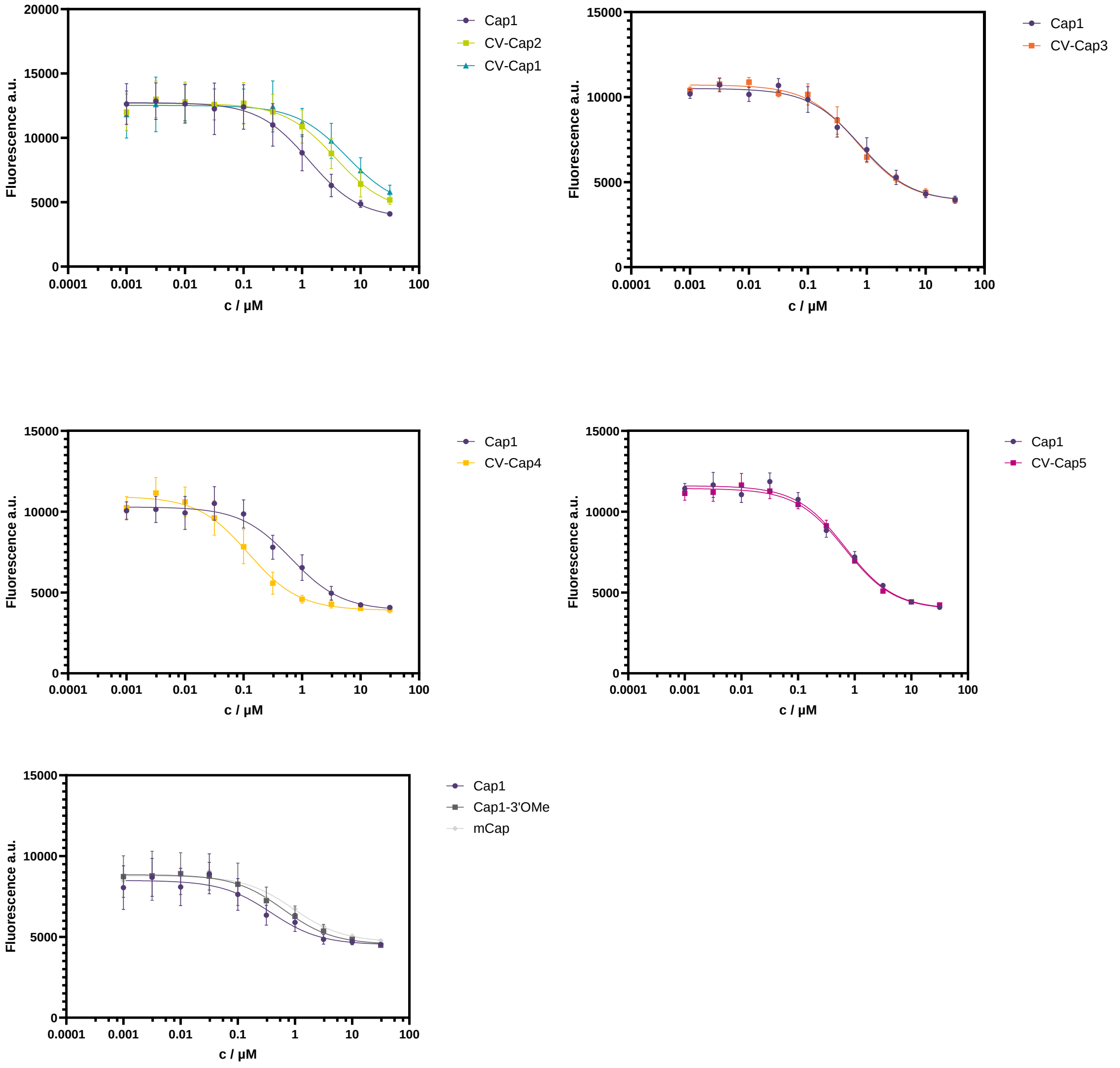

**Supporting Figure 3.** Dose-response curves for CV-Cap analogues and Cap1 reference.

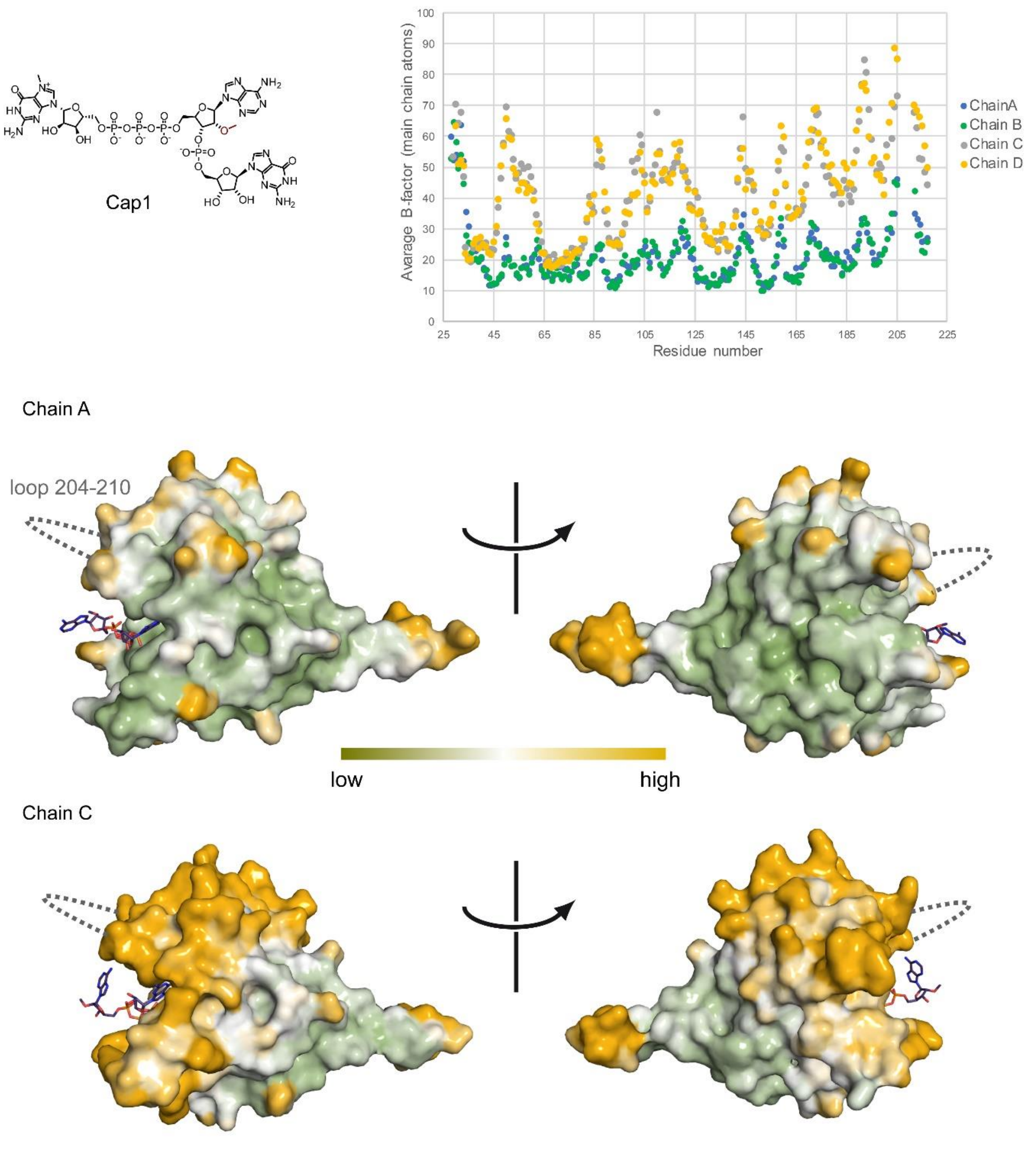

**Supporting Figure 4.** B-factor plots and B-factors mapped onto the surface representation of eIF4E-Cap1 complex, highlighting the different intrinsic flexibility of the four protein complexes in the asymmetric unit of the crystal in the P1 space group.

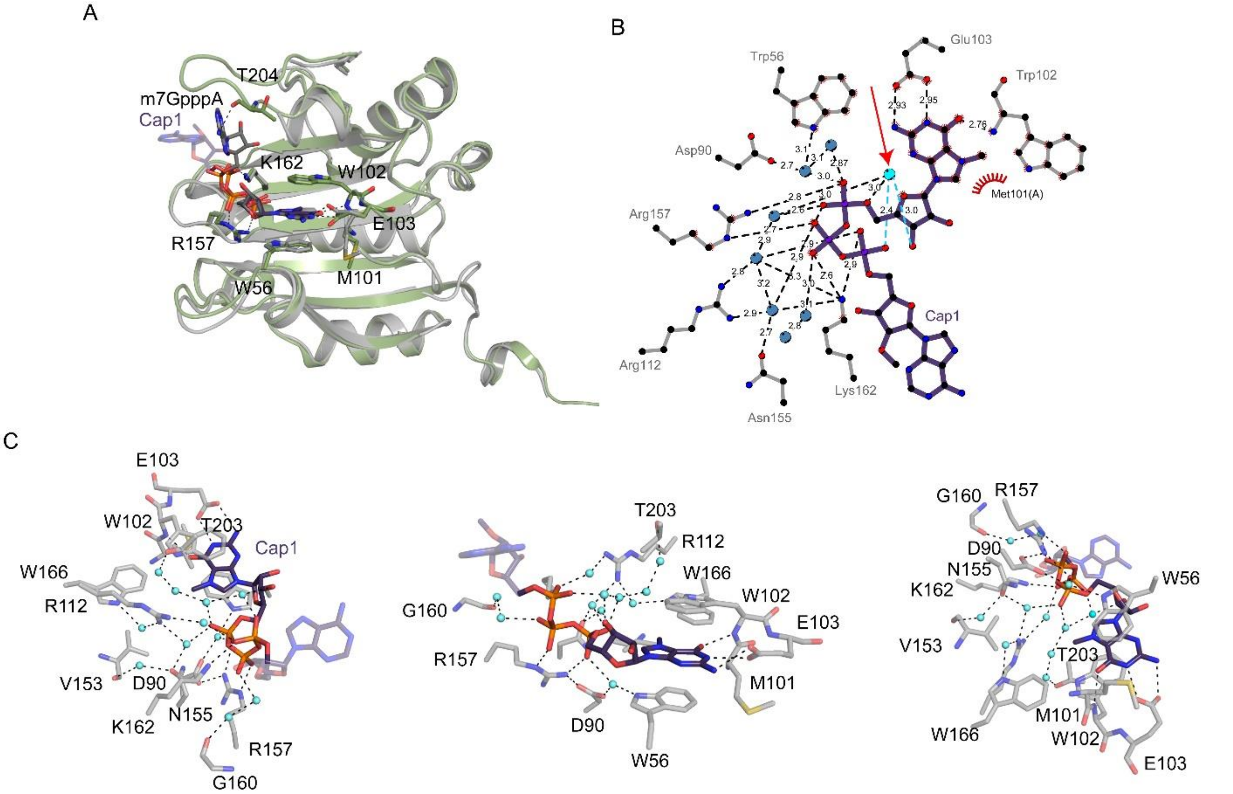

**Supporting Figure 5.** Structure of eIF4E in complex with Cap1. A) Superposition of the eIF4E-Cap1 (light grey ribbon/purple stick model) and m$^7$GpppA (PDB code 1IPB; green) complexes. B) Ligplot diagram (3) highlighting direct and water-mediated contacts. The red arrow indicates the water molecule bridging the m$^7$G 2’OH and the α-phosphate of pppA. C) Intricate water-network of the eIF4E-Cap1 interaction.

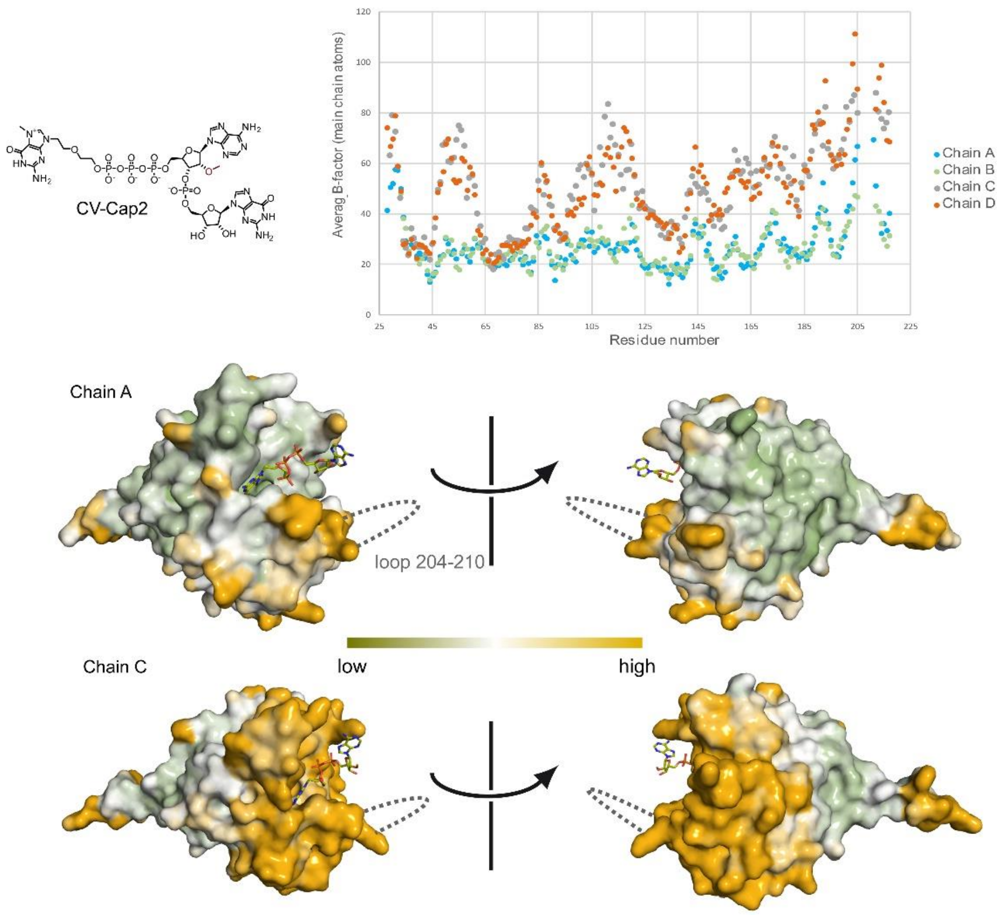

**Supporting Figure 6.** B-factor plots and B-factors mapped onto the surface representation of eIF4E-CV-Cap2 highlighting the different intrinsic flexibility of the four protein complexes in the asymmetric unit of the crystal in the P1 space group.

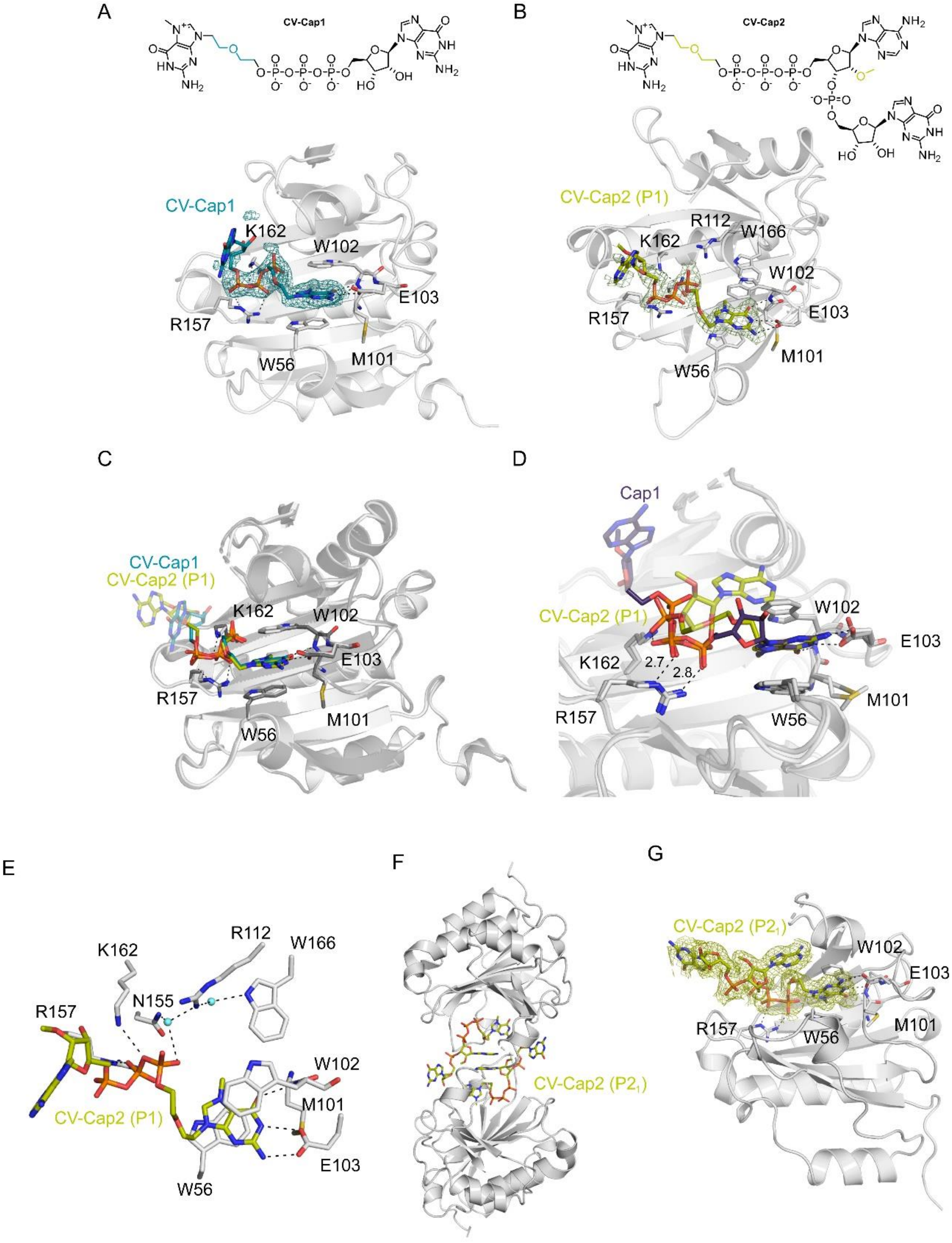

**Supporting Figure 7.** Crystal structures of eIF4E in complex with CV-Cap1 and CV-Cap2 A) CV-Cap1 binding mode and simulated-annealing omit-electron density map (teal mesh) contoured at 2.5σ. B) CV-Cap2 binding mode and simulated annealing omit-electron density map (light green mesh) contoured at 2.5σ, of the eIF4E complex in the P1 symmetry. C) Superposition of CV-Cap1 (light grey/teal) and CV-Cap2 (dark grey/light green, space group P1) complex structures. D)

Superposition of the eIF4E complexes with CV-Cap2 (light grey/light green) and Cap1 (P1 symmetry; dark grey/purple) highlighting the shifted position of the triphosphate bridge and the reduced direct contacts. E) Water network in the CV-Cap2 (P1) complex. F) Crystal packing of the eIF4E- CV-Cap2 complex in the space group $P2_1$. G) The simulated annealing omit-electron density of CV-Cap2, contoured at 2.5$\sigma$, is shown as green mesh in the eIF4E complex in the $P2_1$ symmetry.

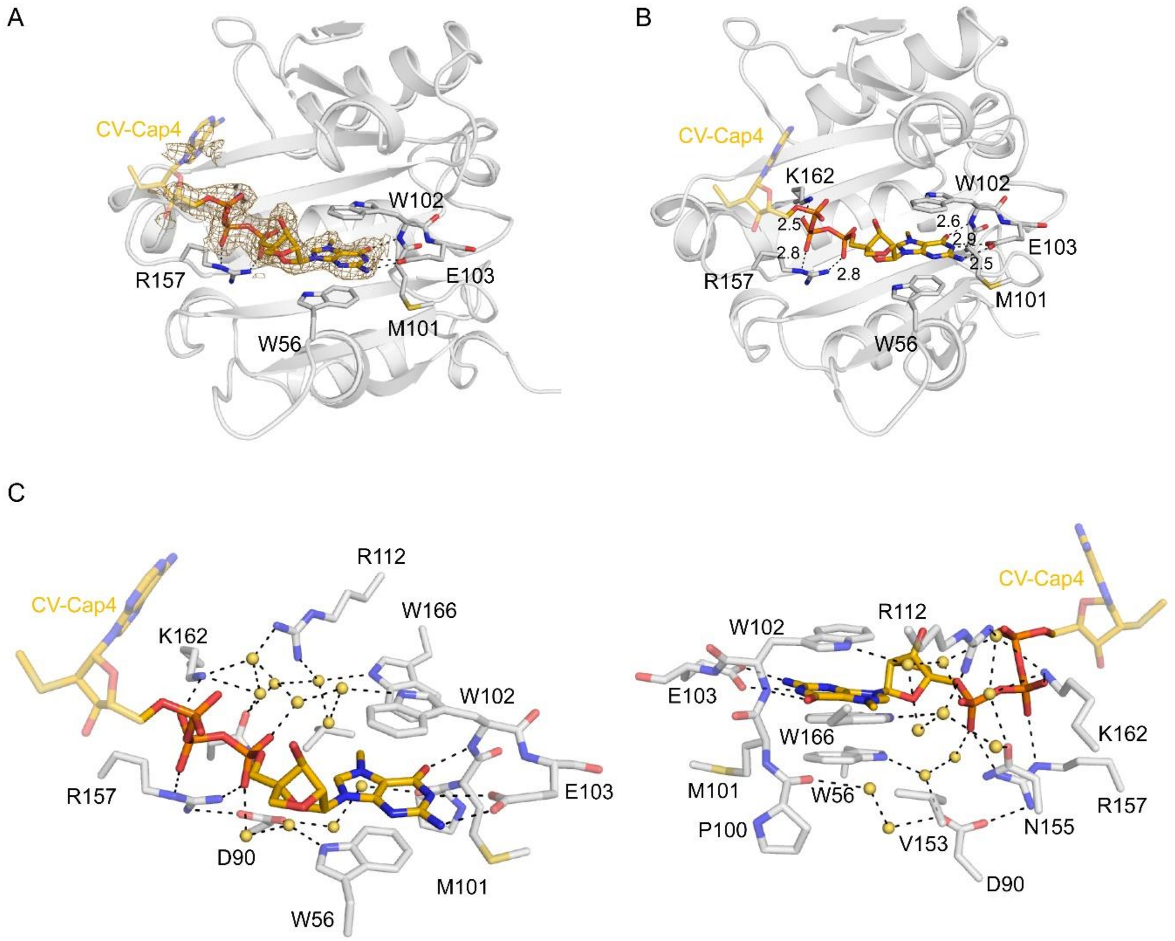

**Supporting Figure 8.** Structure of eIF4E in complex with CV-Cap4. A) Binding mode of CV-Cap4 (golden stick model) to eIF4E (grey ribbon), with residues forming direct interactions shown as stick model. The simulated annealing omit-electron density of CV-Cap4 contoured at 2.5 $\sigma$ is shown as golden mesh. B) Direct Interactions between eIF4E with CV-Cap4, with residues highlighted as stick model. C) Water-network in the complex of eIF4E with CV-Cap4.

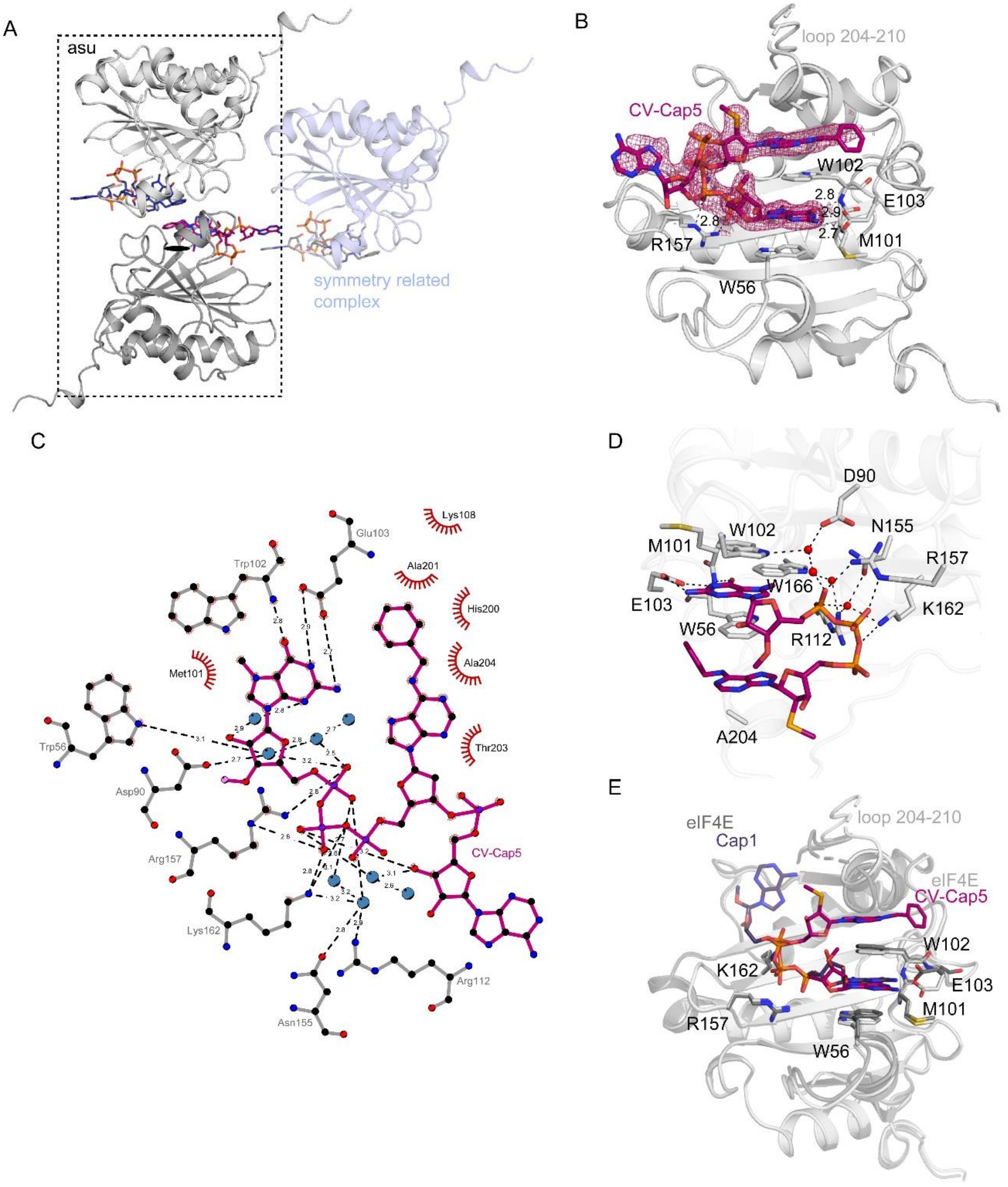

**Supporting Figure 9.** Structure of eIF4E in complex with CV-Cap5. A) Crystal packing interactions. The second transcribed nucleotide forms packing interactions with its symmetry related equivalent. The asymmetric unit (asu) of the crystal lattice is highlighted by the dashed line. B) Binding mode of CV-Cap5 (pink stick model) to eIF4E (grey ribbon), with residues forming direct interactions shown as stick model. The simulated annealing omit-electron density of CV-Cap5 contoured at 2.5 $\sigma$ is shown as pink mesh. C) Ligplot diagram (3) highlighting direct and water-mediated contacts. D) Water-network in the complex of eIF4E with CV-Cap5. E) Superposition of the eIF4E complexes with CV-Cap5 (light grey/pink) and Cap1 (P1 symmetry; dark grey/purple).

**Supporting Table 1.** Amino acid sequences of eIF4E used in this study. The affinity tag is highlighted in bold and the Precission protease cleavage site underlined.

| Name | Sequence |
|---|---|
| *H. sapiens* eIF4E Δ1-27 | **MGSSHHHHHH**SSGLEVLFQG\|PVANPEHYIKHPLQNRWALWFFKNDKSKTWQANLRL<br>ISKFDTVEDFWALYNHIQLSSNLMPGCDYSLFKDGIEPMWEDEKNKRGGRWLITLNK<br>QQRRSDLDRFWLETLLCLIGESFDDYSDDVCGAVVNVRAKGDKIAIWTTECENREAV<br>THIGRVYKERLGLPPKIVIGYQSHADTATKSGSTTKNRFVV |

**Supporting Table 2.** Crystallisation conditions for the different eIF4E-cap analogue complexes and synchrotron beamlines were data have been collected. Please note: for the CV-Cap2 complex structure in the P1 symmetry data collected from two different crystals were combined.

| Cap-analog | Crystallisation condition | Data collection |
|---|---|---|
| Cap1 | 12% w/v PEG 4000 100mM HEPES; pH 7.5 100 mM Sodium acetate | DESY P13 |
| | 8% w/v PEG 10000<br>100 mM Na-acetate, pH 4.6 | ESRF ID30A-3<br>DOI: 10.15151/ESRF-ES-2212959277 |
| CV-Cap1 | 18 % w/v PEG 10,000<br>20 % w/v Glycerol<br>100 mM TRIS; pH 8.5<br>100 mM Sodium chloride | DESY P14 |
| CV-Cap2 P1 | 14% w/v PEG 10,000<br>100 mM Imidazole; pH 8 | DESY P13 |
| | 18% w/v PEG 8,000<br>500 mM LiSO4 | ESRF ID30A-3<br>DOI: 10.15151/ESRF-ES-2212959277 |
| CV-Cap2 $P2_1$ | 18 % w/v PEG 10,000<br>20 % w/v Glycerol<br>100 mM TRIS; pH 8.5<br>100 mM Sodium chloride | ESRF ID30A-3<br>DOI 10.15151/ESRF-ES-2232202474 |
| CV-Cap4 | 14% w/v PEG 10,000<br>100mM Hepes pH, 7.5 | ESRF ID30A-3<br>DOI: 10.15151/ESRF-ES-2232202474 |
| CV-Cap5 | 32% PEG4000<br>800mM LiCl<br>100mM Tris | DESY P14 |

**Supporting Table 3.** X-ray data processing and structure refinement statistics.

| | CV-Cap1 PDB code 34CV | CV-Cap2 (P1) PDB code 34CO | CV-Cap2 ($P2_1$) PDB code 34DD | CV-Cap4 PDB code 34CY | CV-Cap5 PDB code 34DH | Cap1 PDB code 34AK |
|---|---|---|---|---|---|---|
| **Wavelength (Å)** | 0.976 | 0.968 | 0.972 | 0.968 | 0.976 | 0.968 |
| **Resolution range** | 37.0 - 1.75 (1.77 - 1.75) | 48.9 - 2.0 (2.04 - 2.0) | 47.7 - 1.75 (1.77 - 1.75) | 48.9 - 2.0 (2.04 - 2.0) | 37.2 - 2.0 (2.05 - 2.0) | 37.1 - 2.05 (2.09 -2.05) |
| **Space group** | P1 | P1 | $P2_1$ | P1 | $P2_1$ | P1 |
| **Unit cell (a, b, c, α, β, γ)** | 38.1 38.1 146.6 84.4 88.9 77.0 | 38.2 38.2 147.6 84.0 87.4 76.6 | 40.0 128.5 71.6 90 97.3 90 | 38.1 38.0 147.5 83.8 87.8 76.9 | 86.0 38.3 92.104 90 100.46 90 | 38.1 38.1 147.4 84.8 88.8 76.8 |
| **Total reflections** | 587,645 (175) | 198,086 (10076) | 1,150,200 (40,807) | 695,966 (732) | 695,917 (48,099) | 791,977 (1,308) |
| **Unique reflections** | 158,256 (71) | 25,609 (1295) | 157,470 (5,531) | 180,311 (187) | 97,923 (6,764) | 58,310 (165) |
| **Multiplicity** | 3.7 (2.5) | 7.7 (7.8) | 7.3 (7.4) | 3.9 (3.9) | 7.1 (7.1) | 13.6 (7.9) |
| **Completeness (%)** | 93.0 (92.7) | 97.5 (93.3) | 98.7 (98.6) | 92.5 (93.5) | 99.5 (99.7) | 99.6 (99.2) |
| **Mean I/sigma(I)** | 8.4 (0.7) | 5.3 (1.0) | 6.4 (0.5) | 3.4 (0.2) | 4.6 (0.3) | 4.88 (1.64) |
| **Wilson B-factor** | 26.9 | 25.9 | 25.34 | 31.84 | 31.0 | 22.43 |
| **R-merge** | 0.073 (1.125) | 0.307 (1.35) | 0.157 (2.751) | 0.286 (5.711) | 0.314 (5.681) | 0.468 (1.048) |
| **CC1/2** | 0.998 (0.38) | 0.984 (0.54) | 0.998 (0.19) | 0.985 (0.16) | 0.992 (0.15) | 0.976 (0.54) |
| **CC*** | 1 (0.74) | 0.996 (0.84) | 0.999 (0.56) | 0.996 (0.53) | 0.998 (0.52) | 0.994 (0.84) |
| **Reflections used in refinement** | 74,849 (2,555) | 53,258 (2,677) | 71,190 (2,805) | 50,001 (2,730) | 40,416 (2,835) | 50,362 (2,796) |
| **Reflections used for R-free** | 3,856 (133) | 2,663 (134) | 3,558 (141) | 2,501 (136) | 2,021 (142) | 2,505 (141) |
| **R-work** | 0.250 | 0.292 | 0.206 | 0.260 | 0.212 | 0.226 |
| **R-free** | 0.283 | 0.354 | 0.239 | 0.287 | 0.239 | 0.276 |
| **Number of non-hydrogen atoms** | 6,414 | 6,387 | 6,639 | 6,579 | 3,413 | 6,614 |
| **macromolecules** | 6,017 | 6,067 | 6,106 | 6,163 | 3,149 | 6,110 |
| **ligands** | 197 | 196 | 196 | 212 | 122 | 216 |
| **solvent** | 200 | 124 | 337 | 204 | 142 | 288 |
| **Protein residues** | 723 | 731 | 725 | 740 | 374 | 733 |
| **RMS(bonds)** | 0.010 | 0.012 | 0.010 | 0.010 | 0.011 | 0.011 |
| **RMS(angles)** | 1.59 | 1.82 | 1.54 | 1.67 | 1.69 | 1.76 |
| **Ramachandran favored (%)** | 97.7 | 94.8 | 98.3 | 96.4 | 98.1 | 96.8 |
| **Ramachandran allowed (%)** | 2.3 | 5.0 | 1.7 | 3.4 | 1.9 | 3.2 |
| **Ramachandran outliers (%)** | 0 | 0 | 0 | 0 | 0 | 0 |
| **Rotamer outliers (%)** | 2.3 | 2.8 | 1.99 | 3.00 | 3.3 | 2.4 |
| **Clashscore** | 2.72 | 7.14 | 2.69 | 2.90 | 4.9 | 3.5 |
| **Average B-factor** | 44.9 | 41.0 | 31.9 | 40.9 | 38.2 | 34.9 |
| **macromolecules** | 44.0 | 40.0 | 31.3 | 40.1 | 37.5 | 34.0 |
| **ligands** | 80.9 | 80.3 | 40.3 | 76.3 | 44.2 | 68.2 |
| **solvent** | 36.5 | 25.1 | 33.8 | 31.3 | 37.4 | 28.2 |

**Supporting Table 4. Design and sequence of the reporter mRNA used in this study.** Schematic representation of the reporter mRNA comprising the HSD17B4 5′-UTR, PpLuc coding sequence, PSMB3 3′-UTR, histone stem-loop structure and a 100-nt poly(A) tail. The complete nucleotide sequence is provided below.

| Name | Sequence 5’-3’ |
|---|---|
| PpLuc-reporter mRNA | AGGAGAGUCCCGCAGUCGGCGUCCAGCGGCUCUGCUUGUUCGUGUGUGUGUCGUUGCAG<br>GCCUUAUUCAAGCUUACCAUGGAGGACGCCAAGAACAUCAAGAAGGGCCCCGCCCCGUU<br>CUACCCCCUGGAGGACGGGACCGCGGGCGAGCAGCUCCACAAGGCCAUGAAGCGGUACG<br>CCCUGGUGCCCGGGACCAUCGCCUUCACGGACGCCCACAUCGAGGUCGACAUCACCUAC<br>GCGGAGUACUUCGAGAUGAGCGUGCGCCUGGCCGAGGCCAUGAAGCGGUACGGCCUCAA<br>CACCAACCACCGCAUCGUGGUCUGCUCCGAGAACAGCCUGCAGUUCUUCAUGCCCGUGC<br>UGGGGGCCCUCUUCAUCGGCGUGGCGGUCGCCCCGGCCAACGACAUCUACAACGAGCGG<br>GAGCUGCUGAACUCCAUGGGCAUCAGCCAGCCCACCGUGGUGUUCGUCUCCAAGAAGGG<br>GCUCCAGAAGAUCCUGAACGUGCAGAAGAAGCUGCCGAUCAUCCAGAAGAUCAUCAUCA<br>UGGACAGCAAGACGGACUACCAGGGCUUCCAGUCCAUGUAUACCUUCGUGACCAGCCAC<br>CUCCCCCCGGGGUUCAACGAGUACGACUUCGUCCCCGAGUCCUUCGACCGCGACAAGAC<br>CAUCGCCCUGAUCAUGAACAGCUCCGGCAGCACGGGGCUGCCCAAGGGCGUGGCCCUCC<br>CCCACCGGACCGCGUGCGUGCGCUUCUCCCACGCCCGGGACCCGAUCUUCGGCAACCAG<br>AUCAUCCCCGACACCGCCAUCCUGAGCGUCGUGCCCUUCCACCACGGGUUCGGCAUGUU<br>CACCACGCUGGGGUACCUCAUCUGCGGCUUCCGCGUGGUCCUGAUGUACCGGUUCGAGG<br>AGGAGCUGUUCCUCCGCUCCCUGCAGGACUACAAGAUCCAGAGCGCCCUGCUCGUGCCC<br>ACCCUGUUCUCCUUCUUCGCCAAGAGCACCCUGAUCGACAAGUACGACCUCUCCAACCU<br>GCACGAGAUCGCGAGCGGCGGGGCCCCGCUGAGCAAGGAGGUGGGCGAGGCCGUCGCCA<br>AGCGGUUCCACCUCCCCGGGAUCCGCCAGGGCUACGGGCUGACCGAGACGACCUCCGCC<br>AUCCUGAUCACCCCCGAGGGCGACGACAAGCCCGGCGCGGUGGGGAAGGUGGUCCCGUU<br>CUUCGAGGCCAAGGUGGUCGACCUCGACACCGGCAAGACGCUGGGGGUGAACCAGCGGG<br>GCGAGCUGUGCGUGCGCGGGCCCAUGAUCAUGAGCGGCUACGUCAACAACCCCGAGGCC<br>ACCAACGCCCUCAUCGACAAGGACGGCUGGCUGCACUCCGGGGACAUCGCCUACUGGGA<br>CGAGGACGAGCACUUCUUCAUCGUGGACCGGCUCAAGAGCCUCAUCAAGUACAAGGGCU<br>ACCAGGUGGCGCCCGCCGAGCUGGAGUCCAUCCUGCUCCAGCACCCGAACAUCUUCGAC<br>GCCGGGGUCGCCGGCCUGCCCGACGACGACGCGGGGGAGCUGCCCGCCGCCGUGGUGGU<br>CCUCGAGCACGGCAAGACCAUGACCGAGAAGGAGAUCGUGGACUACGUGGCCAGCCAGG<br>UCACGACCGCCAAGAAGCUGCGCGGCGGGGUGGUGUUCGUCGACGAGGUGCCCAAGGGC<br>CUGACCGGGAAGCUGGACGCGCGGAAGAUCCGCGAGAUCCUCAUCAAGGCCAAGAAGGG<br>CGGCAAGAUCGCCGUCUGAGGACUAGUCCCUGUUCCCAGAGCCCACUUUUUUUUCUUUU<br>UUUGAAAUAAAAUAGCCUGUCUUUCAGAUCUCAAAGGCUCUUUUCAGAGCCACCAAAAA<br>AAAAAAAAAAAAAAAAAAAAAAAAAAAAAAAAAAAAAAAAAAAAAAAAAAAAAAAAAAAA<br>AAAAAAAAAAAAAAAAAAAAAAAAAAAAAAAAAAAAA |

**Supporting Table 5.** In vitro transcription conditions used for the synthesis of capped mRNAs. All reactions contained 50 µg/mL linearized plasmid DNA template and 5 mM of the respective cap analog and were incubated at 37°C for 3 h.

| Parameter | CV-Cap1-2 | CV-Cap3 | CV-Cap4 | CV-Cap5 |
|---|---|---|---|---|
| Buffer | 80 mM HEPES/KOH, pH 7.5 | 40 mM Tris-HCl, pH 1.5 | 40 mM BisTris, pH 6.5 | 40 mM Tris-HCl, pH 7.9 |
| $MgCl_2$ | 24 mM | 16 mM | 25 mM | 16 mM |
| Spermidine | 2 mM | 2.12 mM | 2 mM | 2 mM |
| DTT | 40 mM | 10 mM | 10 mM | 10 mM |
| Pyrophosphatase | 5 U/mL (Thermo) | 2 U/mL (NEB) | 2 U/mL (NEB) | 2 U/mL (NEB) |
| RNase inhibitor | 200 U/mL RiboLock (Thermo Fisher) | 1000 U/mL (NEB) | 1000 U/mL (NEB) | 1000 U/mL (NEB) |
| RNA polymerase | T7-Polymerase | T7-Polymerase | T7-Polymerase | T7-Polymerase |
| DMSO | – | – | 5% | 5% |
| GTP | 3.81 mM | 3.81 mM | 3.81 mM | 1.91 mM |
| ATP | 3.18 mM | 3.18 mM | 3.18 mM | 1.59 mM |
| CTP | 4.33 mM | 4.33 mM | 4.33 mM | 2.17 mM |
| N1-methyl-pseudouridine-TP | 2.13 mM | 2.13 mM | 2.13 mM | 1.07 mM |

**Experimental section**

Chemical synthesis of CV-Cap1

2-(2-(2-amino-6-chloro-purin-9-yl)ethoxy)ethan-1-ol (**1**)

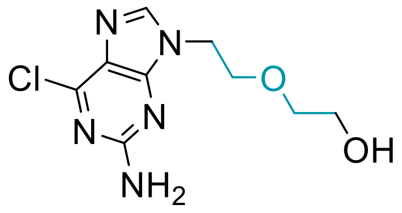

**1**

In a 1 L oven-dried Schlenk flask, triphenylphosphine (37.07 g, 141 mmol, 1.5 eq.), 2-amino-6-chloropurine (23.98 g, 141 mmol, 1.5 eq.), and diethylene glycol (10 g, 8.93 mL, 94.7 mmol, 1.0 eq.) were suspended in dry dioxane (700 mL). Diisopropyl azodicarboxylate (28.5 g, 27.75 mL, 141 mmol, 1 eq.) was added dropwise over 20 min and the solution was stirred for 48 h at ambient temperature. After completion of the reaction, the solvent was evaporated at 0.1 mbar. The residue was dissolved in dichloromethane/methanol (1:1) and adsorbed onto silica (150 g). Purification was performed in three identical runs using flash chromatography on a 330 g silica Ecoflex column (Büchi) and a gradient of 0-15% methanol in ethyl acetate. The alkylated 2-amino-6-chloropurine (**1**) was isolated as an off-white solid (4.53 g, 17.6 mmol) in a yield of 19%.

$^1$H NMR (400 MHz, DMSO-d$_6$) δ 8.11 (s, 1H), 6.90 (s, 2H), 4.61 (t, *J* = 5.0 Hz, 1H), 4.20 (t, *J* = 5.4 Hz, 2H), 3.75 (t, *J* = 5.3 Hz, 2H), 3.50 – 3.38 (m, 4H).

2-(2-(Guanin-9-yl)ethoxy)ethan-1-ol (**2**)

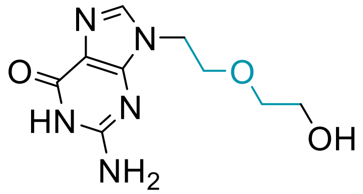

**2**

In a 250 mL flask, compound **1** (5.48 g, 21.27 mmol) was dissolved in a premixed solution of TFA (45 mL) and $H_2O$ (15 mL) and stirred for 72 h at ambient temperature. After completion of the reaction, the solvent was evaporated at 0.1 mbar. The residue was dissolved in methanol and adsorbed onto silica (50 g). Purification was performed by flash chromatography on a 330 g silica Ecoflex column using a gradient of 0-100% methanol in ethyl acetate. The alkylated guanine (**2**) was isolated as a white solid (4.72 g, 19.73 mmol) in a yield of 93%.

$^1$H NMR (400 MHz, DMSO-d$_6$) δ 10.86 (s, 0H), 8.06 (s, 1H), 6.65 (s, 1H), 4.13 (t, *J* = 5.4 Hz, 1H), 3.72 (t, *J* = 5.4 Hz, 1H), 3.50 – 3.39 (m, 2H).

G-C2OC2-p (**3**)

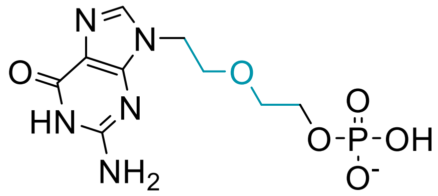

**3**

In a 250 mL flask, compound **2** (3.44 g, 14.4 mmol, 1 eq.) was dissolved in trimethyl phosphate (22.9 mL). The mixture was cooled to 0 °C and phosphor oxychloride (4.4 g, 2.62 mL, 28.8 mmol, 2 eq.) was added dropwise. The reaction was stirred for 3 h at 0 °C. After completion of the reaction, the solvent was evaporated at 0.1 mbar. The residue was diluted with water to a conductivity of < 0.6 mS/cm and purified by anion exchange chromatography using MacroPrep HighQMedia and a linear gradient of 0-50% 1M TEAB in water. The solvent was evaporated at 0.1 mbar, redissolved in 50 mL water and lyophilized. The phosphorylated alkylguanine (**3**) was isolated as a white foam (2.85 g, 8.07 mmol) in a yield of 56 %.

$^{1}H$ NMR (400 MHz, $D_2O$) δ 7.83 (s, 1H), 4.17 (t, J = 4.6 Hz, 2H), 3.89 – 3.84 (m, 2H), 3.83 (t, J = 4.6 Hz, 2H), 3.66 – 3.59 (m, 2H).

$^{31}P$ NMR (162 MHz, $D_2O$) δ 0.43.

m7G-C2OC2-p (**4**)

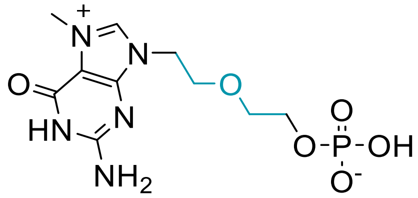

**4**

In a 250 mL flask, compound **3** (3.08 g, 8.72 mmol, 1 eq.) was dissolved in DMSO (87 mL). Methyl iodide (9.9 g, 4.32 mL, 69.7 mmol, 8 eq.) was added and the reaction was stirred for 24 h at 0 °C. After completion of the reaction, the mixture was diluted with water (870 mL) and washed with diethyl ether (3 x 300 mL). The sample was pH adjusted to 7.0 and diluted with water to a conductivity of < 0.6 mS/cm. Purification was performed by anion exchange chromatography using MacroPrep HighQMedia and a linear gradient of 0-50% 1M TEAB in water. The solvent was evaporated at 0.1 mbar, redissolved in 50 mL water and lyophilized. The phosphorylated N7-methyl alkylguanine (**4**) was isolated as a white foam (1.55 g, 4.21 mmol) in a yield of 48%.

$^{1}H$ NMR (300 MHz, $D_2O$) δ 8.95 (s, 1H), 4.43 (t, 2H), 4.12 (s, 3H), 4.03 – 3.91 (m, 4H), 3.80 – 3.71 (m, 2H).

$^{31}P$ NMR (121 MHz, $D_2O$) δ 0.39.

Guanosine diphosphate imidazolide (**5**)

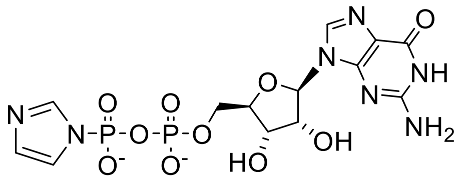

**5**

Commercial guanosine diphosphate sodium salt was purified by anion exchange chromatography to remove monophosphate and triphosphate contaminants. Guanosine diphosphate disodium salt (5 g, 10.3 mmol) was diluted with water to a conductivity of <0.6 mS/cm and purified by anion exchange chromatography using MacroPrep HighQMedia and a linear gradient of 0-100% 1M TEAB in water. The solvent of the product fractions was evaporated at 0.1 mbar, redissolved in 50 mL water and lyophilized, resulting in a guanosine diphosphate triethylammonium salt (5.1 g, 7.9 mmol, 77%) as a white solid.

Guanosine diphosphate triethylammonium salt (1.0 g, 1.59 mmol, 1 eq.) was dissolved in DMF (10 mL). Triethylamine (0.16 g, 0.22 mL, 1.59 mmol, 1 eq.), dipyridyl disulfide (0.70 g, 3.18 mmol, 2 eq.), imidazole (0.54 g, 7.94 mmol, 5 eq.) and triphenylphosphine (0.42 g, 1,59 mmol. 1 eq.) were added. The reaction was stirred for 15 h at ambient temperature. After completion of the reaction, the reaction mixture was precipitated by addition of ice-cold sodium perchlorate in acetone (5.0 g in 50 mL). The precipitate was isolated by centrifugation at 3000 g. The resulting pellet was washed thrice with ice-cold acetone (50 mL) and dried at 0.1 mbar.

Analytical data are in agreement with previously reported data by Kore et. al (4)

CV-Cap1

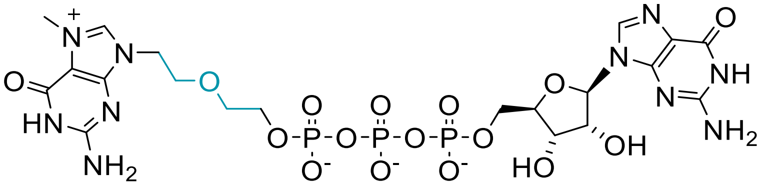

**CV-Cap1**

Guanosine diphosphate 5’-imidazolide (5, 460 mg, 0.86 mmol, 1.5 eq.) was added to a vigorously stirred solution of compound 4 (200 mg, 0.57 mmol, 1.0 eq.) in H2O/MeCN (40 mL, 1/1, v/v). Magnesium chloride (544 mg, 5.71 mmol, 10 eq.) was added to the reaction mixture and stirred at ambient temperature for 24 h. The reaction was quenched by the addition of an EDTA solution (250 mM, 40 mL) and the pH of the resulting solution was adjusted to 7 using sat. aqueous ammonia. The reaction mixture was desalted by RP-HPLC using a Phenomenex Gemini column (250 x 21.2 mm, 5 µm) and a linear gradient of methanol in ammonium acetate (5 mM) from 0-15%. The desalted reaction mixture was purified by ion exchange chromatography on a DNAPac PA200 column (22 x 250 mm) using a linear gradient of 0-75% (Buffer A: 20 mM Tris, pH 9; Buffer B: 20 mM Tris, 33 mM sodium perchlorate). The product fractions were desalted by RP-HPLC using the above-mentioned column with a linear gradient of methanol in ammonium acetate (5 mM) from 0-30%. The solvent was evaporated at 0.1 mbar. The obtained solid was dissolved in 50 mL water and lyophilized. The Product was dissolved in water and concentration was determined by UV measurement using a molar extinction coefficient of ε260=27400 resulting in 0.142 mmol, 29% yield.

Analytical data:

$^1$H NMR (400 MHz, $D_2O$) δ 8.86 (s, 1H), 8.01 (s, 1H), 5.80 (d, *J* = 5.6 Hz, 1H), 4.65 (t, *J* = 5.4 Hz, 1H), 4.49 (t, *J* = 3.8 Hz, 1H), 4.38 – 4.20 (m, 5H), 4.17 – 4.08 (m, 2H), 4.04 (s, 3H), 3.90 (t, 2H), 3.81 – 3.72 (m, 2H).

$^{31}$P NMR (162 MHz, $D_2O$) δ -11.19 (d, *J* = 18.5 Hz), -11.39 (d, *J* = 17.9 Hz), -22.92 (t, *J* = 18.4 Hz).

Chemical synthesis of CV-Cap2

2-Cyanoethylphosphate P-imidazolate (**6**)

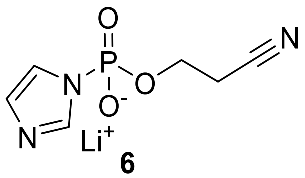

**6**

2-Cyanoethylphosphate-Barium salt monohydrate (20 g, 61.64 mmol) was transformed into triethylammonium salt using DOWEX 50 WX8 triethyl ammonium form (200 g).

2-Cyanoethylphosphate-triethylammonium salt (12.42 g, 49.24 mmol, 1 eq.) was dissolved in dry DMF (150 mL). Carbonyl diimidazole (39.92 g, 246.18 mmol, 5 eq.) were added and the mixture was stirred for 1h. The reaction was quenched by addition of methanol (42.3 mL) and stirred another 30 min. The reaction mixture was precipitated in an acetonitrile (800 mL)/lithium perchlorate (20 g) mixture at 0 °C. The product was isolated by centrifugation of 3000 g. The pellet was washed with acetonitrile at 0 °C (4 x 500 mL) and dried at 0.1 mbar for 2 h. Product was isolated as a brownish powder (8.12 g, 39.22 mmol) in a yield of 80%.

Analytical data are in agreement with previously reported data by Kowalska et.al (5,6)

$^{Bz}$A(2’-OMe)-p-$^{iBu}$G(2,3’-OAc) (**7**)

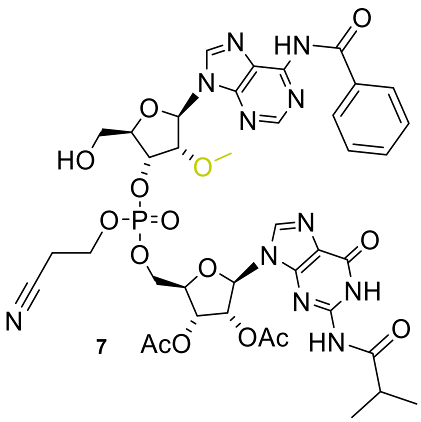

**7**

The two starting materials, 2´,3´-*O*-diacetyl-$N^2$-isobutyryl guanosine (11.2 g, 25.59 mmol, 1.0 eq.) and $N^6$-Benzoyl-2´-*O*-Methyl-5´-*O*-(4,4-dimethoxytrityl)-adenosine 3´-(2-cyanoethyl)-*N,N*-diisopropylphosphoramidite (25 g, 28.15 mmol, 1.1 eq.) were placed in a round-bottom flask and dried under high vacuum overnight. They were then dissolved in acetonitrile under argon atmosphere and tetrazole (114 mL, 51.2 mmol, 2.0 eq., 0.45 M in acetonitrile) was introduced. The reaction mixture was stirred at ambient temperature for 3 h. It was then cooled to 0 °C and a solution of 1 M *tert*-butyl hydroperoxide (*t*BuOOH) in toluene (8.53 mL tBuOOH, 51.2 mmol, 2.0 eq. in 42.65 mL toluene, prepared from 5.0-6.0 M *t*BuOOH in decane) was added dropwise and the reaction mixture was allowed to stir at 0 °C for 1 h. All volatiles were *removed in vacuo* and the remaining residue was dried for 5 min under high vacuum. It was then dissolved in 80% aqueous acetic acid (HOAc) and stirred for 2 h. The solvent was evaporated, and the residue was coevaporated three times with methanol (MeOH). The crude compound was purified by flash chromatography using an Ecoflex Si 330 g column and a gradient of 0-10% Methanol in dichloromethane. The solvent of the product fractions was removed in vacuo and the product was obtained as a foam (20.9 g, 22.29 mmol) in a yield of 87%.

Analytical data:

$^1$H NMR (400 MHz, DMSO-$d_6$) δ 12.13 (s, 1H), 11.59 (d, 1H), 11.26 (d, 1H), 8.77 (m, 2H), 8.29 (m, 1H), 8.05 (d, 2H), 7.65 (t, 1H), 7.55 (t, 1H), 6.20 (m, 1H), 6.14 (m, 1H), 5.85 (m, 1H), 5.55 (m, 1H), 5.40 (m, 1H), 5.21 (m, 1H), 4.83, (q, 1H), 4.48 (m, 3H), 4.33 (m, 1H), 4.28 (m, 2H), 3.64 (m, 2H), 3.42-3.33 (m, 5H), 2.98 (q, 2H), 2.76 (m, 1H), 2.15 (d, 3H), 2.03 (d, 3H), 1.11 (m, 6H).

$^{31}$P-NMR (162 MHz, DMSO-$d_6$), δ -2.38 (d)

pA(2'-OMe)-G (**8**)

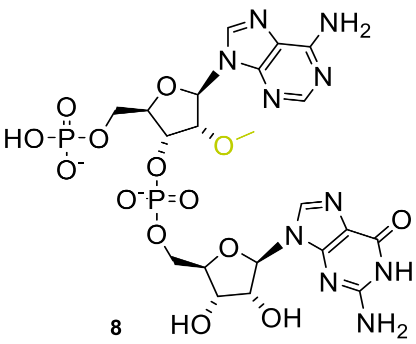

**8**

Compound **7** (4.0 g, 4.27 mmol, 1 eq.) was dissolved in dry acetonitrile under an argon atmosphere. Bis(2-cyanoethyl)-N,N-diisopropylphosphoramidite (2.31 g, 8.53 mmol, 2 eq.) and tetrazole (19.2 mL, 8.53 mmol, 0.45 M in dry acetonitrile, 2 eq.) are added and stirred for 30 min. Oxidizer (0.1 M Iodine in THF/pyridine/water (77:21:2, v/v/v)) was added until red color was observed and stayed for 15 min. A 1:1 mixture of aqueous sodium disulfite solution (5 wt.%) and citric acid solution (5 wt.%) (total volume 600 mL) were added and the crude product was extracted with dichloromethane (3 x 300 mL). Combined organic layers were washed with brine (1 x 300 mL). The solvent was evaporated and the residue was coevaporated with methanol (3 x 50 mL). The protected crude product was dissolved in a 1:1 mixture of methanol/concentrated ammonia (35%) (total volume 150 mL) and stirred for 48 h. The ammonia was evaporated, the residue was coevaporated with methanol (3x 50 mL) and the crude product was dissolved in water.

Conductivity of the sample was measured and adjusted to < 800 µS/cm by further dilution. Purification of diluted sample was performed on a BioRad NGC chromatography system equipped with a HiScale 26/20 column filled with 100 mL MacroPrep High Q resin. Purification was performed using a linear gradient of triethylammonium bicarbonate (1M) in water 0-100%. Product fractions were dried at reduced pressure redissolved in water (50 mL) and freeze dried for 48 h, yielding in 73.2% of an off-white amorph solid.

Analytical data:

$^{1}$H-NMR (400 MHz, DMSO-d$_6$) δ 8.48 (s, 1H), 8.15 (s, 1H), 7.96 (s, 1H), 7.32 (s, 2H), 6.69 (s, 2H), 6.02 (d, 1H), 5.70 (d, 1H), 4.88 (m, 1H), 4.58 (t, 1H), 4.54 (t, 1H), 4.32 (m, 1H), 4.22 (m, 1H), 4.02 (m, 1H), 3.96 (m, 2H), 3.90 (s, 2H), 3.35 (s, 3H).

$^{31}$P-NMR (162 MHz, DMSO-d$_6$), δ -0.84 (s), -1.85 (s)

HR-ESI-MS: [M-H]: 705.1190, calculated 705.1189

ppA(2'-OMe)-G (**9**)

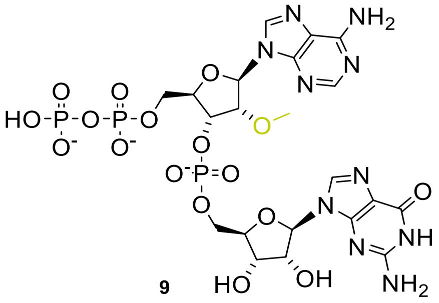

9

Compound **8** (3.19 g, 3.51 mmol, 1 eq.) was dissolved in dry dimethyl sulfoxide. Cyanoethylphosphate-imidazolide lithium salt (**6**) (2.18 g, 10.53 mmol, 3 eq.) and magnesium chloride (3.34 g, 35.1 mmol, 10 eq.) were added. The mixture was stirred for 24 h. After completion of the reaction, DL-dithiothreitol (1.62 g, 10.53 mmol, 3 eq.) and 1,8-diazabicyclo[5.4.0]undec-7-ene (DBU, 66 mL, 438 mmol, 125 eq.) were added and the reaction was stirred for 5 h. DBU and DMSO were removed at 80 °C by high-vacuum distillation. The crude product was dissolved in water and the pH was adjusted to 8.0. The conductivity of the sample was measured and adjusted to < 800 µS/cm by further dilution. The sample was purified by anion exchange chromatography using MacroPrep HighQMedia and a linear gradient of 0-100% 1M TEAB in water. The solvent from the product fractions was evaporated at 0.1 mbar. The residue was redissolved in 50 mL water and lyophilized, resulting in a white powder (2.64 g, 2.41 mmol) in a yield of 69%.

Analytical data:

$^{1}$H NMR (400 MHz, DMSO-d$_6$) δ 8.50 (s, 1H), 8.15 (s, 1H), 7.96 (s, 1H), 7.31 (s, 2H), 6.76 (s, 2H), 6.00 (d, J = 7.2 Hz, 1H), 5.68 (d, J = 6.8 Hz, 1H), 4.95 – 4.86 (m, 1H), 4.67 – 4.60 (m, 2H), 4.32 – 4.27 (m, 1H), 4.26 – 4.22 (m, 1H), 4.10 – 3.94 (m, 4H), 3.89 – 3.81 (m, 1H), 3.51 – 3.47 (m, 2H), 3.31 (s, 3H).

$^{31}$P-NMR (162 MHz, DMSO-d$_6$), δ 1.95 (s), -10.70 (d), -11.50 (d)

HR-ESI-MS: [M+H]: 787.098, calculated 787.099

CV-Cap2

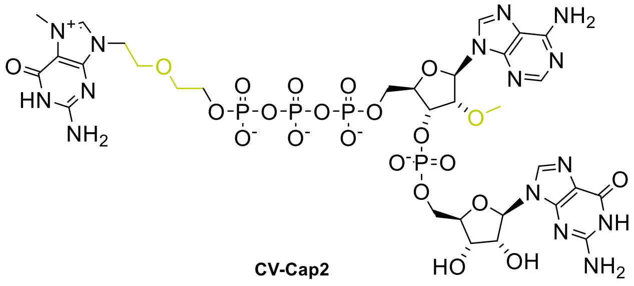

**CV-Cap2**

m7G-$C_2OC_2$-p **4** (100 mg, 0.3 mmol, 1.0 eq) was dissolved in dimethyl sulfoxide (940 µL). 1,1´-Carbonyldiimidazole (487 mg, 3.0 mmol, 10.0 eq.) was added, and the reaction mixture was stirred for 48 h. The reaction was quenched with water (108 mg, 6 mmol, 20.0 eq.) and the reaction mixture was used immediately in the next step.

The above synthesized product m7G-$C_2OC_2$-p-Im (19.2 mg, 174 µL, 0.05 mmol, 1 eq.) was mixed with aqueous N-methylmorpholine (NMM) buffer (2 M, 25 µL, pH 7) and water (50 µL). Manganese chloride (6.3 mg, 0.05 mmol, 1 eq.) and pp-AmG (39.4 mg, 0.05 mmol, 1 eq.) were added and the reaction mixture was stirred for 4 days. The reaction was quenched by the addition of EDTA solution (400 µL, 2.0 eq., 250 mM).
The reaction mixture was then desalted using RP-HPLC on a Phenomenex Gemini C18, 250 x 21.2 mm, 5 µm column using a linear gradient from 0-20% and the solvents A: 5 mM ammonium acetate in water; solvent B: 90% methanol in water. The desalted reaction mixture was purified by RP-HPLC on a Phenomenex Gemini C18, 250x21.2 mm, 5 µm column using a linear gradient from 0-40% 100 mM triethylammonium acetate in water (pH 7, Buffer A) and 90% MeOH in water (Buffer B). The product fractions were finally desalted using RP-HPLC on a Phenomenex Gemini C18, 250x21.2 mm, 5 µM column using a linear gradient from 0-50% of solvent B in solvent A. Solvent A: 5 mM ammonium acetate in water; solvent B: 90% methanol in water. The product fractions were dissolved in 20 mL water and lyophilized resulting in a white powder (40 mg, 36 µmol) in a yield of 72%.
$^1$H NMR (400 MHz, $D_2O$) δ 8.84 (s, 1H), 8.42 (s, 1H), 8.16 (s, 1H), 7.96 (s, 1H), 6.03 (d, J = 5.9 Hz, 1H), 5.84 (d, J = 6.0 Hz, 1H), 4.98 – 4.90 (m, 1H), 4.82 (q, J = 5.6 Hz, 1H), 4.59 – 4.52 (m, 1H), 4.50 (dd, J = 5.2, 3.6 Hz, 1H), 4.44 (t, 1H), 4.39 – 4.33 (m, 1H), 4.35 – 4.24 (m, 3H), 4.23 – 4.18 (m, 2H), 4.13 – 4.06 (m, 2H), 4.03 (s, 3H), 3.82 (t, J = 4.5 Hz, 2H), 3.77 – 3.70 (m, 2H), 3.43 (s, 3H).

$^{31}$P NMR (162 MHz, $D_2O$) δ -0.90 (s, 1P), -11.13 (d, 1P, J = 18.6 Hz), -11.58 (d, 1P, J = 17.0 Hz), -22.89 (t, 1P, J = 18.0 Hz).

HR-ESI-MS: $[C_{31}H_{44}N_{15}O_{22}P_4]^+$: 1102.1733, calculated 1102.1730

Chemical synthesis of CV-Cap3

3′,5′-O-(1,1,3,3-tetraisopropyl-1,3-disiloxanediyl)-9-β-D-arabinofuranosyladenine (**10**)

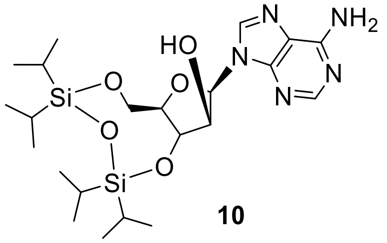

**10**

3′,5′-O-(1,1,3,3-tetraisopropyl-1,3-disiloxanediyl)-9-β-D-arabinofuranosyladenine (50.0 g, 187 mmol, 1.0 eq.) was dissolved in dry pyridine (624 mL). The mixture was repeatedly evacuated *in vacuo* and flushed with nitrogen to remove residual water. The suspension was then cooled to -25 °C using a dry ice/acetone bath. 1,3-dichloro-1,1,3,3-tetraisopropyldisiloxane (59 g, 187 mmol, 1.0 eq.) was added and the reaction mixture was stirred for 16 h while it was allowed to warm to ambient temperature. The turbid reaction mixture was then heated to 50 °C for 30 min. The clear solution was then evaporated, and the residue was partitioned between ethyl acetate (1000 mL) and water (500 mL). The organic layer was washed with 0.2 M HCl, saturated $NaHCO_3$ solution and brine (500 mL each) and then dried over $Na_2SO_4$, filtered and concentrated. The crude product was dissolved in DCM, dry loaded onto hydromatrix (Isolute) and purified by flash chromatography in 5 batches using a linear gradient of 0-60% from DCM to DCM/MeOH (9:1). The product was obtained as white foam (77.2 g, 151.5 mmol) in a yield 81%.

$^1$H NMR (400 MHz, DMSO-$d_6$): δ 8.10 (s, 1H), 8.03 (s, 1H), 7.29 (s br., 2H), 6.20 (d, 1H), 5.79 (d, 1H), 4.59-4.48 (m, 2H), 4.10 (dd, 1H), 3.91 (dd, 1H), 3.79 (m, 1H), 1.19-0.94 (m, 28 H) ppm.

ESI-MS: $[C_{22}H_{40}N_5O_5Si_2]^+$: 510.2, calculated 510.3

3′,5′-O-(1,1,3,3-tetraisopropyl-1,3-disiloxanediyl)-9-β-D-arabinofuranosyl-N6-benzoyl-adenine (**11**)

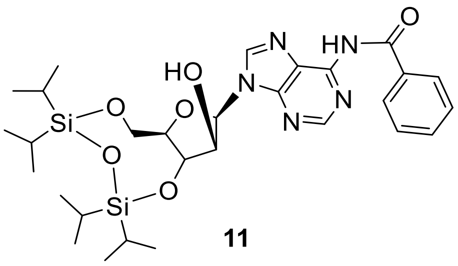

**11**

Compound **10** (71.0 g, 139 mmol, 1.0 eq.) was dissolved in dry pyridine (310 mL) under a nitrogen atmosphere. The reaction mixture was cooled to 0 °C using an ice bath and chlorotrimethylsilane (22.7 g, 209 mmol, 1.5 eq.) was introduced dropwise. The ice bath was removed, and the white suspension was stirred for 1 h at ambient temperature. It was then cooled again to 0 °C and treated dropwise with benzoyl chloride (21.5 g, 153 mmol, 1.1 eq.). The reaction mixture was allowed to warm to ambient temperature and stirred overnight. Additional benzoyl chloride (1.95 g, 13.9 mmol, 0.1 eq.) was added and stirring was continued for an additional hour. The reaction mixture was then cooled again to 0 °C and water (500 mL) was added dropwise while keeping the temperature below 7 °C. Upon complete addition, the reaction mixture was stirred for 1 h at ambient temperature and was then diluted with ethyl acetate (1000 mL). The layers were separated, and the organic phase was washed with water (1000 mL), saturated ammonium chloride solution (1000 mL), $NaHCO_3$ solution (500 mL) and brine (500 mL). It was dried over $Na_2SO_4$, filtered and concentrated to afford a slightly yellow oil which was dissolved in THF (155 mL), MeOH (155 mL) and 7 N ammonia in MeOH (40.1 mL). The reaction mixture was stirred over the weekend, followed by evaporation of the solvent and purification by flash chromatography using a linear gradient from 0-50% (silica 330 g, A: DCM, B: DCM/MeOH 9/1). The product was obtained as a white solid (56.3 g, 91.7 mmol) in a yield 66%.

$^1$H NMR (400 MHz, DMSO-$d_6$): δ 11.22 (s br., 1H), 8.70 (s, 1H), 8.33 (s, 1H), 8.05 (m, 2H), 7.65 (m, 1H), 7.55 (m, 2H), 6.38 (d, 1H), 5.88 (d, 1H), 4.63-4.55 (m, 2H), 4.14 (dd, 1H), 3.94 (dd, 1H), 3.84 (m, 1H), 1.14-1.02 (m, 28 H) ppm.

ESI-MS: $[C_{29}H_{43}N_5O_6Si_2]^+$: 614.2, calculated 614.3

3′,5′-O-(1,1,3,3-tetraisopropyl-1,3-disiloxanediyl)-2’-trifluoracetyl-9-β-D-arabinofuranosyl-N6-benzoyl-adenine (**12**)

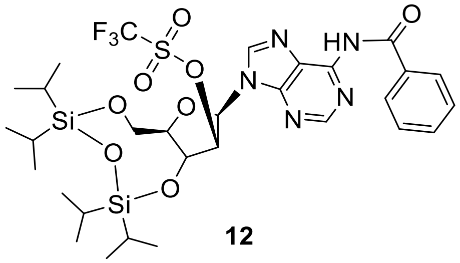

**12**

Compound **11** (69.0 g, 113 mmol, 1.0 eq.) was dissolved in DCM (2.1 L) in an oven-dried flask and cooled to 0 °C. DMAP (41.3 g, 338 mmol, 3.0 eq.) and trifluoromethanesulfonic anhydride (47.7 g, 169 mmol, 1.5 eq.) were added consecutively, followed by stirring for 1.5 h at 0 °C. The reaction mixture was diluted with DCM (1 L), was washed with aq. $NaHCO_3$ solution (5 %, 750 mL) and was dried over $Na_2SO_4$. Filtration and concentration afforded compound **3** as a yellow oil (111.4 g, 45% purity, determined by LC-MS) which was used in the next step without further purification.

ESI-MS: $[C_{30}H_{43}F_3N_5O_8SSi_2]^+$: 746.2, calculated 746.2

3′,5′-O-(1,1,3,3-tetraisopropyl-1,3-disiloxanediyl)-2’-mercaptomethyl-9-β-D-N6-benzoyl-adenine (**13**)

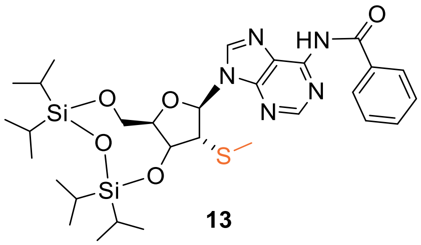

**13**

Sodium thiomethanolate (3.2 g, 45.7 mmol, 1.5 eq.) was dissolved in DMF (250 mL) and compound **12** (47.3 g, 30.4 mmol, 1.0 eq., 48% purity) in dry DMF (100 mL) was introduced. The reaction mixture was stirred at room temperature overnight followed by the addition of more sodium thiomethanolate (3.2 g, 45.7 mmol, 1.5 eq.). Stirring was continued for 4 h after which another portion of sodium thiomethanolate (1.6 g, 22.8 mmol, 0.7 eq.) was added. The reaction was deemed complete by MS after 2 additional hours. It was poured into water (1500 mL) while stirring. Ethyl acetate (2000 mL) was added and the layers were separated using a small amount of brine in order to achieve proper phase separation. The aqueous layer was extracted once with ethyl acetate (1000 mL) and the combined organic phases were washed 4 times with brine (1000 mL each). Drying over $Na_2SO_4$ was followed by filtration, concentration and flash chromatography with a linear gradient of ethyl acetate in n-heptane (5-50%, silica 330 g).

Product fractions were analyzed via ESI-MS and NMR. The product was obtained as a white solid (12.72 g, 19.76 mmol) in a yield of 65% over 2 steps.

$^1$H NMR (400 MHz, DMSO-$d_6$): δ 11.26 (s br., 1H), 8.71 (s, 1H), 8.67 (s, 1H), 8.05 (m, 2H), 7.65 (m, 1H), 7.55 (m, 2H), 6.30 (d, 1H), 5.10 (dd, 1H), 4.28 (dd, 1H), 4.03-3.98 (m, 3H), 2.06 (s, 3H), 1.12-1.01 (m, 28 H) ppm.

ESI-MS: $[C_{30}H_{46}N_5O_5SSi_2]^+$: 644.2, calculated 644.3

2'-mercaptomethyl-9-β-D-N6-benzoyl-adenine (**14**)

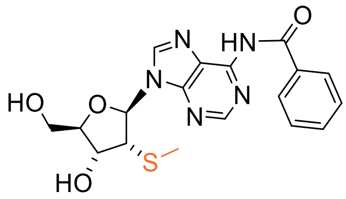

**14**

Tetrabutylammonium fluoride solution (93.2 mL, 93.2 mmol, 1 M in THF, 3.0 eq.) and HOAc (5.6 g, 5.34 mL, 93.2 mmol, 3.0 eq.) were added to a solution of compound **13** (20.0 g, 31.1 mmol, 1.0 eq.) in THF (200 mL) with mechanical stirring. The reaction was deemed complete by MS after 1 h. It was then concentrated under reduced pressure and the residue was redissolved in DCM (200 mL). Extraction with water (200 mL) and brine (200 mL) was followed by drying over $Na_2SO_4$, filtration and concentration. The crude product was purified by flash chromatography in two batches using the gradient program from Table-Ex-6 and Table Ex-7 for each batch respectively (silica 220 g, A: DCM, B: DCM/MeOH 9/1).

**Supporting Table 6.**

| Total time [min] | Flow [mL/min] | %B |
|---|---|---|
| 0 | 100 | 5 |
| 3 | 100 | 5 |
| 46 | 100 | 50 |
| 55 | 100 | 62 |

**Supporting Table 7.**

| Total time [min] | Flow [mL/min] | %B |
|---|---|---|
| 0 | 100 | 5 |
| 3 | 100 | 5 |
| 46 | 100 | 50 |

Product fractions were analyzed via ESI-MS and NMR. The product was obtained as a white foam (9.61 g, 23.9 mmol) in a yield of 77%

$^1$H NMR (400 MHz, DMSO-$d_6$): δ 11.24 (s br., 1H), 8.79 (s, 1H), 8.78 (s, 1H), 8.05 (m, 2H), 7.65 (m, 1H), 7.55 (m, 2H), 6.26 (d, 1H), 5.78 (d, 1H), 5.15 (t, 1H), 4.39 (m, 1H), 4.11 (q, 1H), 4.02 (m, 1H), 3.68 (m, 1H), 3.59 (m, 1H), 1.85 (s, 3H) ppm.

ESI-MS: $[C_{18}H_{20}N_5O_4S]^+$: 402.0, calculated 402.1

5’-Dimethoxytrityl-2’-mercaptomethyl-9-β-D-N6-benzoyl-adenine (**15**)

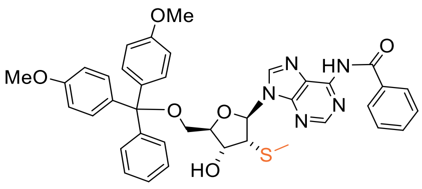

**15**

Compound **14** (19.0 g, 47.3 mmol, 1.0 eq.) was dissolved in pyridine (400 mL). DMAP (2.9 g, 23.7 mmol, 0.5 eq.) and DMTCl (24.0 g, 71.0 mmol, 1.5 eq.) were added consecutively, and the reaction mixture was stirred at ambient temperature overnight. MeOH (15 mL) was added, and the solution was allowed to stir for an additional 30 min. All volatiles were evaporated. The residue was then dissolved in DCM (300 mL) and washed with 5% citric acid (300 mL) and saturated $NaHCO_3$ solution (300 mL). The organic layer was dried over $Na_2SO_4$, was filtered, concentrated, and purified by flash chromatography in three batches using Hydromatrix (Isolute) and a linear gradient from 0-50% of DCM to DCM/MeOH 9/1.

Product fractions were analyzed via ESI-MS and NMR. The product was obtained as a white foam (29.0 g, 41.2 mmol) in a yield of 87%.

$^1$H NMR (400 MHz, DMSO-d$_6$): δ 11.25 (s br., 1H), 8.67 (s, 1H), 8.64 (s, 1H), 8.05 (m, 2H), 7.65 (m, 1H), 7.55 (m, 2H), 7.39 (m, 2H), 7.28-7.22 (m, 7 H), 6.86 (m, 4 H), 6.26 (d, 1H), 5.83 (d, 1H), 4.39 (m, 1H), 4.27 (dd, 1H), 4.14 (m, 1H), 3.73 (s, 3H), 3.72 (s, 3H), 3.32 (dd, 1H), 3.24 (dd, 1H), 1.86 (s, 3H) ppm.

ESI-MS: $[C_{39}H_{38}N_5O_6S]^+$: 704.2, calculated 704.3

5’-Dimethoxytrityl-2’-mercaptomethyl-9-β-D-N6-benzoyl-adenosyl-3’-phosphoramidite (**16**)

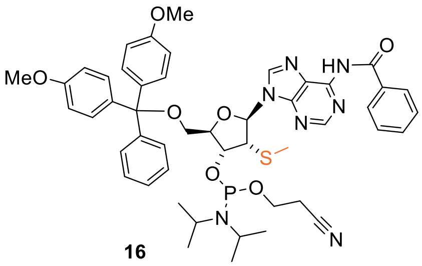

**16**

DIPEA (48.2 g, 65 mL, 373 mmol, 7.5 eq.), 1-methylimidazole (2.0 g, 24.9 mmol, 0.5 eq.) and 2-cyanoethyl *N,N*-diisopropylchlorophosphoramidite (11.8 g, 49.7 mmol, 1.0 eq.) were added consecutively to a solution of compound **15** (35.0 g, 49.7 mmol, 1.0 eq.) in dry DCM (500 mL). After 1.5 h, additional 1-methylimidazole (0.4 g, 5.0 mmol, 0.1 eq.) and 2-cyanoethyl *N,N*-diisopropylchlorophosphoramidit (1.76 g, 7.5 mmol, 0.15 eq.) were added and the reaction mixture was stirred for an additional hour. The solvent was evaporated and the crude product was purified by flash chromatography in three batches using a linear gradient from 30-70% ethyl acetate in n-heptane. The diastereomeric product was obtained as a yellow solid (30.1 g, 33.3 mmol) in a yield of 67%.

$^{1}$H NMR (400 MHz, DMSO-d$_6$): δ 11.26 (s br., 1H), 8.71 (2x s, 1H), 8.61 (2x s, 1H), 8.05 (m, 2H), 7.65 (m, 1H), 7.55 (m, 2H), 7.39 (m, 2H), 7.28-7.18 (m, 7H), 6.84 (m, 4H), 6.26 (t, 1H), 4.66-4.61 + 4.58-4.54 (2x m, 1H), 4.50 (m, 1H), 4.34 + 4.27 (2x m, 1H), 3.92-3.84 + 3.70-3.54 (2x m, 5H), 3.73 + 3.72 + 3.72 (3x s, 6H), 3.38-3.27 (m, 1H), 2.79 + 2.61 (2x t, 2H), 1.91 + 1.83 (2x s, 3H), 1.18-1.04 (m, 12H) ppm.

$^{31}$P NMR (162 MHz, DMSO-d$_6$): δ 151.12, 149.91 (2x s, 1P, diastereomeric signals) ppm.

ESI-MS: $[C_{48}H_{55}N_7O_7PS]^+$: 904.4, Calculated 904.4

A(2′-SMe)-p-G (**17**)

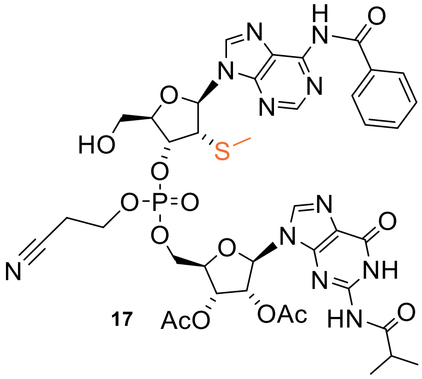

17

Compound **16** (2.5 g, 2.8 mmol, 1.0 eq.) and 2´,3´-*O*-diacetyl-$N^2$-isobutyryl guanosine (1.33 g, 3.0 mmol, 1.1 eq.) were placed in a round-bottom flask and dried under high vacuum overnight. The starting materials were then dissolved in MeCN (12 mL) under an argon atmosphere, and tetrazole (12.4 mL, 5.6 mmol, 2 eq., 0.45 M in acetonitrile) was added. The reaction mixture was stirred at ambient temperature for 2 h. The mixture was cooled to 0 °C and a premixed solution of *t*BuOOH (1.4 mL, 8.4 mmol, 3 eq., 6 M in decane, diluted with 7 mL toluene) was added dropwise and the reaction mixture was stirred for 30 min at 0 °C and for 90 min at ambient temperature. All volatiles were *removed in vacuo* and the remaining residue was dried for 5 min under high vacuum. The product was detritylated by addition of 80% aqueous acetic acid (80 mL, acetic acid/water, 80/20, v/v) for 4 h. The solvent was removed, and the crude product was coevaporated twice with methanol, then dry-loaded onto silica and purified by flash chromatography using the gradient program from Table-Ex-8 (silica 120 g, A: DCM, B: MeOH).

**Supporting Table 8.**

| **Total time [min]** | **Flow [mL/min]** | **%B** |
|---|---|---|
| 0 | 85 | 0 |
| 7 | 85 | 15 |
| 32 | 85 | 15 |
| 37 | 85 | 80 |
| 47 | 85 | 80 |

The diastereomeric product was obtained as a white foam (2.0 g, 2.1 mmol) in a yield of 75%.

$^1$H NMR (400 MHz, DMSO-d$_6$): δ 12.12 (s, 1H), 11.59 (m, 1H), 11.28 (s, 1H), 8.78 (m, 2H), 8.28 (s, 1H), 8.05 (m, 2H), 7.65 (m, 1H), 7.56 (m, 2H), 6.25 (dd, 1H), 6.14 (dd, 1H), 5.84 (t, 1H), 5.56 (m, 1H), 5.33 (s br., 1H), 5.18 (m, 1H), 5.56-4.40 (m, 4H), 4.35-4.26 (m, 3H), 3.64 (m, 2H), 2.97 (m, 2H), 2.78 (m, 1H), 2.15 (s, 3H), 2.03 (s, 3H), 1.97 + 1.93 (2x s, 3H), 1.12 (m, 6H) ppm.

$^{31}$P NMR (162 MHz, DMSO-d$_6$): δ -2.59 (m, 1P, diastereomeric signals) ppm.

HR-ESI-MS: [$C_{39}H_{44}N_{11}O_{14}PS$]$^+$: 954.260, Calculated 954.258

pA(2′-SMe)-p-G (**18**)

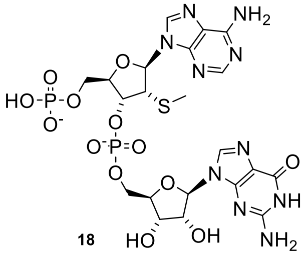

18

Compound **17** (380 mg, 0.40 mmol, 1.0 eq.) was dissolved in acetonitrile (4.0 mL) under an argon atmosphere. Bis(2-cyanoethyl)-N,N-diisopropylphosphoramidite (0.326 g, 1.20 mmol, 3.0 eq.) and tetrazole (2.67 mL, 1.20 mmol, 3 eq., 0.45 M in acetonitrile) were added, and the reaction mixture was stirred for 30 min at ambient temperature. The mixture was cooled to 0 °C and a premixed solution of *t*BuOOH (0.22 mL, 2.4 mmol, 6 eq., 5.5 M in decane, diluted with 2.3 mL toluene) was added dropwise and the reaction mixture was stirred for 60 min at 0 °C. All volatiles were removed *in vacuo*. The remaining residue was coevaporated twice with methanol and was then further dried under high vacuum for 5 min. It was then transferred using methanol (4.0 mL) into a pressure tube and concentrated aqueous ammonia solution (45.0 mL) was added. The reaction mixture was stirred for 16 h at ambient temperature, followed by the addition of a second portion of ammonia (20.0 mL). Stirring was continued for another 24 h followed by the removal of all volatiles and purification by ion-exchange chromatography using a linear gradient from 0-100% TEAB (1M) in water. The product was obtained as a white powder (362 mg, 0.392 mmol) in a yield of 98%.

$^{1}H$ NMR (400 MHz, DMSO-$d_6$): δ 10.89 (s br., 1H), 8.51 (s, 1H), 8.17 (s, 1H), 7.97 (s, 1H), 7.34 (s br., 2H), 6.69 (s br., 2H), 6.14 (d, 1H), 5.71 (d, 1H), 4.87 (dd, 1H), 4.56 (d, 1H), 4.42 (t, 1H), 4.27-4.21 (m, 2H), 4.04-3.87 (m, 5H), 1.81 (s, 3H) ppm.

$^{31}P$ NMR (162 MHz, DMSO-$d_6$): δ 1.43 (s, 1P), -0.21 (s, 1P) ppm.

HR-ESI-MS: $[C_{21}H_{27}N_{10}O_{13}P_2S]^-$: 721.096, Calculated 721.096

CV-Cap3

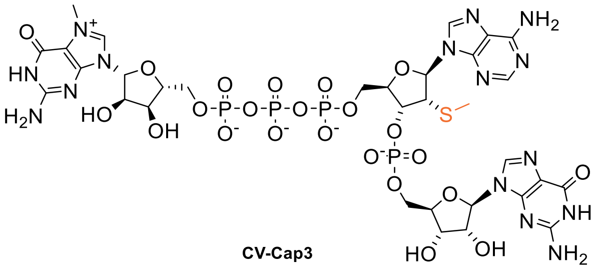

CV-Cap3

N7-Methyl guanosine diphosphate (128 mg, 0.28 mmol, 1.0 eq.) was dissolved in water/DMSO (1.28 mL, 10/90, v/v) and stirred until a clear solution was obtained. EDC·HCl (85.9 mg, 0.45 mmol, 1.6 eq.) and imidazole (57.2 mg, 0.84 mmol, 3.0 eq.) were added consecutively, and the reaction mixture was stirred for 16 h at ambient temperature. A 2 M NMM buffer (pH 7, final concentration 0.2 M), compound **18** (170.1 mg, 0.18 mmol, 0.67 eq.), and $MnCl_2$ (25.7 mg, 0.20 mmol, 0.73 eq.) were added consecutively to the activated nucleoside diphosphate solution. The reaction mixture was stirred for an additional 16 h at ambient temperature. Upon completion of the reaction, a 250 mM EDTA solution (3.2 mL, 0.8 eq.) was added. The reaction mixture was desalted by RP-HPLC using a Phenomenex Gemini C18 column (250 × 21.2 mm, 5 µm) with the gradient program from Table Ex-9 and eluents A (5 mM ammonium acetate in water) and B (90% methanol in water). The desalted material was purified on the same column using 100 mM triethylammonium acetate in water (pH 7, solvent A) and 90% methanol in water (solvent B) with the gradient shown in Table Ex-10. Product-containing fractions were desalted again by RP-HPLC using the gradient program from Table Ex-11.

**Supporting Table 9.**

| Time [min] | %A | %B |
|---|---|---|
| 0 | 100 | 0 |
| 10 | 100 | 0 |
| 20 | 80 | 20 |
| 21 | 5 | 95 |
| 29 | 5 | 95 |
| 30 | 100 | 0 |
| 50 | 100 | 0 |

**Supporting Table 10.**

| Time [min] | %A | %B |
|---|---|---|
| 0 | 100 | 0 |
| 40 | 60 | 40 |
| 50 | 60 | 40 |
| 51 | 100 | 0 |
| 61 | 100 | 0 |

**Supporting Table 11.**

| Time [min] | %A | %B |
|---|---|---|
| 0 | 100 | 0 |
| 10 | 100 | 0 |
| 40 | 50 | 50 |
| 40 | 20 | 80 |
| 51 | 20 | 80 |
| 52 | 100 | 0 |
| 85 | 100 | 0 |

The product was obtained as a white powder (72 mg, 59.4 µmol) in a yield of 33%.

$^{1}$H NMR (400 MHz, $D_2O$): δ 9.04 (s, 1H), 8.43 (s, 1H), 8.15 (s, 1H), 8.01 (s, 1H), 6.10 (d, 1H), 5.86 (m, 2H), 4.94 (m, 1H), 4.82 (m, 1H), 4.55 (m, 3H), 4.45 (t, 1H), 4.42-4.33 (m, 3H), 4.28-4.18 (m, 5H), 4.01 (s, 3H), 3.86 (m, 1H), 1.72 (s, 3H) ppm.

$^{31}$P NMR (162 MHz, $D_2O$): δ -1.00 (s, 1P), -11.39 (d, 1P), -11.51 (d, 1P), -22.86 (t, 1P) ppm.

ESI-MS: $[C_{32}H_{44}N_{15}O_{23}P_4S]^+$: 1162.1397, Calculated 1162.143

Chemical synthesis of CV-Cap4

2′-O,4′-C-methylene-guanosine diphosphate (**19**)

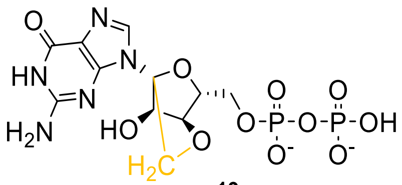

**19**

In a 250 mL flask compound 2′-O,4′-C-methylene guanosine (4.00 g, 13.55 mmol, 1.0 eq.) was dissolved in trimethyl phosphate (30 mL). The mixture was cooled to 0 °C and phosphoryl chloride (2.91 g, 1.73 mL, 18.97 mmol, 1.4 eq.) was added dropwise. The reaction mixture was stirred for 2 h at 0 °C. A premixed imidazole/acetone/water mixture (9.22 g/97.5 mL/1.36 mL) was cooled to 0 °C and added to the reaction mixture. The mixture was stirred for 30 min, during which a precipitate formed. The supernatant was carefully decanted. The solid residue was dissolved in DMF (25.6 mL) and DMSO (12.8 mL). Triethylammonium phosphate (8.09 g, 40.65 mmol, 3.0 eq.)

was added and the mixture was stirred for 16 h. After completion of the reaction, the residue was diluted with water to a conductivity of < 0.6 mS/cm and purified by anion exchange chromatography using MacroPrep HighQ media and a linear gradient of 0-50% 1M TEAB in water. The solvent was removed under reduced pressure, redissolved in water (50 mL) and lyophilized. The 2′-O,4′-C-methylene-guanosine diphosphate (**19**) was isolated as a white foam (5.10 g, 7.76 mmol) in a yield of 57% as triethylammonium salt.

$^{1}$H NMR (400 MHz, $D_2O$) δ 7.87 (s, 1H), 5.73 (s, 1H), 4.58 (s, 1H), 4.47 (s, 1H), 4.35 (dd, *J* = 12.2, 6.5 Hz, 1H), 4.18 (dd, *J* = 12.2, 3.4 Hz, 1H), 4.03 (d, *J* = 8.4 Hz, 1H), 3.90 (d, *J* = 8.4 Hz, 1H).

$^{31}$P NMR (162 MHz, $D_2O$) δ -6.62 (d, *J* = 23.4 Hz), -10.99 (d, *J* = 23.7 Hz).

ESI-MS: [C11H15N5O11P2S] +: 456.0, Calculated 456.0

N7-methyl-2′-O,4′-C-methylene guanosine diphosphate (**20**)

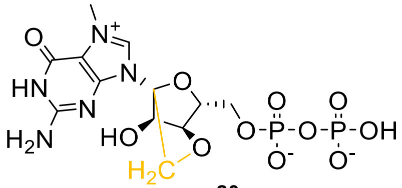

**20**

In a 250 mL flask, compound **19** (5.00 g, 10.98 mmol, 1.0 eq.) was dissolved in water (110 mL). Glacial acetic acid was added until pH 4.0 was reached. Dimethyl sulfate (8.03 g, 6.04 mL, 63.7 mmol, 5.8 eq.) was added dropwise over a period of 1 h. During the reaction, the pH was continuously monitored and maintained between 3.8 and 4.0 by addition of 1 M NaOH. The mixture was stirred for an additional 30 min while maintaining the pH in the range of 3.8–4.0 by addition of 1 M NaOH. After completion of the reaction, the reaction mixture was washed with chloroform (4 × 110 mL). The aqueous phase was adjusted to pH 7.0 and diluted with water to a conductivity of < 0.6 mS/cm. Purification was performed by anion exchange chromatography using MacroPrep HighQ media with a linear gradient of 0–50% 1 M TEAB in water. The solvent was removed under reduced pressure (0.1 mbar), the residue was redissolved in water (50 mL), and lyophilized. N7-methyl-2′-O,4′-C-methylene guanosine diphosphate (**20**) was isolated as a white foam (3.18 g, 4.73 mmol) in 43% yield as the triethylammonium salt.

$^{1}$H NMR (400 MHz, $D_2O$) δ 6.02 (d, *J* = 0.7 Hz, 1H), 4.74 – 4.70 (m, 1H), 4.50 (s, 1H), 4.41 – 4.30 (m, 2H), 4.15 – 4.05 (m, 4H), 4.00 (d, *J* = 8.5 Hz, 1H).

$^{31}$P NMR (162 MHz, $D_2O$) δ -8.61 (d, *J* = 22.3 Hz), -10.86 (d, *J* = 22.6 Hz).

ESI-MS: [C12H18N5O11P2] +: 470.0, Calculated 470.0

CV-Cap4

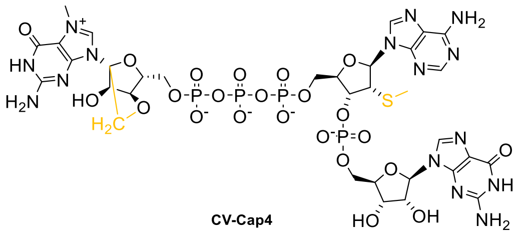

N7-methyl-2′-O,4′-C-methylene guanosine diphosphate di-triethylammonium salt (**20**) (150 mg, 0.22 mmol, 1.5 eq.) was suspended in DMSO (0.70 mL). 1,1′-Carbonyldiimidazole (361 mg, 2.23 mmol, 15.0 eq.) was added, and the reaction mixture was stirred for 2 h. The reaction mixture was quenched by addition of water (0.08 mL, 4.45 mmol, 30.0 eq.). A 2 M N-methylmorpholine buffer (pH 7, final concentration 0.2 M, 0.112 mL), pA(2′-SMe)-p-G di-triethylammonium salt (**18**) (136.9 mg, 0.148 mmol, 1.0 eq.), and DMSO (0.244 mL) were added. The suspension was stirred until large particles had dissolved, and a magnesium(II) chloride solution (3 M, 0.054 mL, 0.163 mmol, 1.1 eq.) was added. The reaction mixture was stirred for 2 days and quenched by addition of a 250 mM EDTA solution (0.71 mL). The reaction mixture was purified by anion exchange chromatography using Q-Sepharose Fast Flow with a linear gradient from solvent A (20% acetonitrile in water) to solvent B (20% acetonitrile in 1.5 M TEAA buffer). The solvent was removed under reduced pressure at 40°C until a final volume of one-tenth of the starting volume was obtained. Ammonium acetate (50 eq.) was added, followed by addition of 10 volumes of ethanol (95%). The mixture was incubated at −20 °C for 16 h and the product was isolated by centrifugation. The resulting solid was dissolved in water and lyophilized, yielding in 30% of a white foam (0.067 mmol).

ESI-MS: $[C_{32}H_{45}N_{15}O_{23}P_4S]^{2+}$: 587.5747, Calculated 587.5737

$^1$H NMR (400 MHz, $D_2O$) δ 8.86 (s, 1H), 8.46 (s, 1H), 8.19 (s, 1H), 7.97 (s, 1H), 6.05 (d, *J* = 8.9 Hz, 1H), 5.78 (d, *J* = 5.7 Hz, 1H), 5.70 (s, 1H), 4.94 – 4.84 (m, 1H), 4.77 (t, *J* = 5.5 Hz, 1H), 4.55 (s, 1H), 4.53 – 4.44 (m, 2H), 4.38 – 4.31 (m, 3H), 4.30 – 4.23 (m, 1H), 4.23 – 4.18 (m, 2H), 4.18 – 4.07 (m, 2H), 4.06 – 4.02 (m, 1H), 4.02 – 4.00 (m, 1H), 3.95 (s, 3H), 3.94 – 3.90 (m, 2H), 3.84 – 3.77 (m, 2H), 1.66 (s, 3H).

$^{31}$P NMR (162 MHz, $D_2O$) δ -1.00, -11.14 (d, *J* = 19.5 Hz), -11.58 (d, *J* = 18.7 Hz), -23.00 (t, *J* = 19.2 Hz).

Chemical synthesis of CV-Cap5

5′-DMT-N6-benzoyl-2′-SMe-3′-OCE phosphoramidite (**21**)

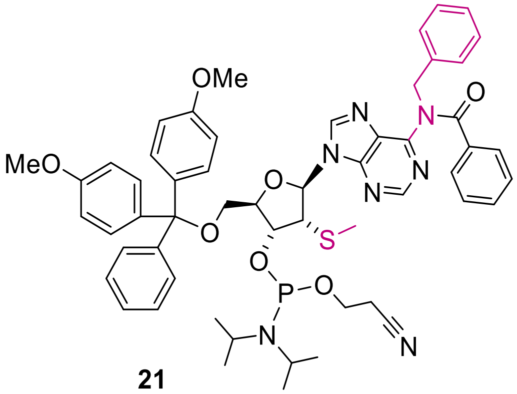

**21**

In an oven-dried 250 mL flask under an inert gas atmosphere, 5′-DMT-N6-benzoyl-2′-SMe-3′-OCE phosphoramidite (2.00 g, 2.21 mmol, 1.0 eq.) was dissolved in DMF (22 mL). Cesium carbonate (959 mg, 2.944 mmol, 1.3 eq.) and benzyl bromide (1090 µL, 4.428 mmol, 2.0 eq.) were added, and the reaction mixture was stirred for 30 min. After completion of the reaction, the mixture was diluted with chloroform (200 mL) and washed with aqueous 0.1% triethylamine (5 × 1 L) and brine (3 × 150 mL). The organic layer was dried over sodium sulfate, and the solvent was removed under reduced pressure. The crude product was purified by flash chromatography using the gradient from

Table Ex-12 (silica, 40 g; solvent A: DCM with 0.1% TEA; solvent B: ethyl acetate).

**Supporting Table 12**

| **Total time [min]** | **Flow [mL/min]** | **%B** |
|---|---|---|
| 0 | 40 | 0 |
| 27.5 | 40 | 20 |
| 28.9 | 40 | 80 |
| 33 | 40 | 80 |
| 38 | 40 | 80 |

The diastereomeric product was obtained as a white foam (1.81 g, 1.82 mmol) in 82% yield.

ESI-MS: $[C_{55}H_{61}N_7O_7PS]^+$: 994.4, Calculated 994.7

$^{Bz}$A(2'-SMe)-p-$^{iBu}$G(2,3'-OAc) (**22**)

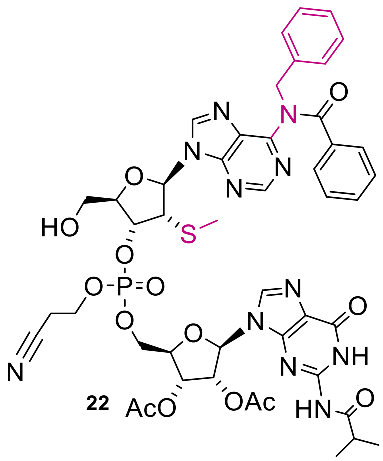

**22**

Compound **21** (2.58 g, 2.59 mmol, 1.0 eq.) and 2′,3′-O-diacetyl-N2-isobutyryl guanosine (1.25 g, 2.85 mmol, 1.1 eq.) were placed in a round-bottom flask and dried under high vacuum overnight. The starting materials were then dissolved in MeCN (14.3 mL) under an argon atmosphere, and tetrazole (11.52 mL, 5.18 mmol, 2.0 eq., 0.45 M in acetonitrile) was added. The reaction mixture was stirred at ambient temperature for 90 min. The mixture was cooled to 0 °C, and a premixed solution of t-BuOOH (1.41 mL, 8.4 mmol, 3.0 eq., 6 M in decane, diluted with 3.59 mL toluene) was added dropwise. The reaction mixture was stirred for 2 h at 0 °C and for 90 min at ambient temperature. All volatiles were removed in vacuo, and the residue was dried under high vacuum for 5 min. The product was detritylated by addition of 80% aqueous acetic acid (94 mL, acetic acid/water, 75.2/18.8, v/v) for 4 h. The solvent was removed, and the crude product was coevaporated twice with methanol. The product was dissolved in ethyl acetate and purified by flash chromatography using the gradient from Table Ex-13 (silica, 40 g; solvent A: ethyl acetate; solvent B: methanol).

**Supporting Table 13**

| **Total time [min]** | **Flow [mL/min]** | **%B** |
|---|---|---|
| 0 | 85 | 0 |
| 30 | 85 | 15 |
| 31 | 85 | 80 |
| 36 | 85 | 80 |

The diastereomeric product was obtained as a white foam (1.82 g, 1.73 mmol) in a yield of 67%.

$^{31}$P NMR (162 MHz, $d_6$-DMSO): δ -3.21 and 3.58 (m, 1P, diastereomeric signals) ppm.

ESI-MS: $[C_{46}H_{50}N_{11}O_{14}PS]^+$: 1044.5, Calculated 1044.3

p-N6-Benzyl-A(2′-SMe)-p-G (**23**)

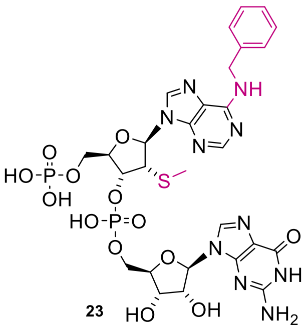

23

Compound **22** (1.80 g, 1.73 mmol, 1.0 eq.) was dissolved in acetonitrile (18 mL) under an argon atmosphere. Bis(2-cyanoethyl)-N,N-diisopropylphosphoramidite (0.937 g, 3.45 mmol, 2.0 eq.) and tetrazole (7.67 mL, 3.45 mmol, 2 eq., 0.45 M in acetonitrile) were added, and the reaction mixture was stirred for 2.5 h at ambient temperature.

The mixture was cooled to 0 °C and a premixed solution of *t*BuOOH (0.58 mL, 3.45 mmol, 2.0 eq., 6 M in decane, diluted with 2.9 mL toluene) was added dropwise and the reaction mixture was stirred for 2.5 h. All volatiles were removed *in vacuo*. The remaining residue was coevaporated twice with methanol and was then further dried under high vacuum for 5 min. It was then transferred using methanol (20.0 mL) into a pressure tube and concentrated aqueous ammonia solution (20.0 mL) was added. The reaction mixture was stirred for 24 h at ambient temperature, followed by 17 h at 40 °C. An additional portion of ammonia (10 mL) was added and the mixture was stirred at 50 °C for 6 h. All volatiles were evaporated and purification by ion-exchange chromatography was performed using a linear gradient from 0-100% TEAB (1 M) in water. The product was obtained as a white powder (1180 mg, 1.108 mmol) in a yield of 64%.

HR-ESI-MS: $[C_{28}H_{35}N_{10}O_{13}P_2S]^+$: 813.155, calculated 813.157

1H NMR (400 MHz, D2O) δ 8.61 (s, 1H), 8.18 (s, 1H), 8.04 (s, 1H), 7.39 – 7.21 (m, 5H), 6.22 (d, J = 9.1 Hz, 1H), 5.88 (d, J = 5.9 Hz, 1H), 4.96 (dd, J = 7.8, 5.0 Hz, 1H), 4.87 (dd, J = 6.0, 5.2 Hz, 1H), 4.56 (dd, J = 5.2, 3.5 Hz, 1H), 4.54 – 4.49 (m, 1H), 4.34 (dq, J = 3.6, 1.8 Hz, 1H), 4.22 (t, J = 3.7 Hz, 2H), 4.04 (ddd, J = 9.4, 5.0, 2.4 Hz, 1H), 3.96 (tt, J = 11.4, 3.7 Hz, 2H), 1.75 (s, 3H)

$^{31}P$ NMR (162 MHz, $d_6$-$D_2O$): δ 3.59 (s, 1P), -0.90 (s, 1P) ppm.

CV-Cap5

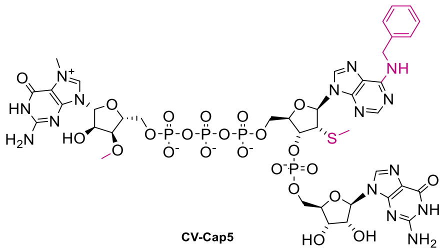

CV-Cap5

N7-methyl-3’-O-methylguanosine diphosphate (258 mg, 0.45 mmol, 3.0 eq.) was suspended in DMSO (1.30 mL). 1,1′-Carbonyldiimidazole (243 mg, 1.5 mmol, 10.0 eq.) was added, and the reaction mixture was stirred for 1 h. The reaction was quenched by addition of water (55 µL, 3.00 mmol, 20.0 eq.). A 2 M N-methylmorpholine buffer (pH 7, final concentration 0.2 M, 0.175 mL) and p-N6-benzyl-A(2′-SMe)-p-G di-triethylammonium salt (**23**) (159.8 mg, 0.15 mmol, 1.0 eq.) were then added. The suspension was stirred until the larger particles had dissolved, after which magnesium(II) chloride (0.016 mg, 0.165 mmol, 1.1 eq.) was added. The reaction mixture was stirred for 3 days and then quenched by addition of 250 mM EDTA solution (0.79 mL).

The reaction mixture was first desalted by RP-HPLC on a Phenomenex Gemini C18 column (250 × 21.2 mm, 5 µm) using the gradient program from Table Ex-14 with solvent A (5 mM ammonium acetate in water) and solvent B (90% methanol in water). The desalted material was then purified by RP-HPLC on the same column using 100 mM triethylammonium acetate in water (pH 7, buffer A) and 90% MeOH in water (buffer B) according to the gradient shown in Table Ex-15. Product-containing fractions were subsequently desalted by RP-HPLC on the same column using a gradient from 0–95% solvent B in solvent A, with solvent A consisting of 5 mM ammonium acetate in water and solvent B of 90% methanol in water, following the gradient shown in Table Ex-16.

**Supporting Table 14.**

| Time [min] | %A | %B |
|---|---|---|
| 0 | 100 | 0 |
| 10 | 100 | 0 |
| 20 | 80 | 20 |
| 21 | 5 | 95 |
| 29 | 5 | 95 |
| 30 | 100 | 0 |
| 58 | 100 | 0 |

**Supporting Table 15.**

| Time [min] | %A | %B |
|---|---|---|
| 0 | 100 | 0 |
| 15 | 70 | 30 |
| 45 | 40 | 60 |
| 46 | 20 | 80 |
| 51 | 20 | 80 |
| 52 | 100 | 0 |
| 74 | 100 | 0 |

**Supporting Table 16.**

| Time [min] | %A | %B |
|---|---|---|
| 0 | 100 | 0 |
| 10 | 100 | 0 |
| 20 | 60 | 40 |
| 45 | 40 | 60 |
| 46 | 20 | 80 |
| 51 | 20 | 80 |
| 52 | 100 | 0 |
| 71 | 100 | 0 |

The resulting solid was dissolved in water and lyophilized, yielding in 15 % of a white foam (0.023 mmol).

HR-ESI-MS: $[[C_{40}H_{52}N_{15}O_{23}P_4S]^+$: 1266.20, calculated 1266.20

$^1$H NMR (400 MHz, $D_2O$) δ 9.03 (s, 1H), 8.47 (s, 1H), 8.17 (s, 1H), 8.09 (s, 1H), 7.24 (s, 5H), 6.10 (d, *J* = 9.0 Hz, 1H), 5.87 (d, *J* = 5.5 Hz, 1H), 5.82 (d, *J* = 4.1 Hz, 1H), 4.95 (t, *J* = 5.3 Hz, 1H), 4.84 – 4.80 (m, 1H), 4.69 (t, *J* = 4.5 Hz, 1H), 4.62 – 4.52 (m, 2H), 4.44 – 4.34 (m, 4H), 4.26 (s, 3H), 4.25 – 4.14 (m, 1H), 4.09 (t, *J* = 4.8 Hz, 1H), 4.00 (s, 3H), 3.95 – 3.84 (m, 1H), 3.44 (s, 3H), 1.72 (s, 3H).

$^{31}$P NMR (162 MHz, $D_2O$) δ -0.94 (s, 1P), -11.45 (d, *J* = 13.4 Hz, 1P), -11.56 (d, *J* = 13.8 Hz, 1P), -22.84 (t, *J* = 18.5 Hz, 1P).

**NMR-Spectra**

2-(2-(2-amino-6-chloro-purin-9-yl)ethoxy)ethan-1-ol (**1**)

$^1$H-NMR

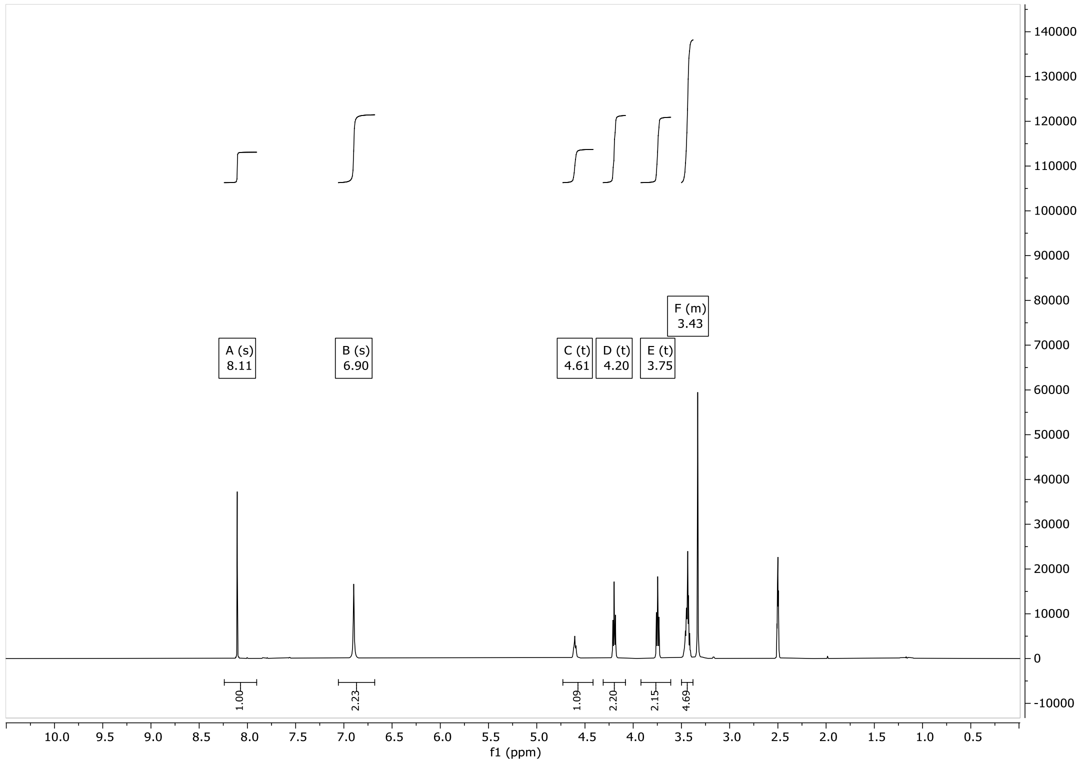

2-(2-(Guanin-9-yl)ethoxy)ethan-1-ol (**2**)

[1]H-NMR

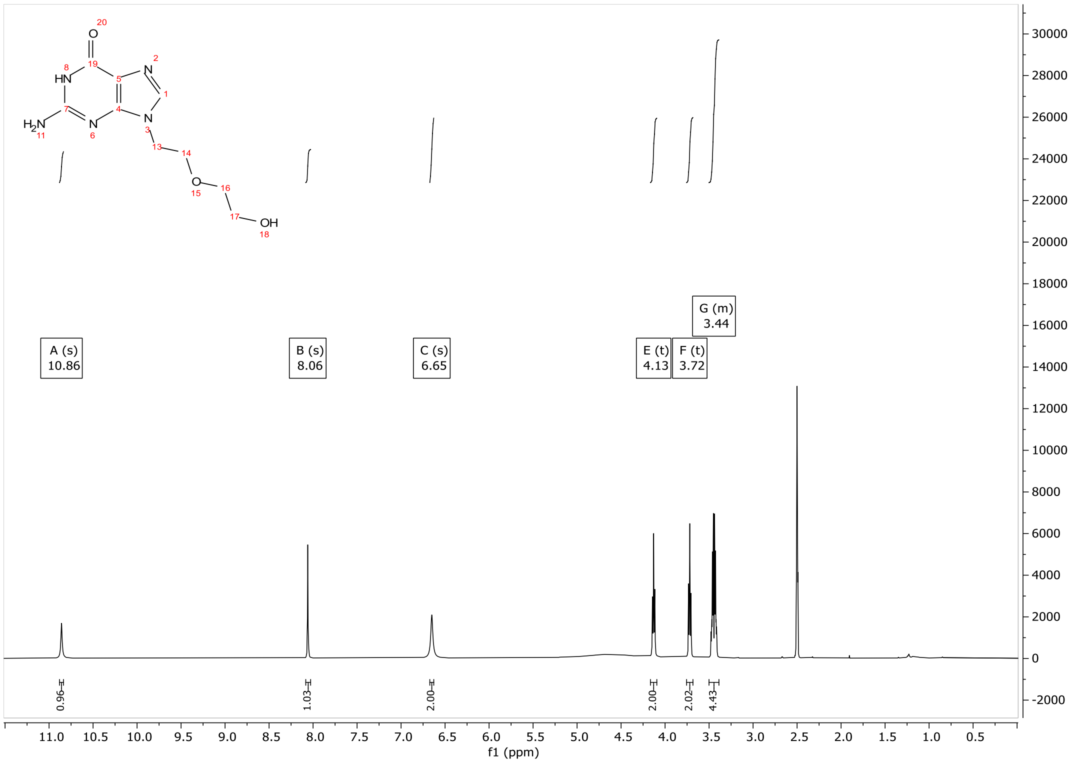

G-C2OC2-p (**3**)

$^1$H-NMR

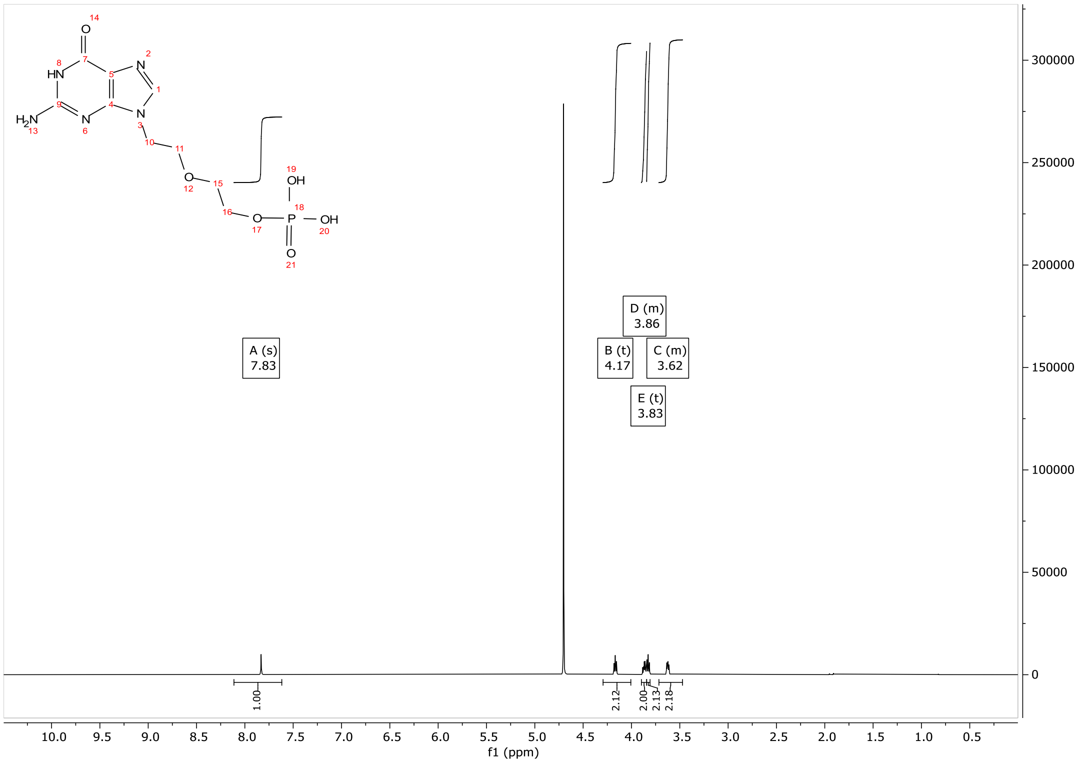

$^{31}$P-NMR

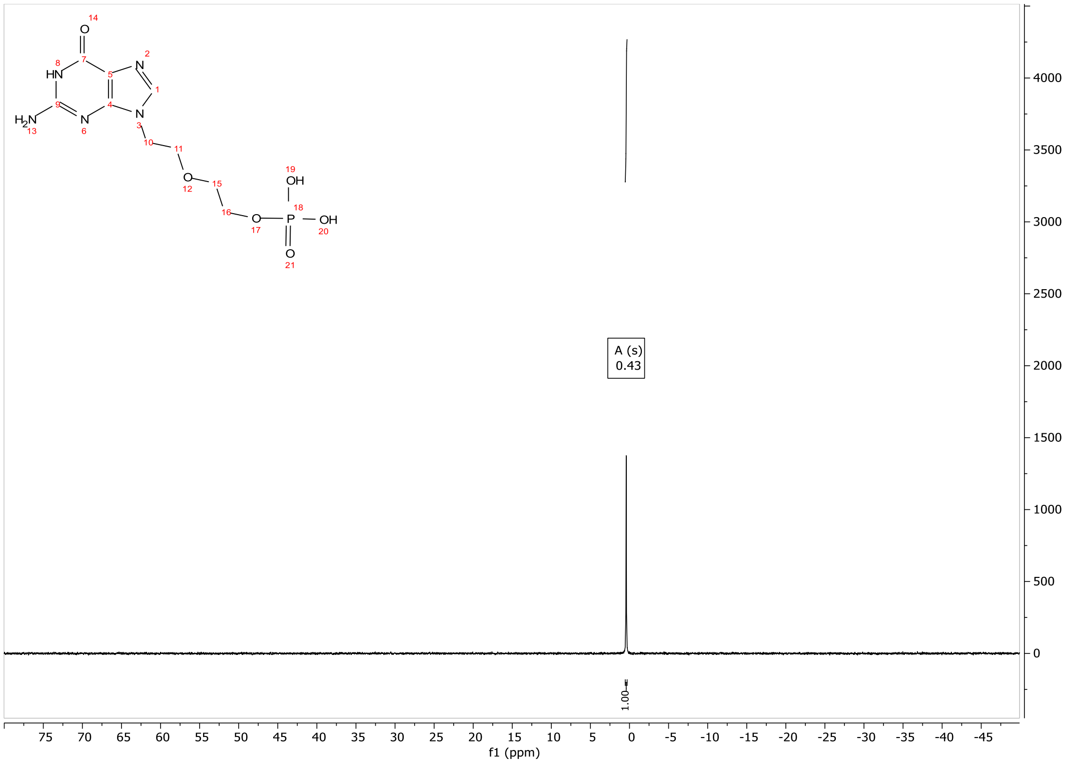

m7G-C2OC2-p (**4**)

$^{1}$H-NMR

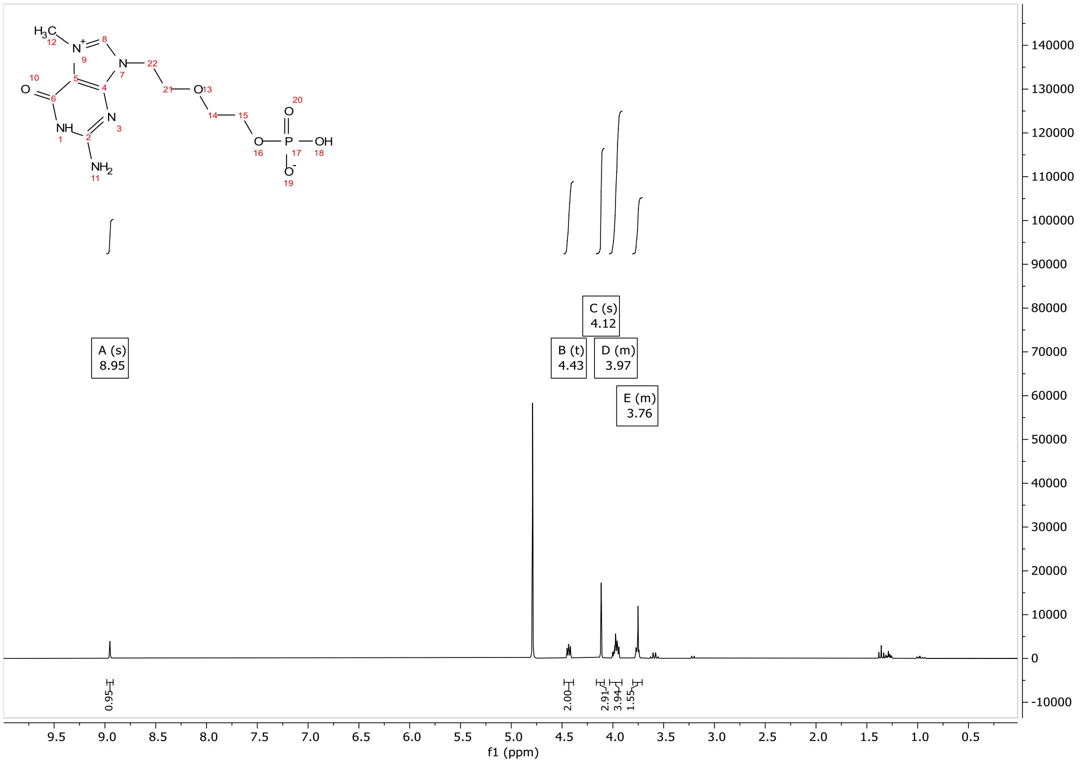

$^{31}$P-NMR

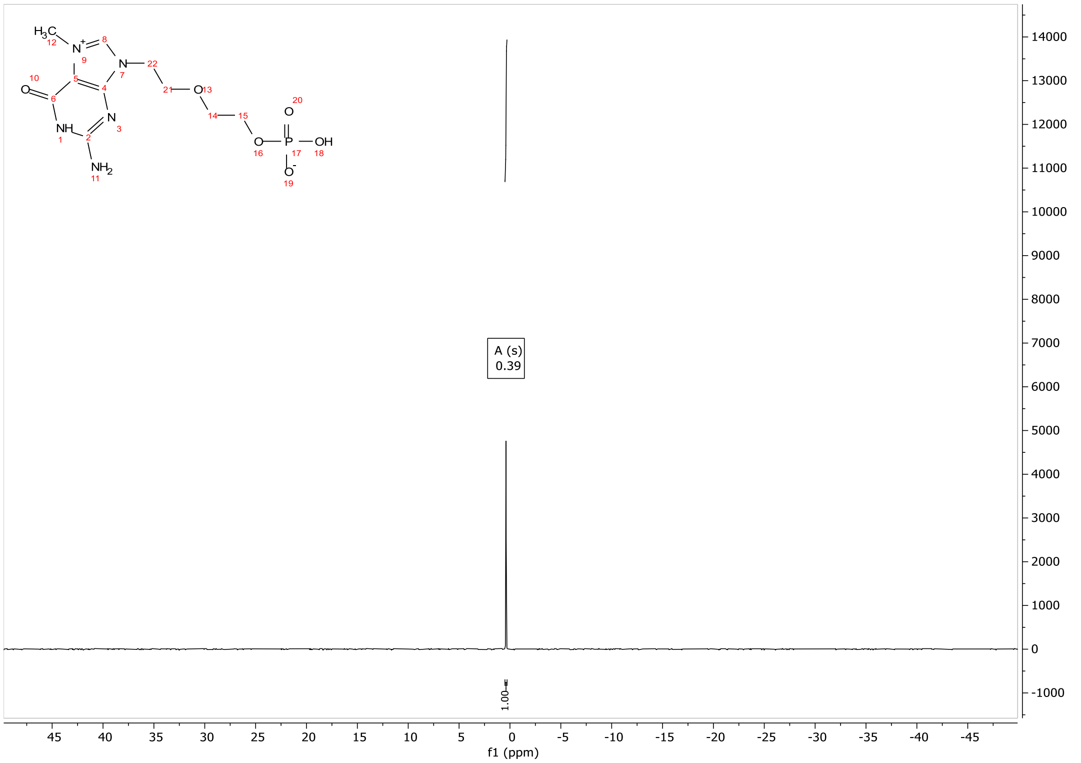

CV-Cap 1

$^{1}$H-NMR

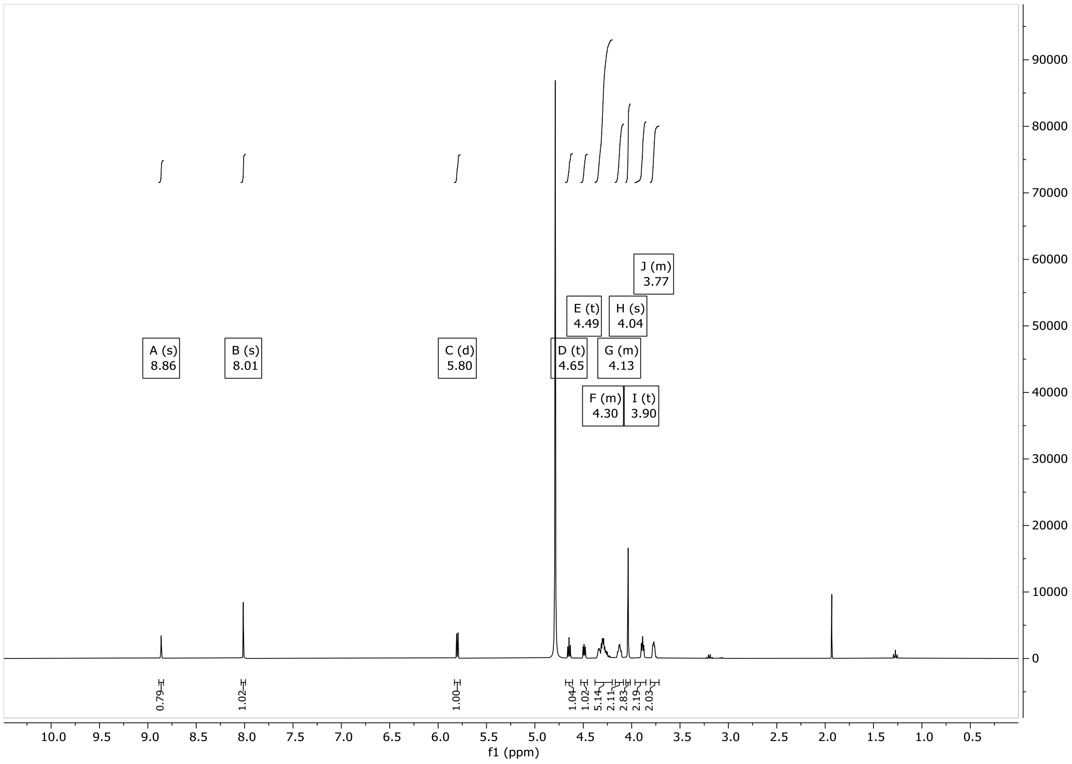

$^{31}$P-NMR

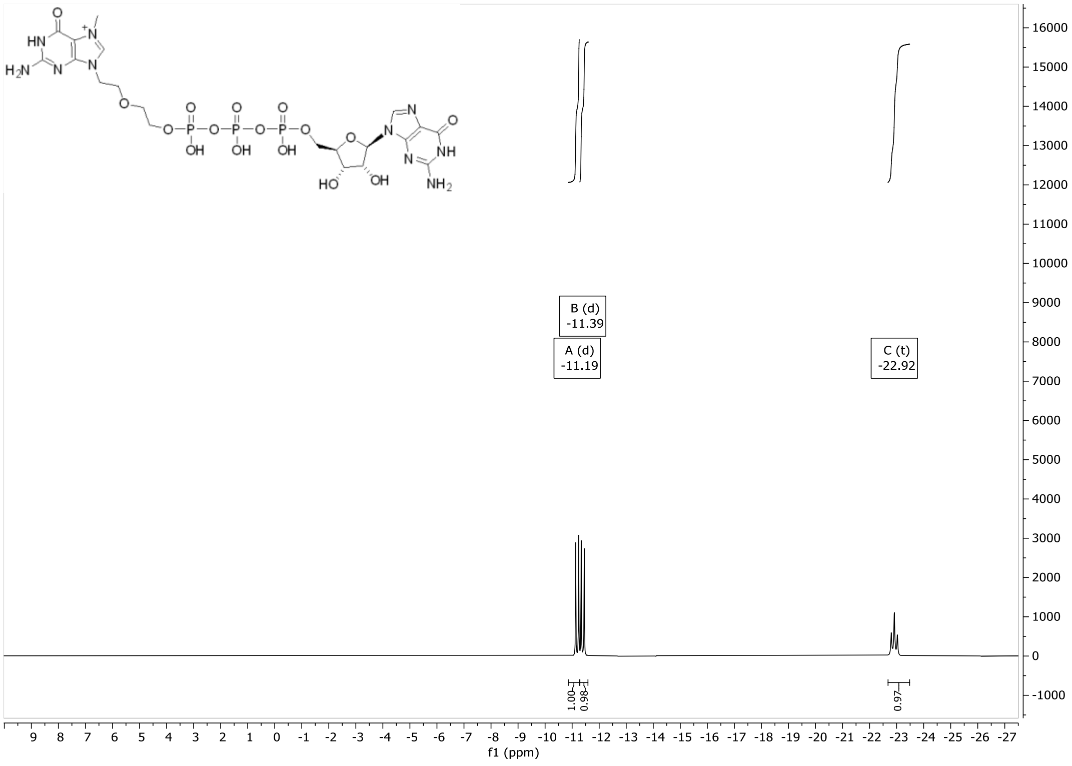

$^{Bz}$A(2'-OMe)-p-$^{iBu}$G(2,3'-OAc) (**7**)

$^{1}$H-NMR

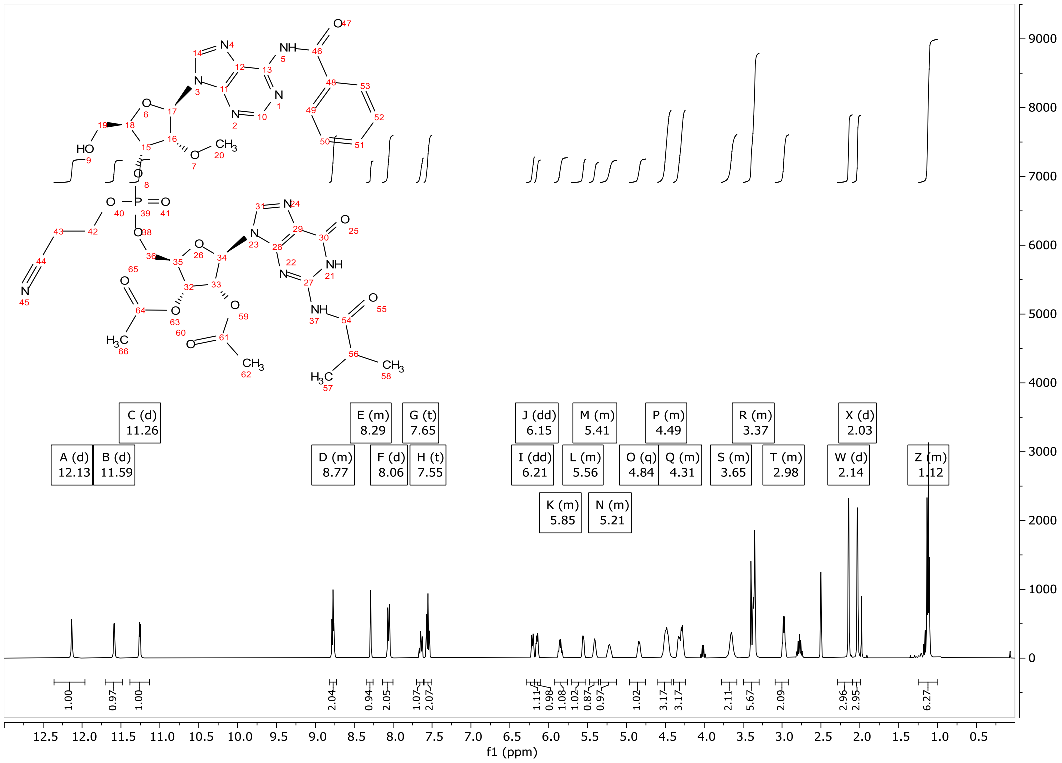

$^{31}$P-NMR

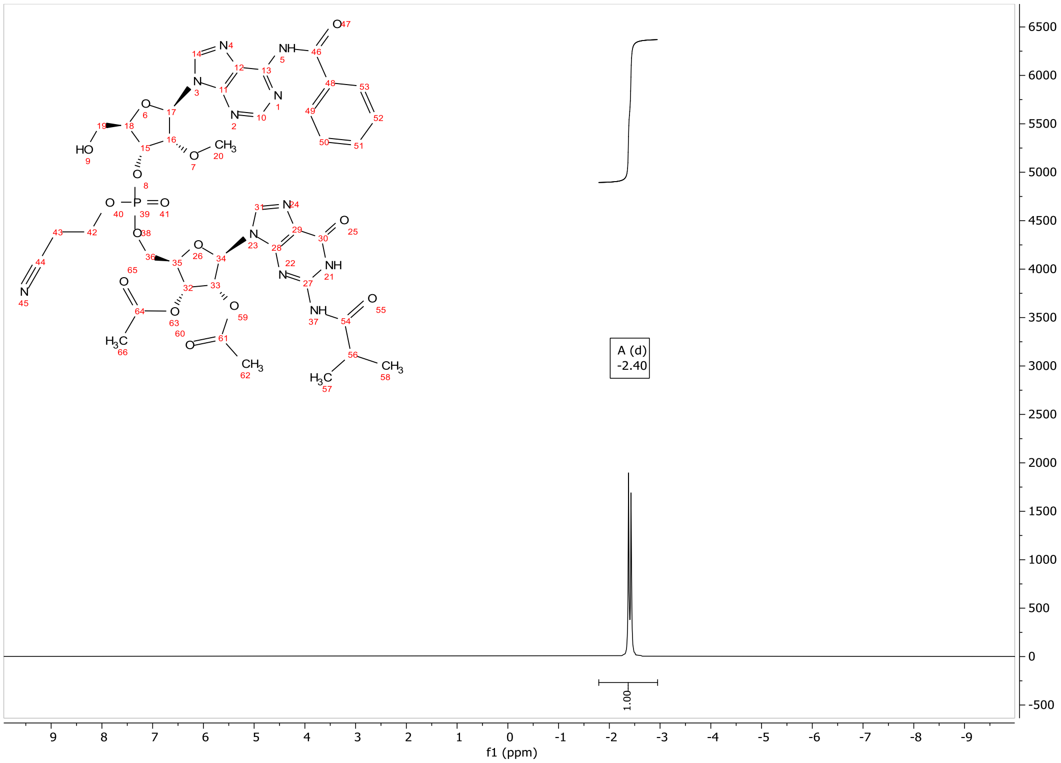

pA(2’-OMe)-G (**8**)

$^{1}$H-NMR

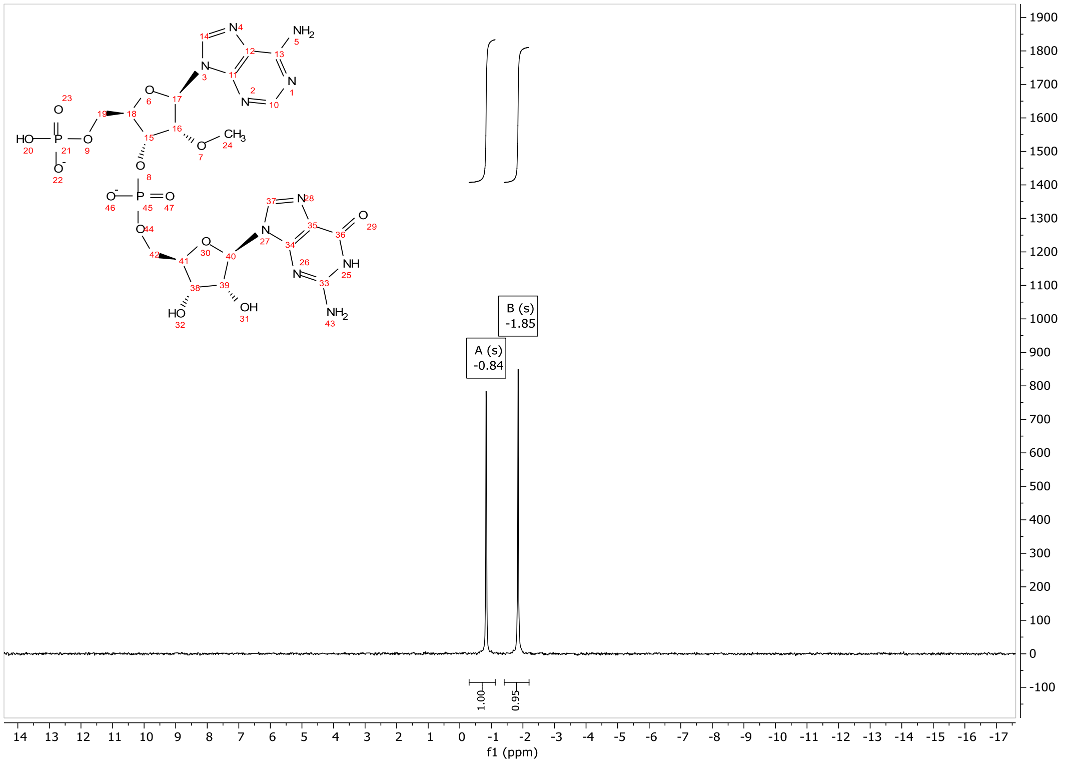

$^{31}$P-NMR

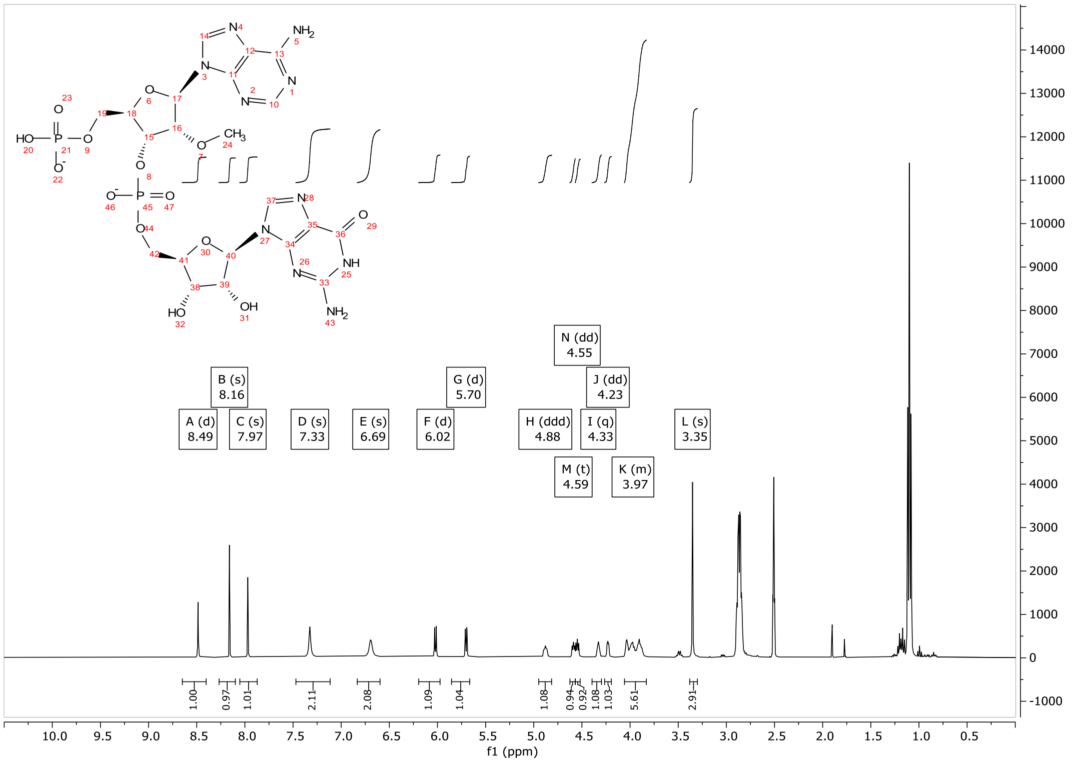

ppA(2'-OMe)-G (**9**)

$^1$H-NMR

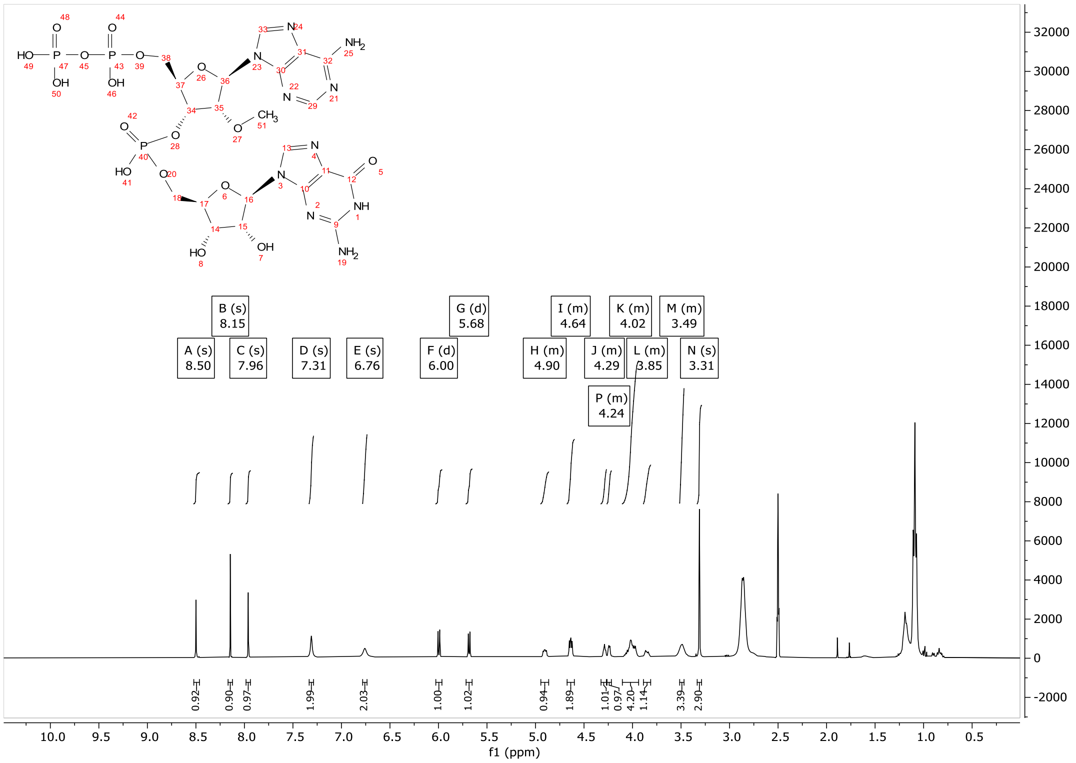

$^{31}$P-NMR

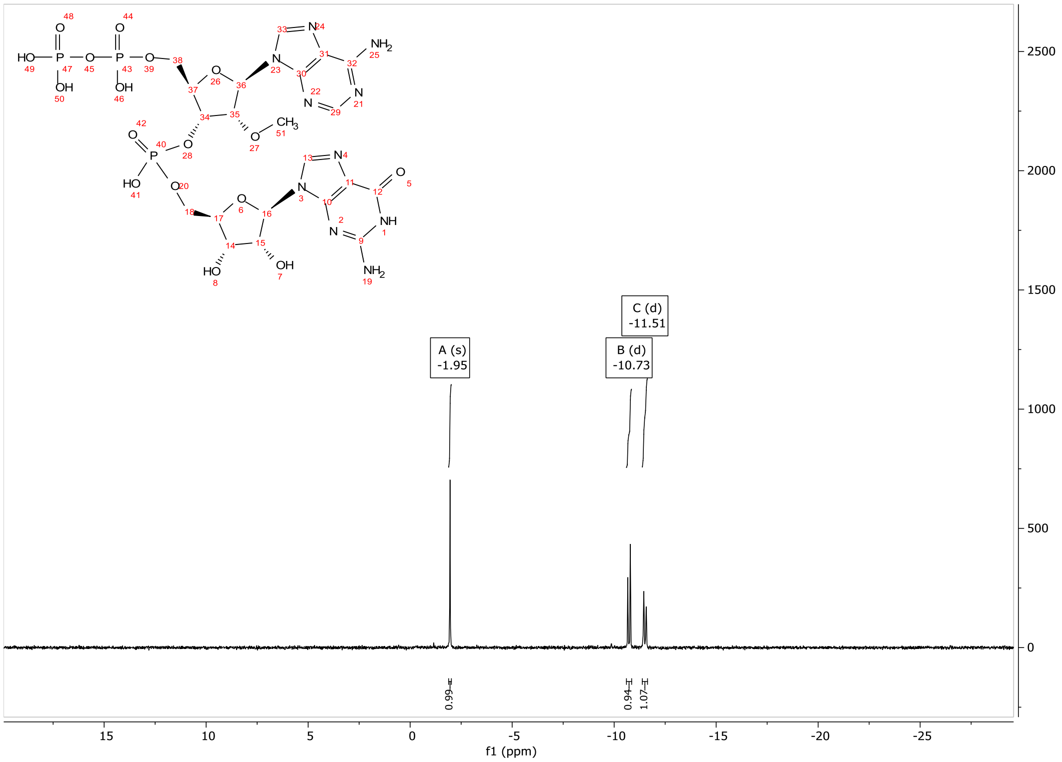

CV-Cap2

$^{1}$H-NMR

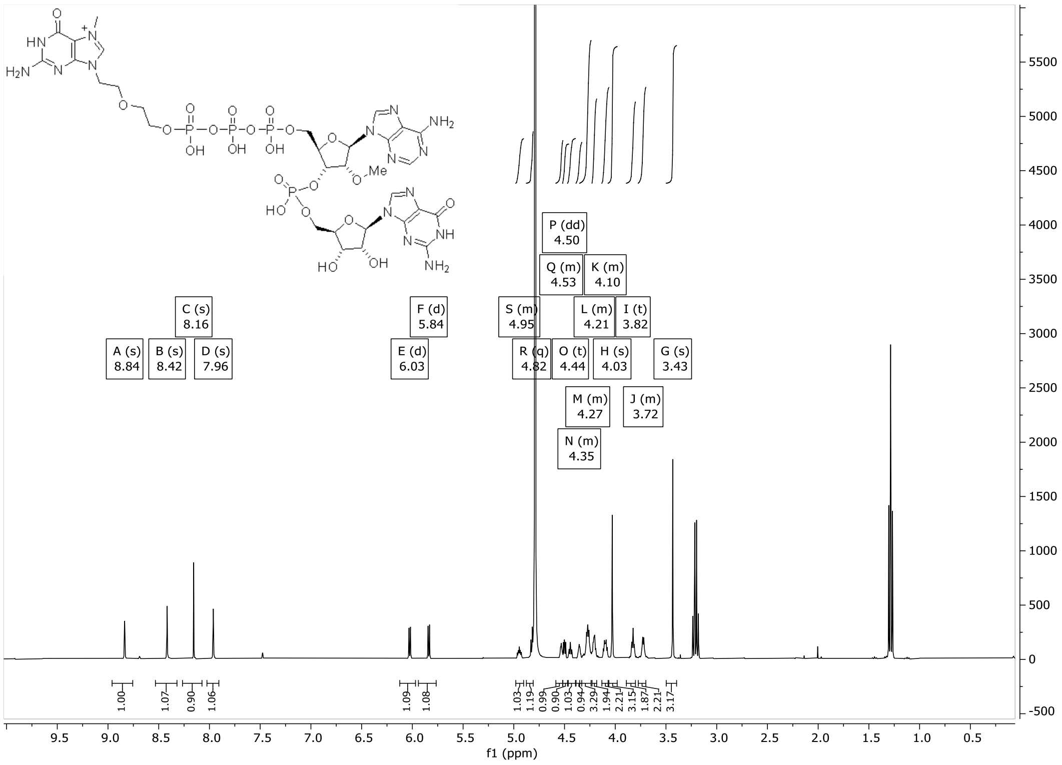

$^{31}$P-NMR

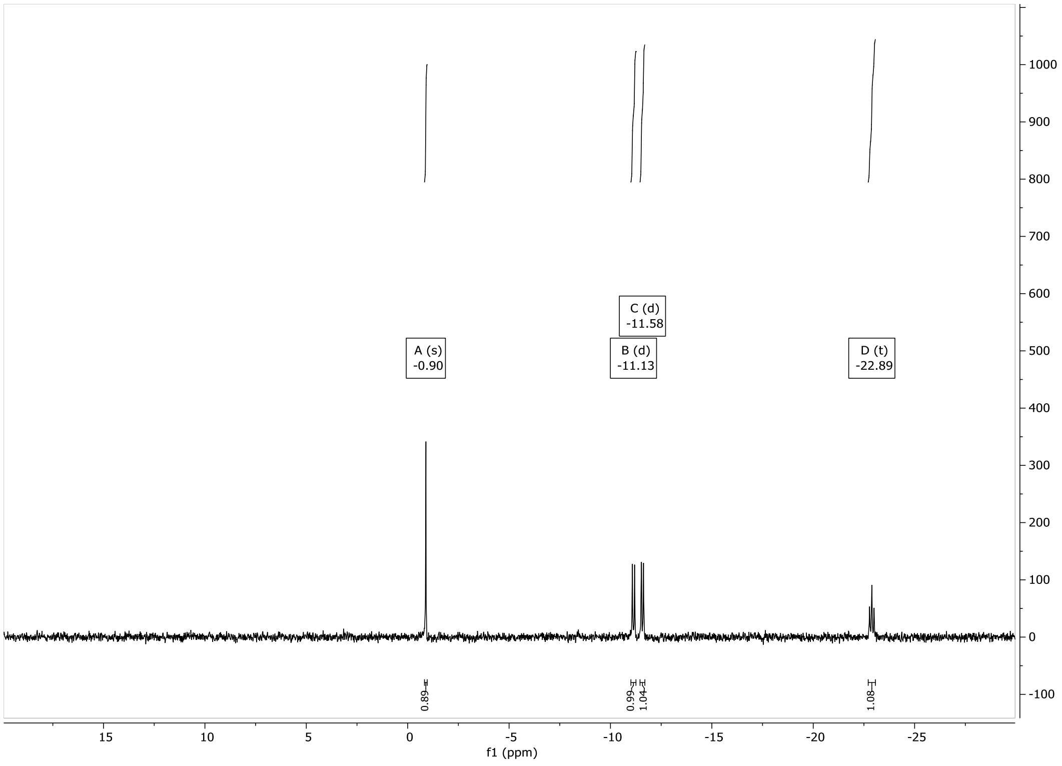

3′,5′-O-(1,1,3,3-tetraisopropyl-1,3-disiloxanediyl)-9-β-D-arabinofuranosyladenine (**10**)

$^{1}$H-NMR

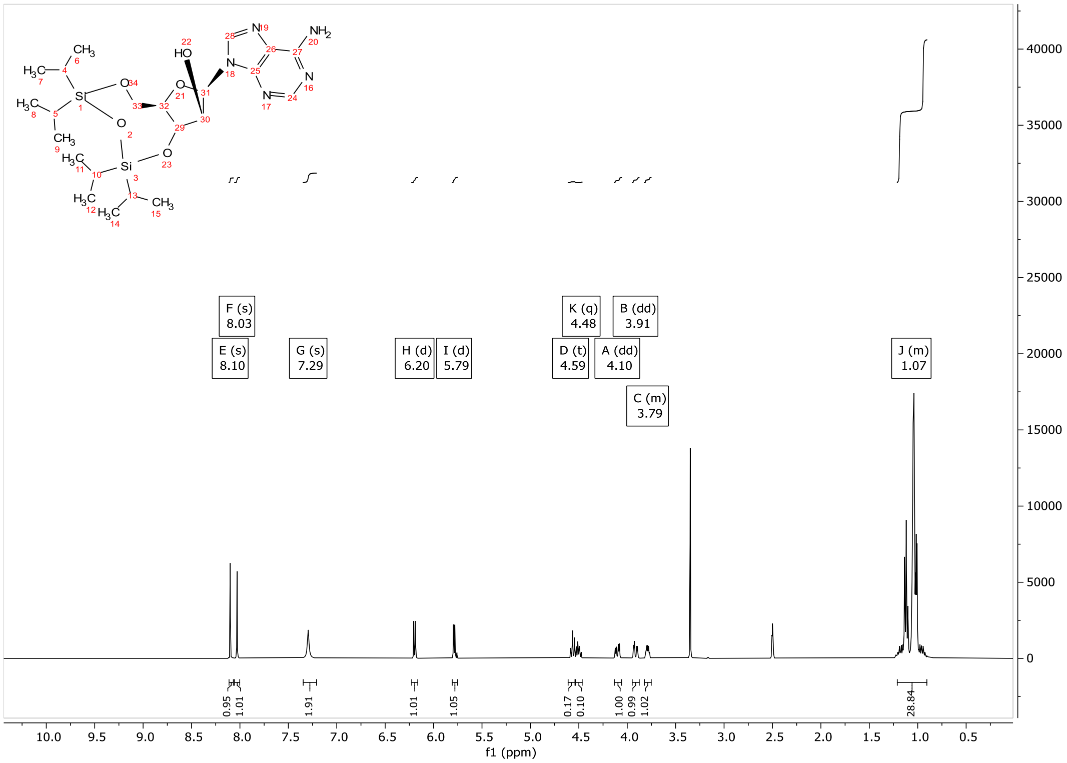

3′,5′-O-(1,1,3,3-tetraisopropyl-1,3-disiloxanediyl)-9-β-D-arabinofuranosyl-N6-benzoyl-adenine (**11**)

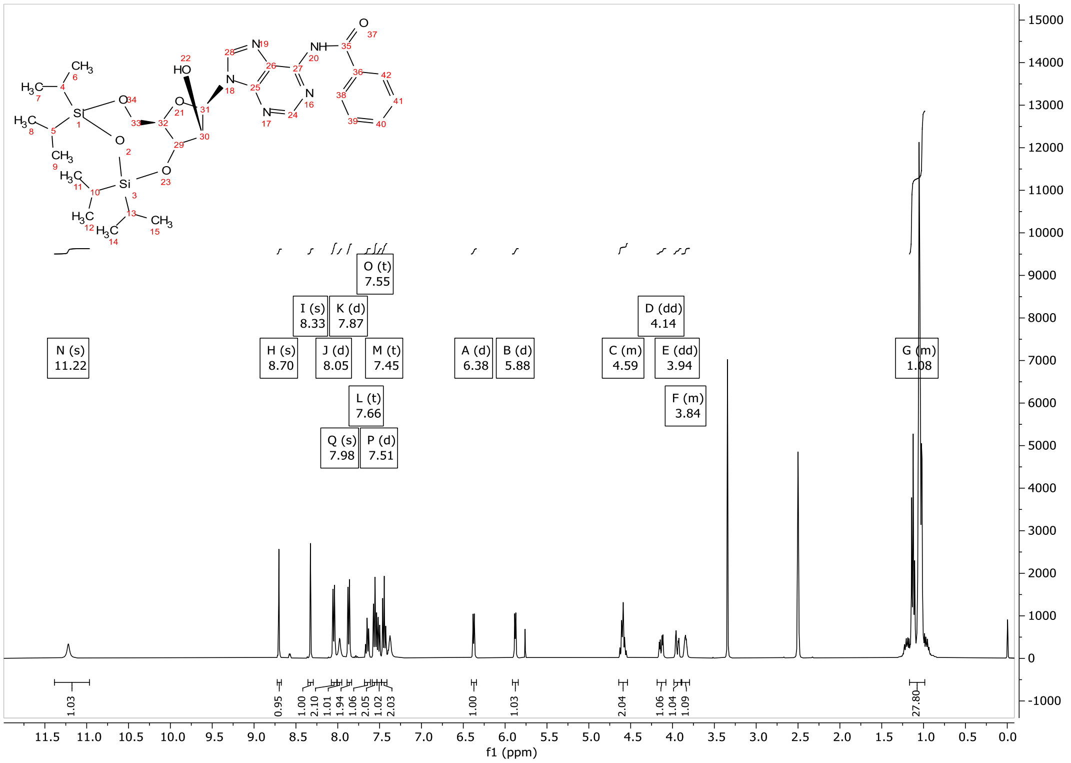

3′,5′-O-(1,1,3,3-tetraisopropyl-1,3-disiloxanediyl)-2’-mercaptomethyl-9-β-D-N6-benzoyl-adenine (**13**)

$^{1}$H-NMR

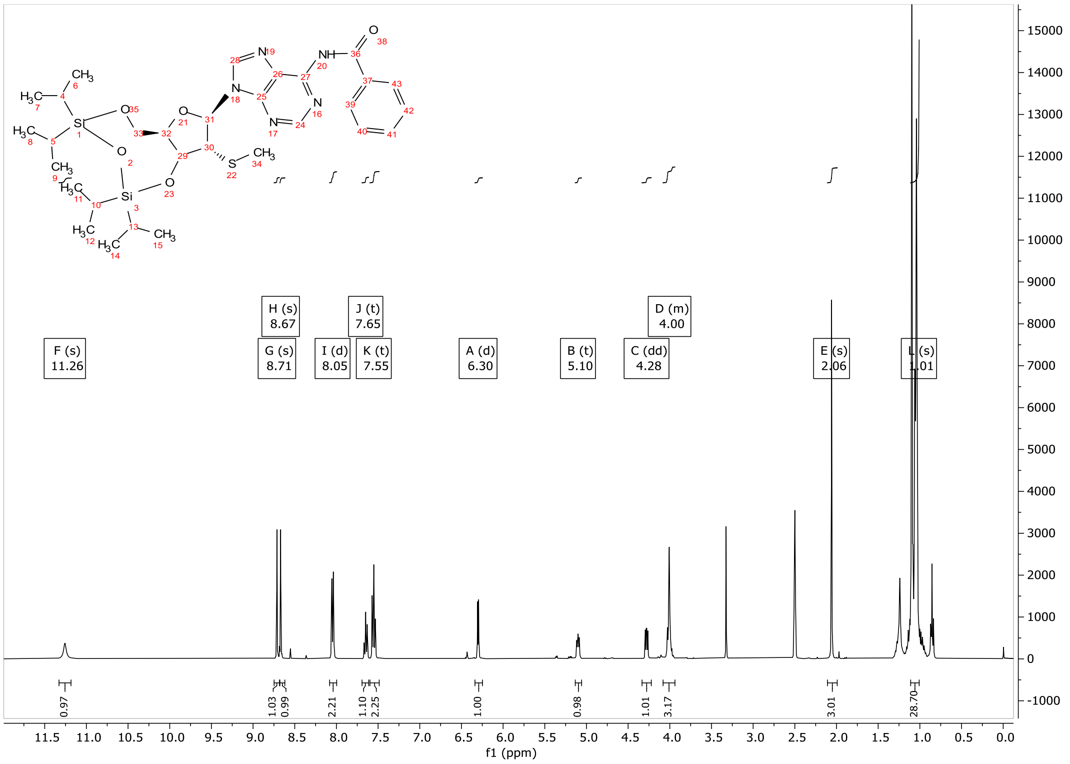

2’-mercaptomethyl-9-β-D-N6-benzoyl-adenine (**14**)

$^{1}$H-NMR

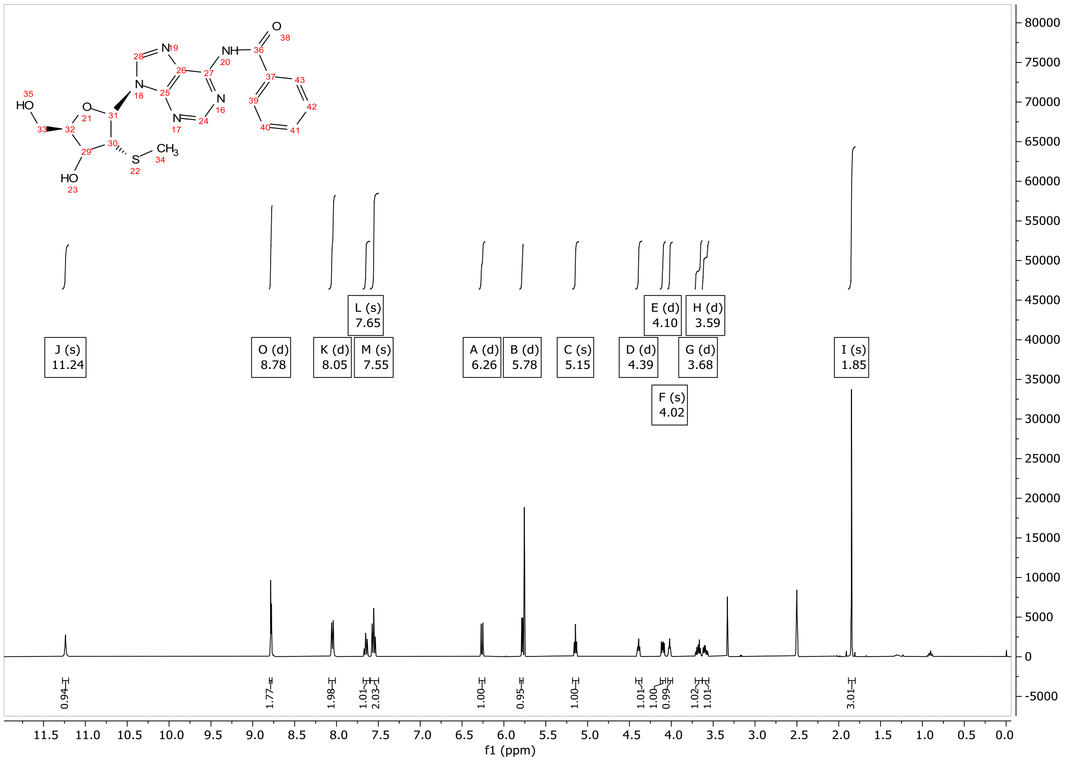

5’-Dimethoxytrityl-2’-mercaptomethyl-9-β-D-N6-benzoyl-adenine (**15**)

$^{1}$H-NMR

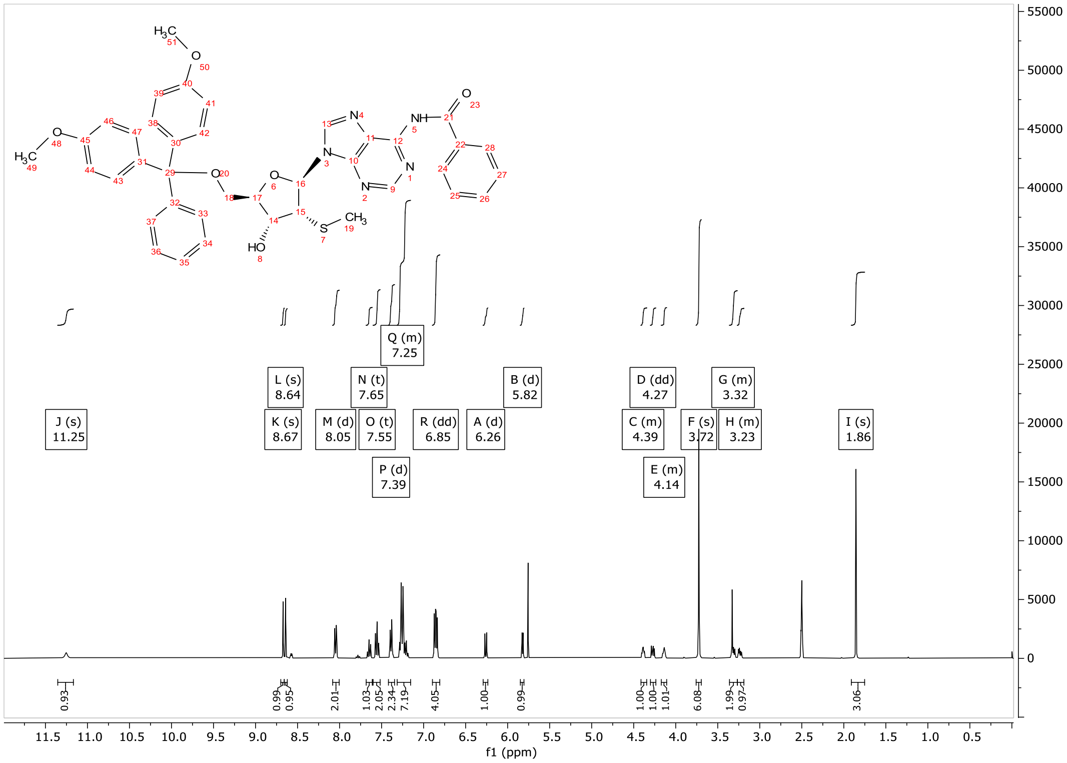

5’-Dimethoxytrityl-2’-mercaptomethyl-9-β-D-N6-benzoyl-adenosyl-3’-phosphoramidite (**16**)

$^{1}$H-NMR

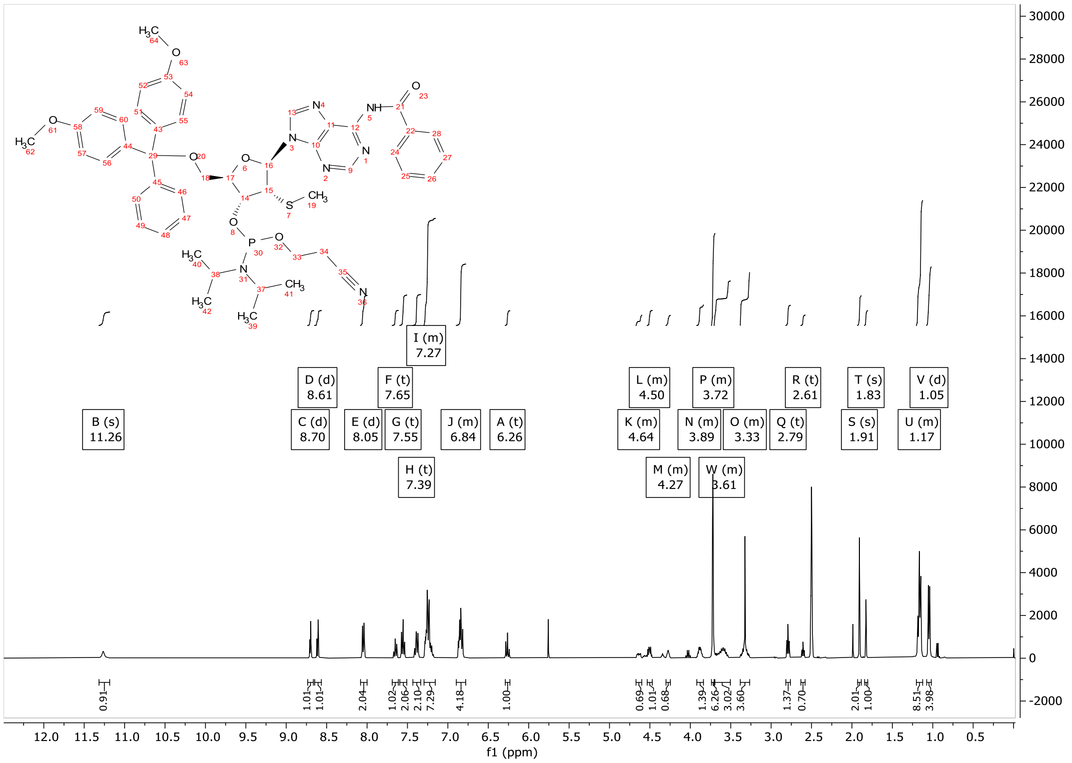

$^{31}$P-NMR

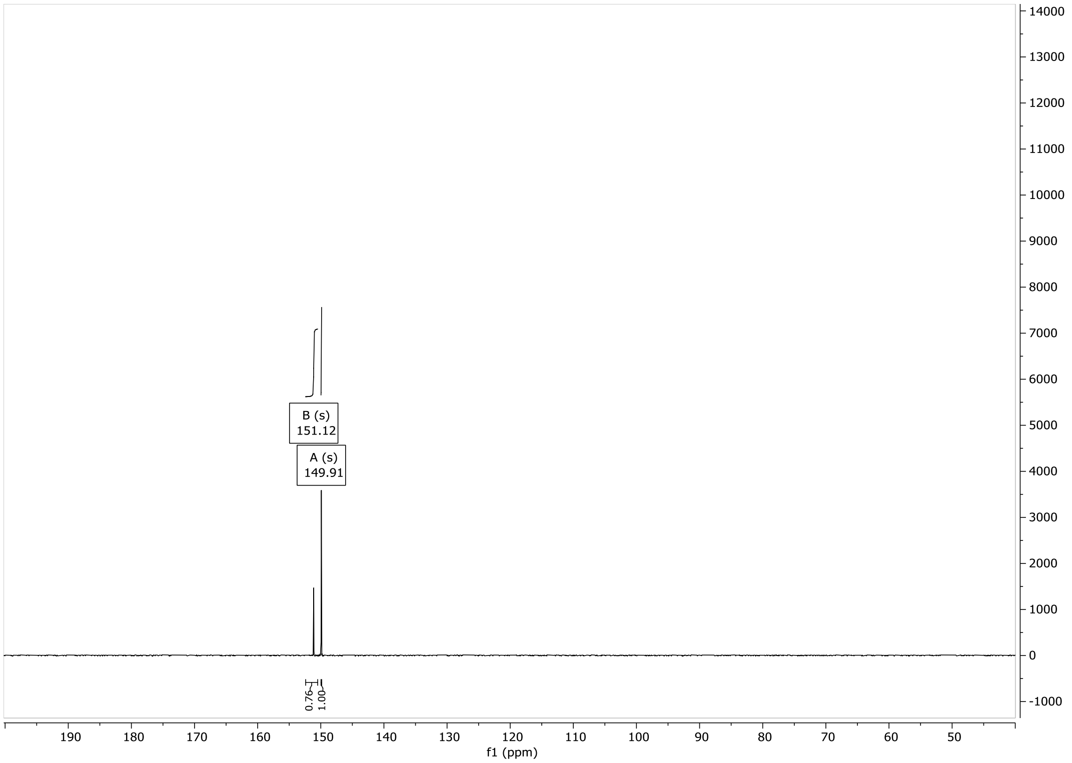

### A(2′-SMe)-p-G (**17**)

#### $^{1}$H-NMR

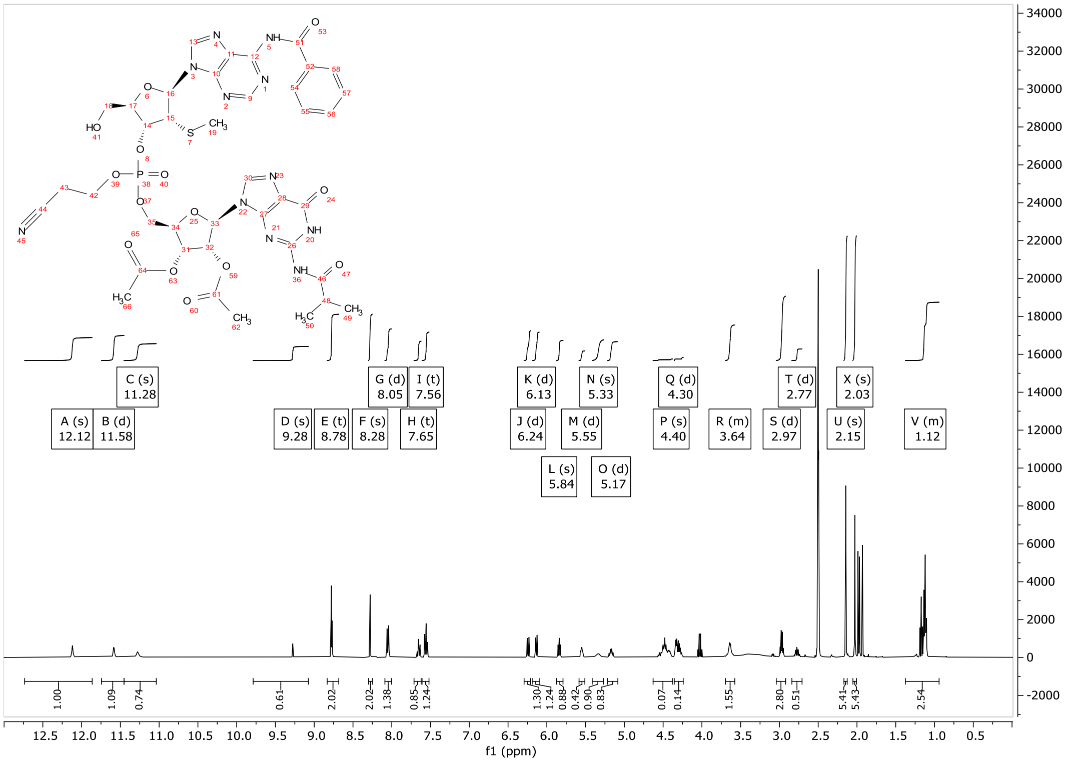

#### $^{31}$P-NMR

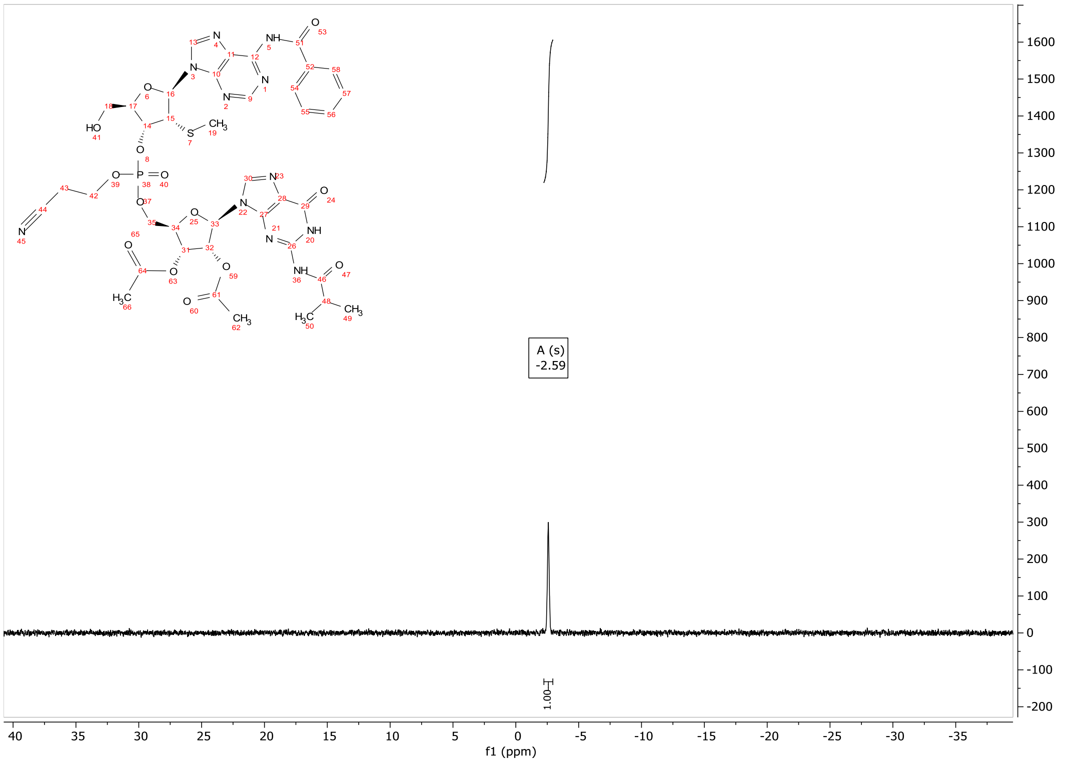

p-A(2′-SMe)-p-G (**18**)

$^{1}$H-NMR

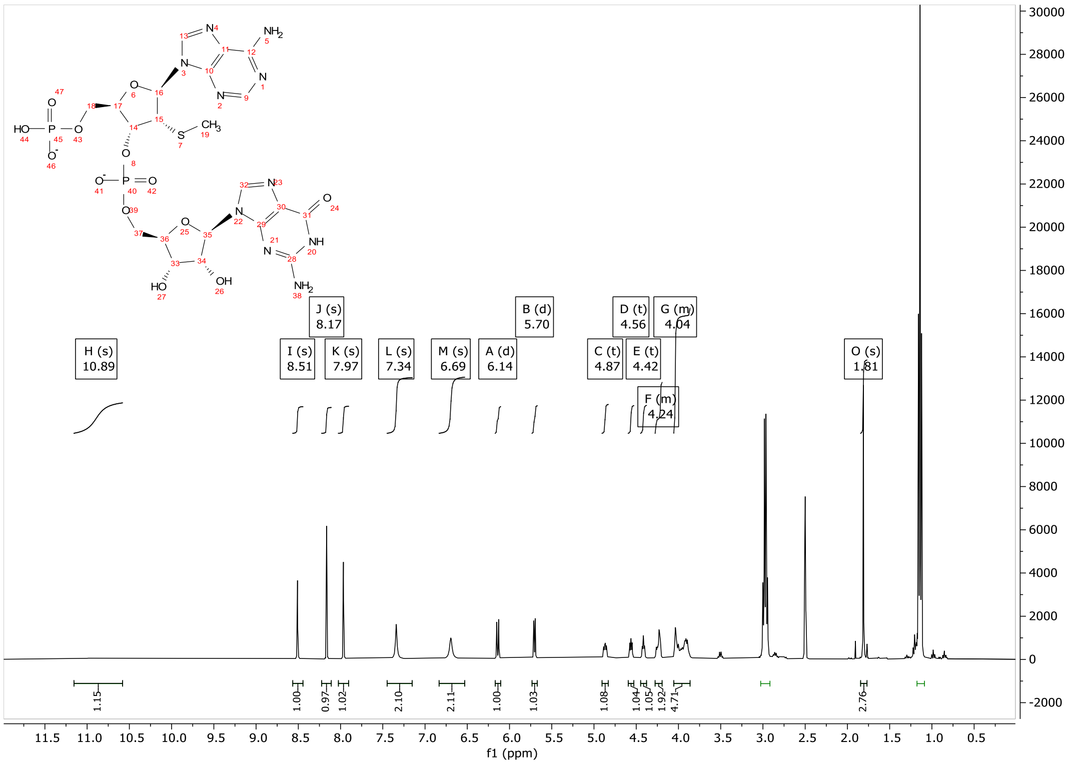

$^{31}$P-NMR

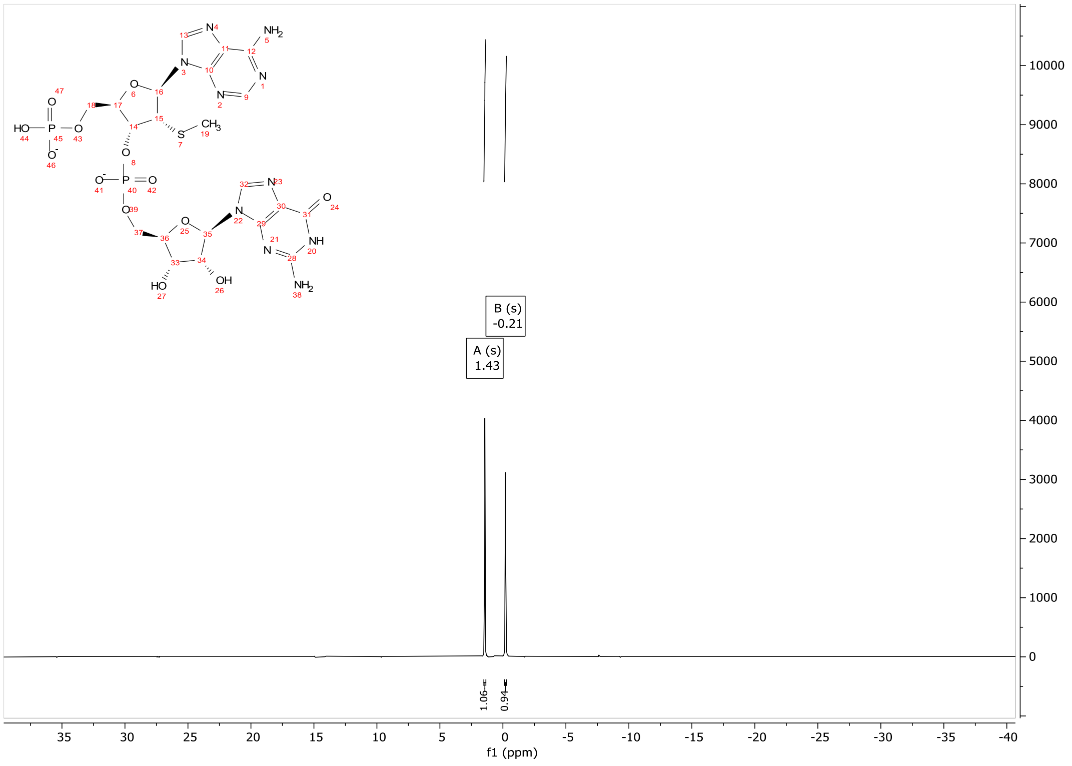

### CV-Cap3

#### $^1$H-NMR

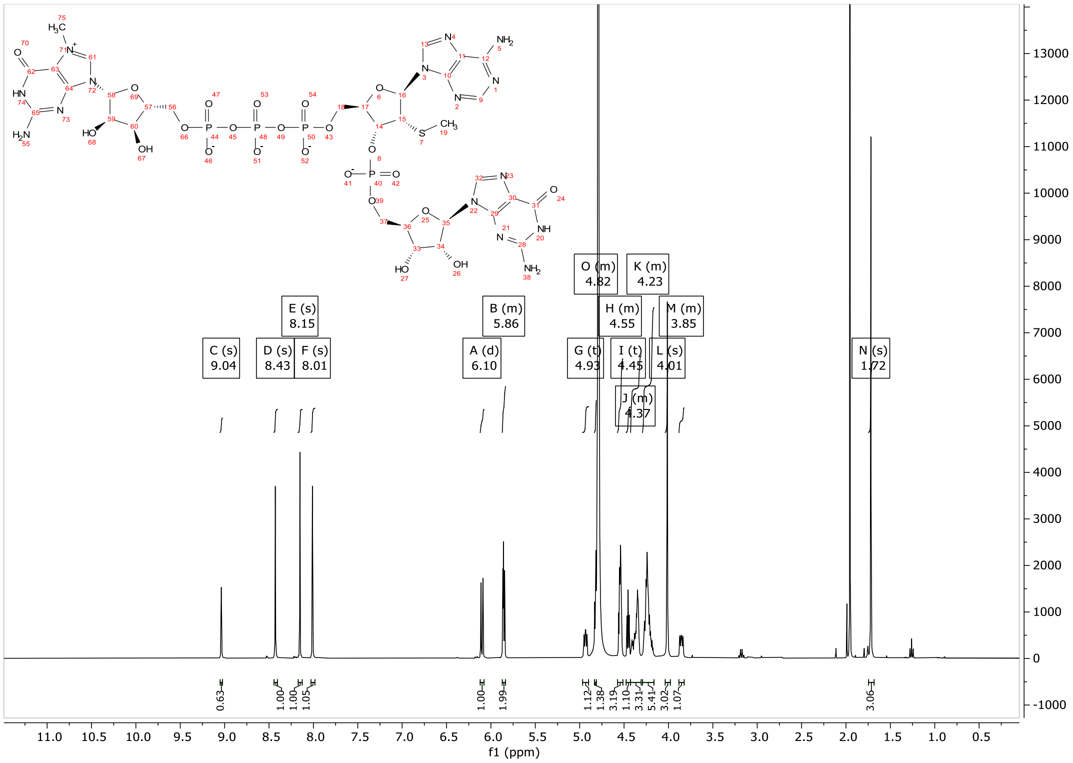

#### $^{31}$P-NMR

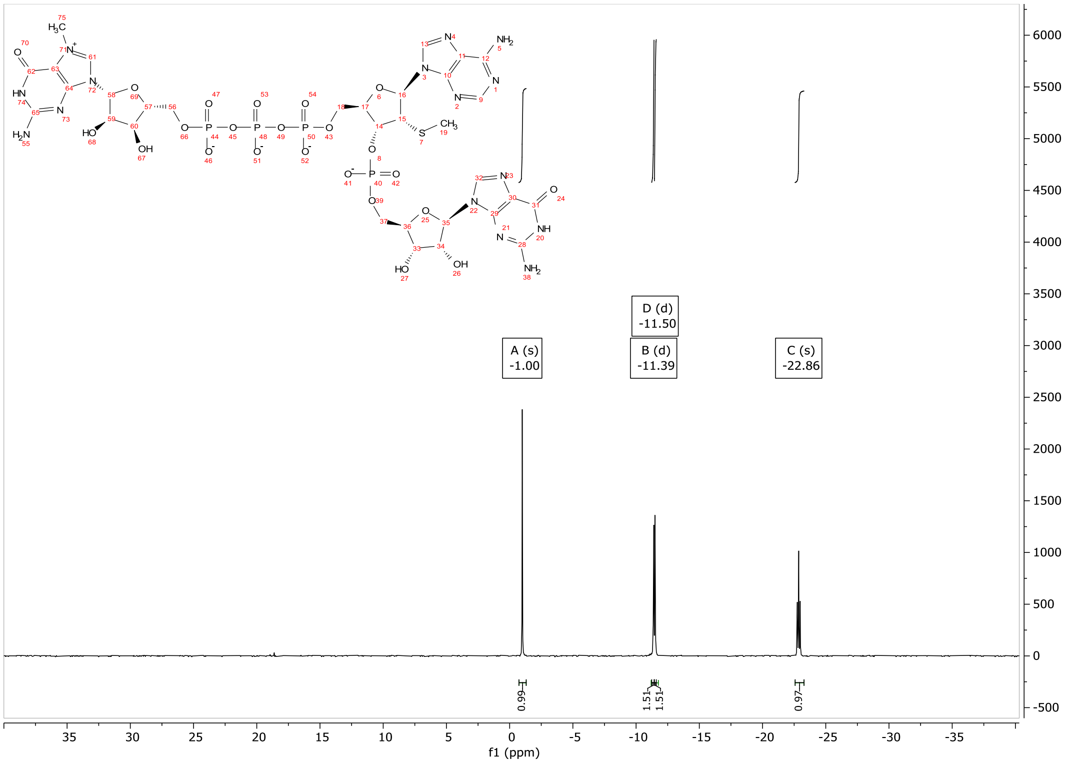

2′-O,4′-C-methylene-guanosine diphosphate (**19**)

$^{1}$H-NMR

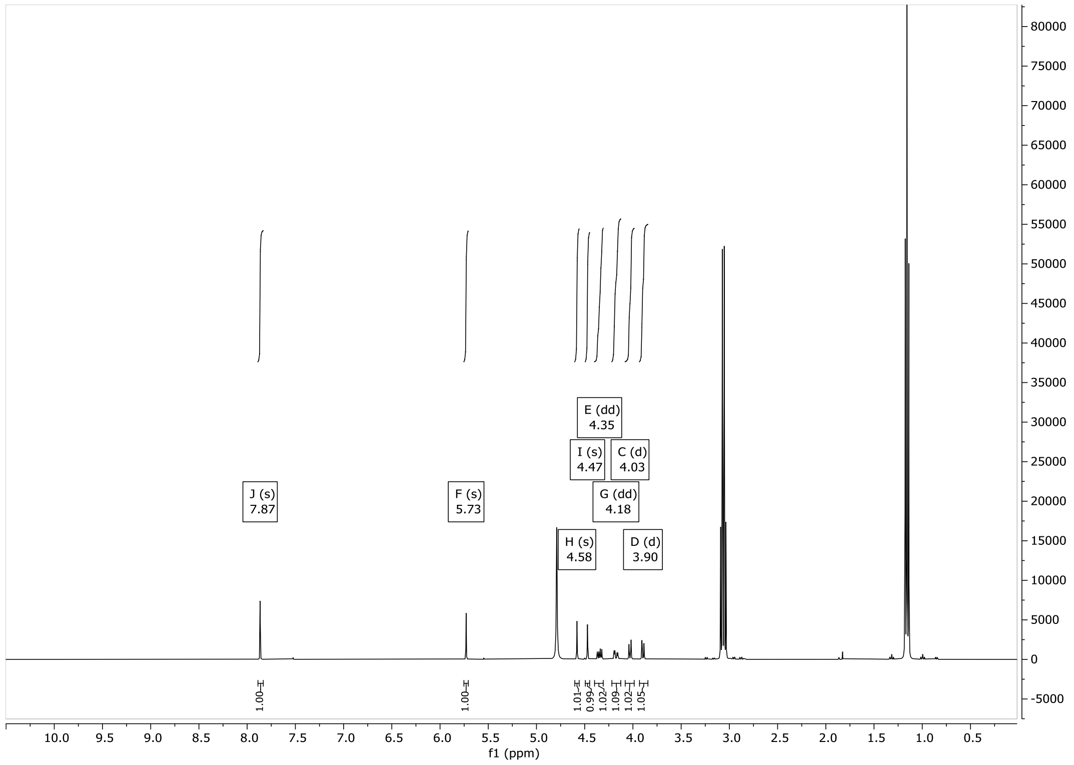

$^{31}$P-NMR

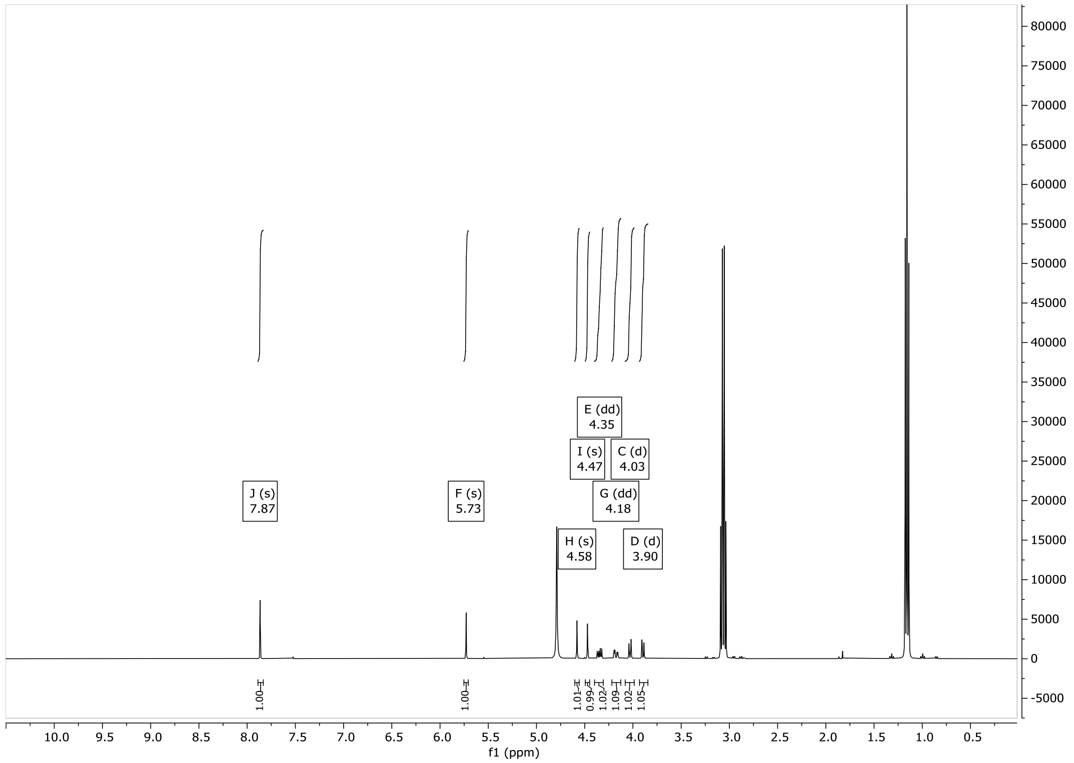

N7-methyl-2′-O,4′-C-methylene guanosine diphosphate (**20**)

$^{1}$H-NMR

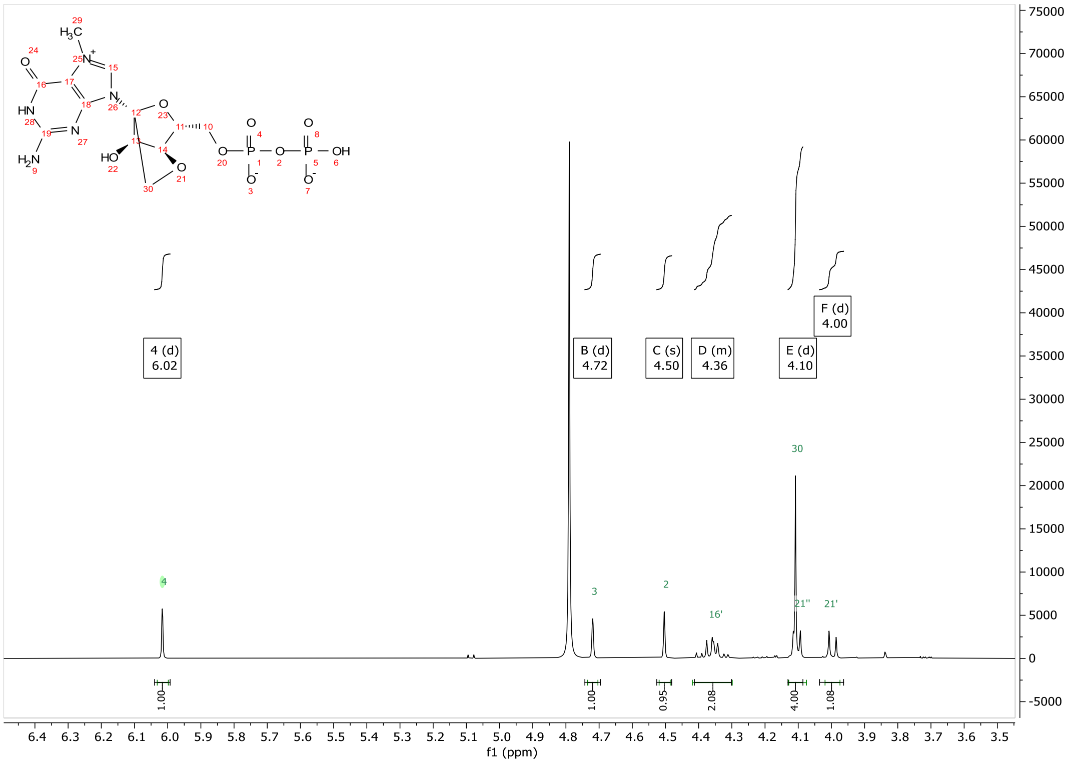

$^{31}$P-NMR

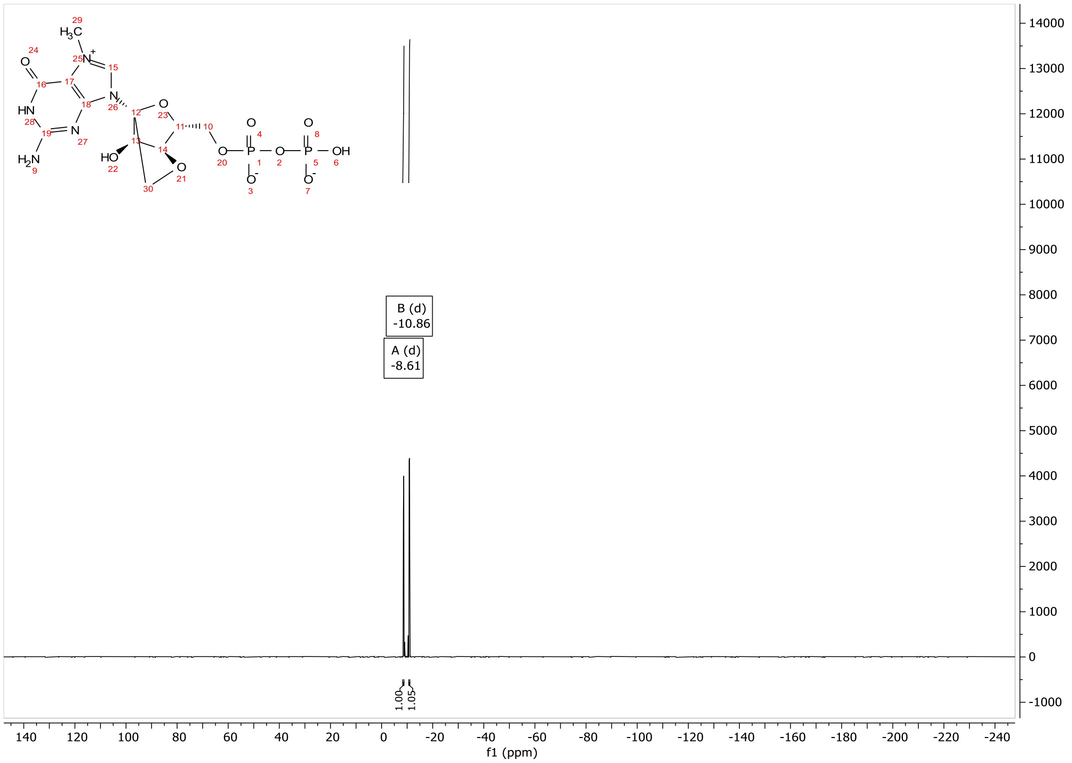

CV-Cap4

1H-NMR

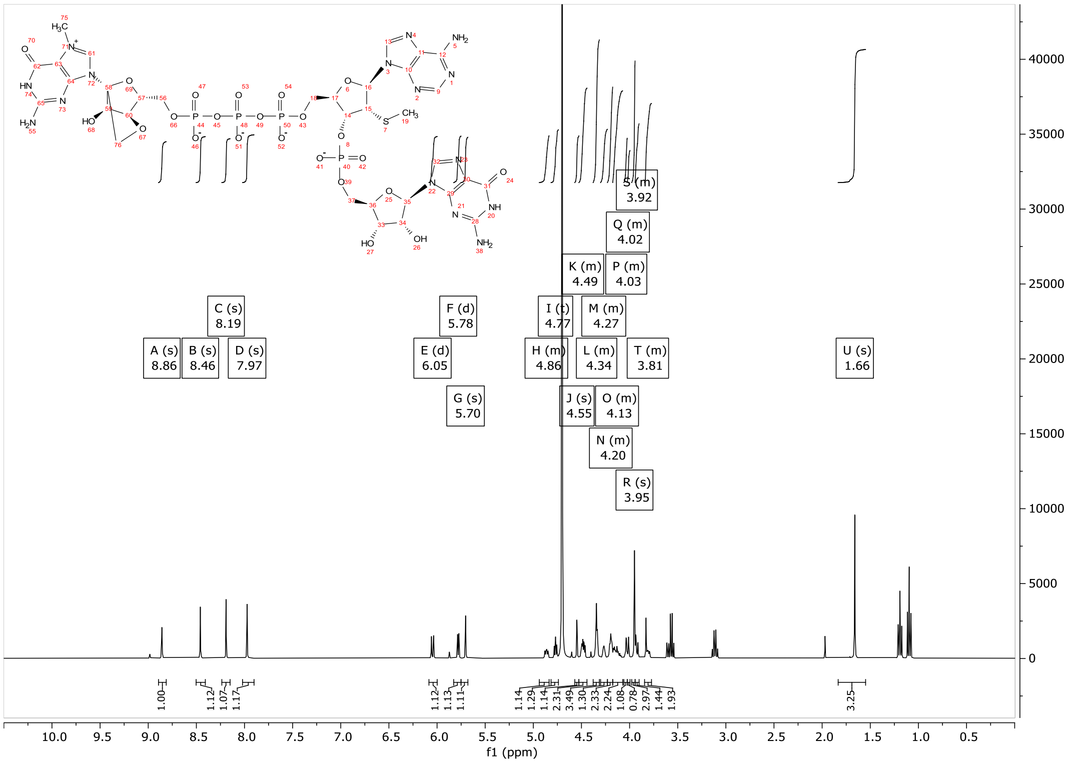

31P-NMR

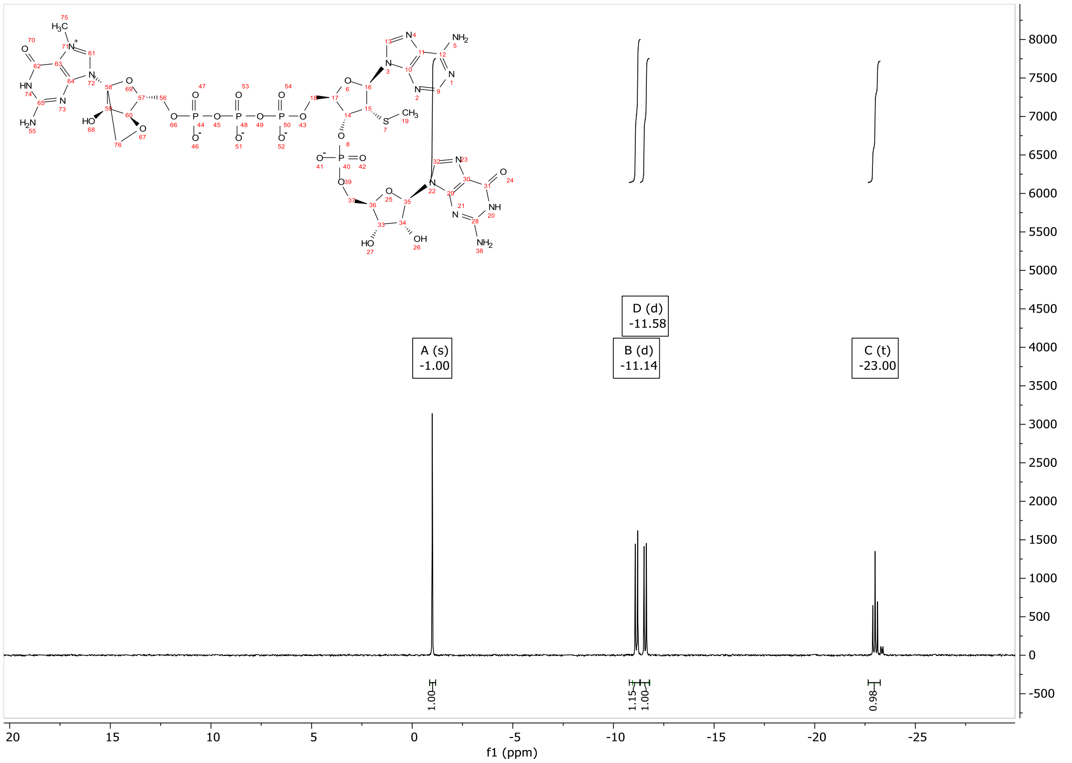

### BzA(2’-SMe)-p-iBuG(2,3’-OAc) (**22**)

#### $^1$H-NMR

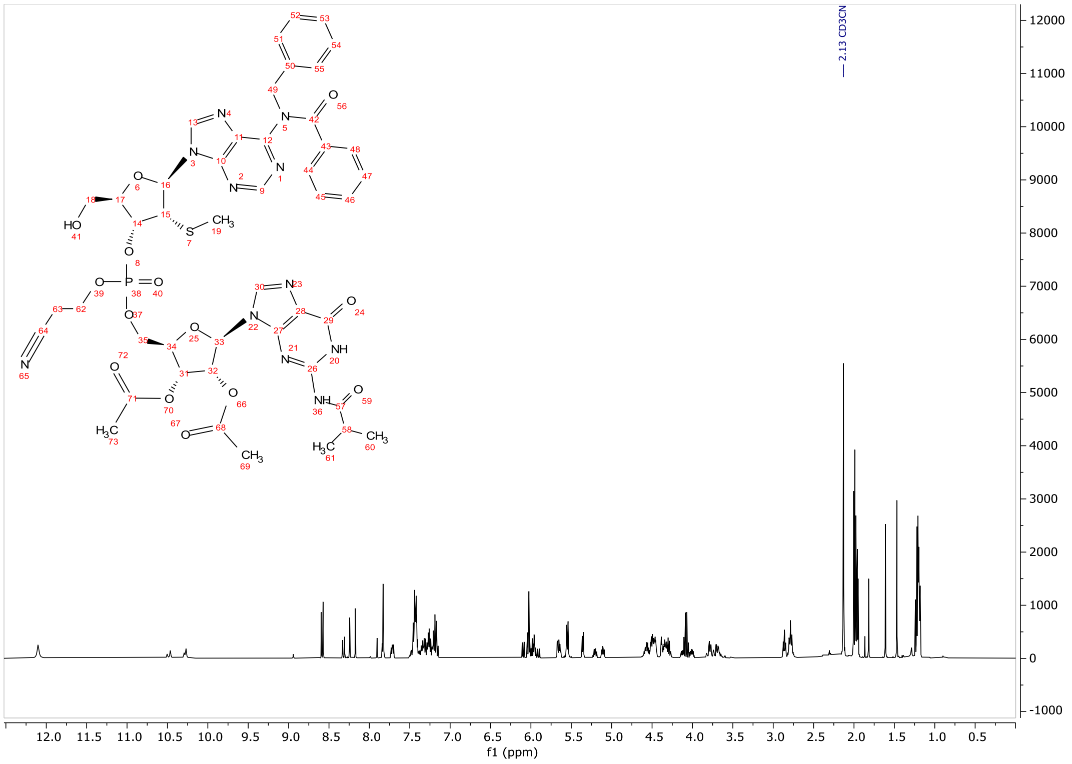

#### $^{31}$P-NMR

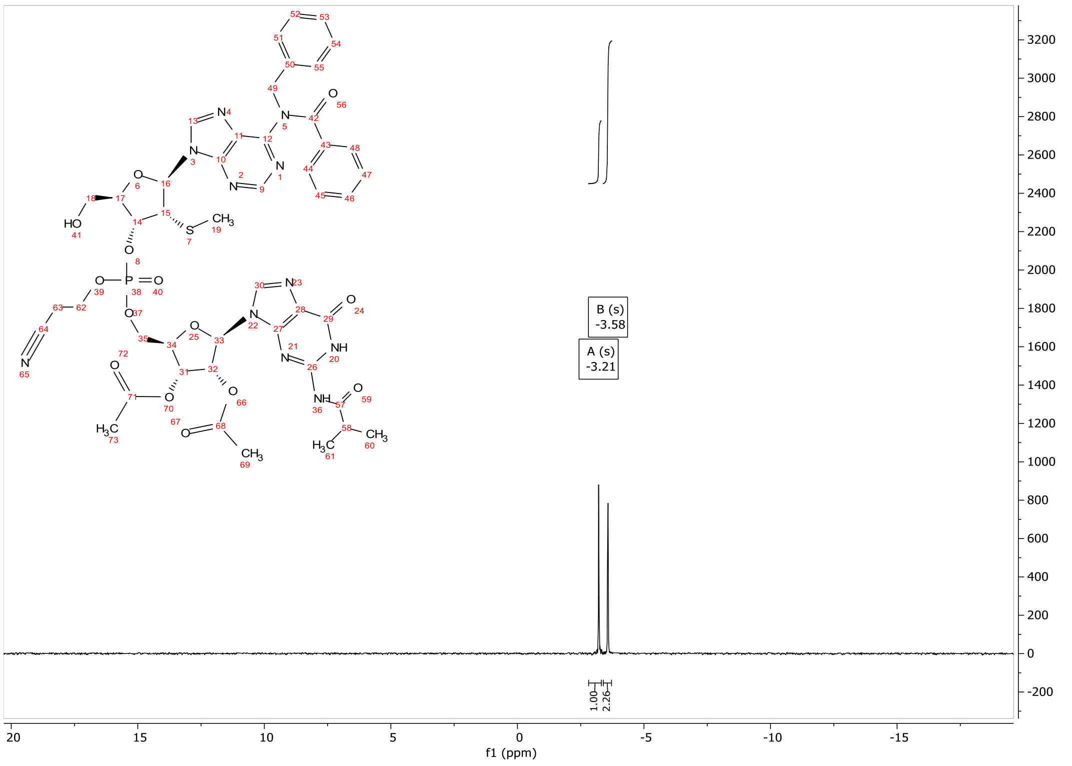

### p-N6-Benzyl-A(2′-SMe)-p-G (**23**)

#### $^{1}$H-NMR

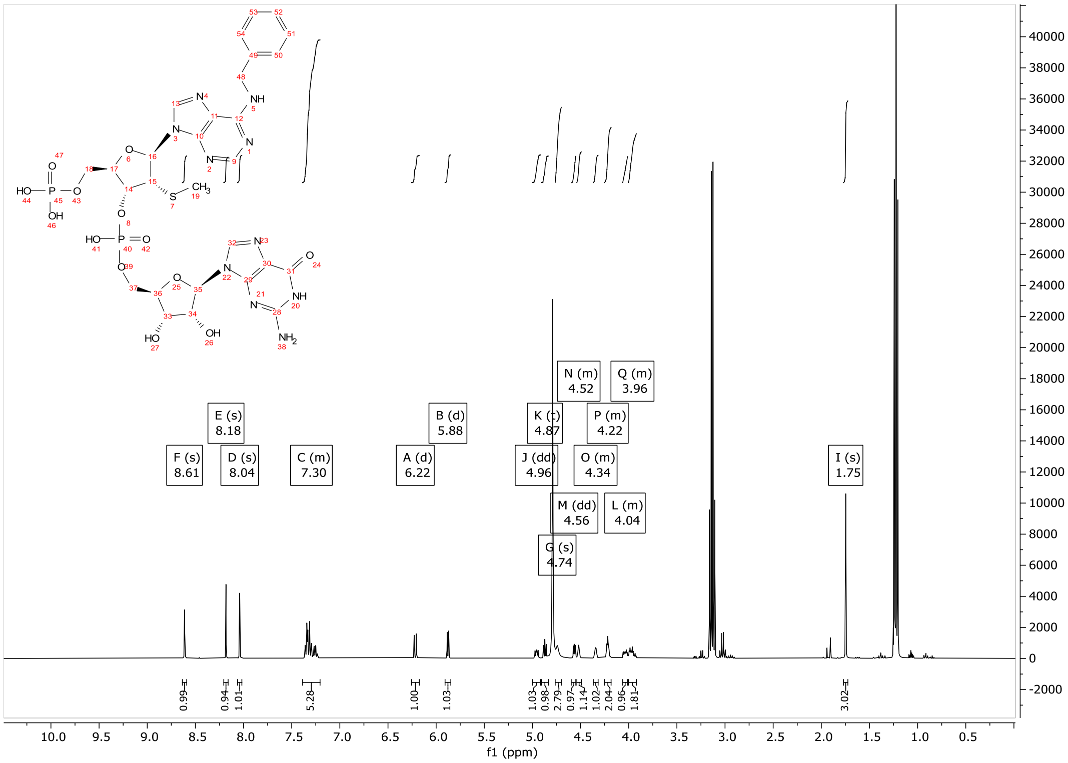

#### $^{31}$P-NMR

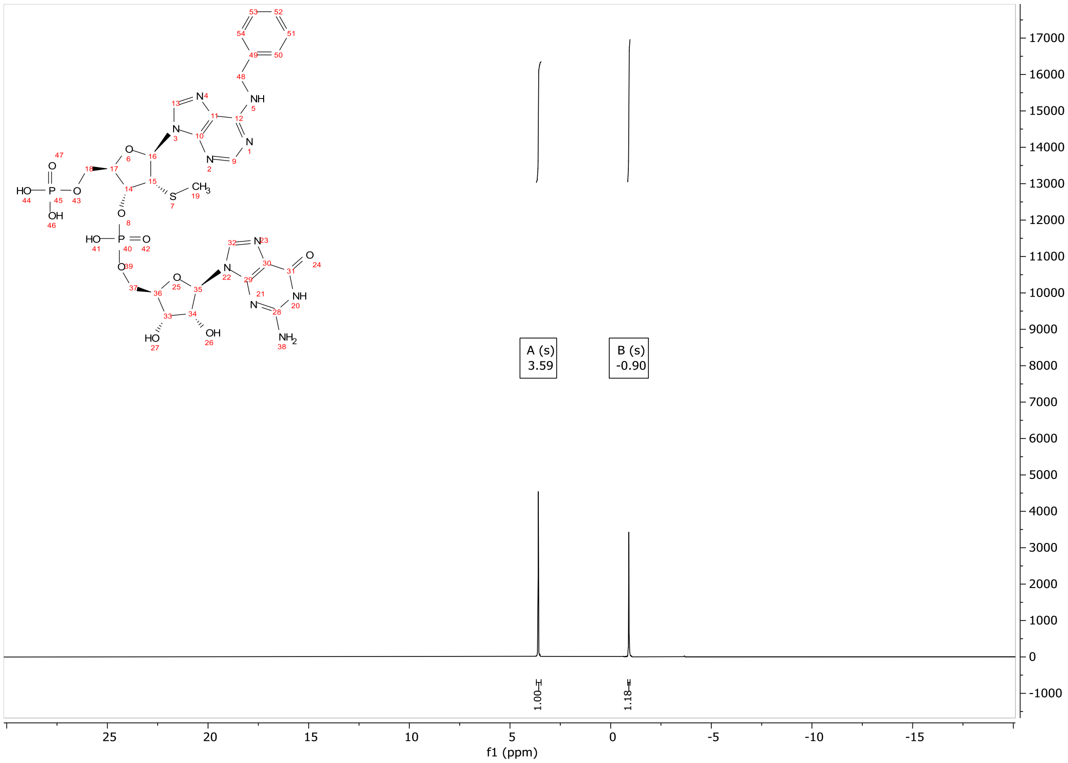

CV-Cap5

$^{1}$H-NMR

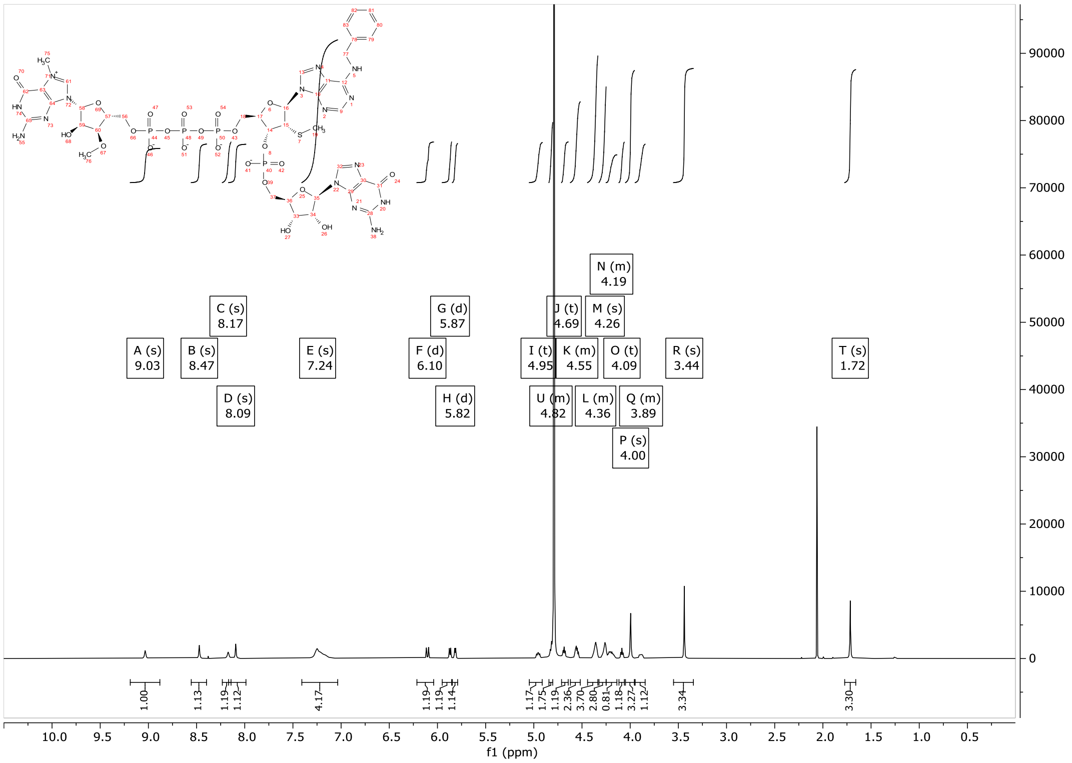

$^{31}$P-NMR

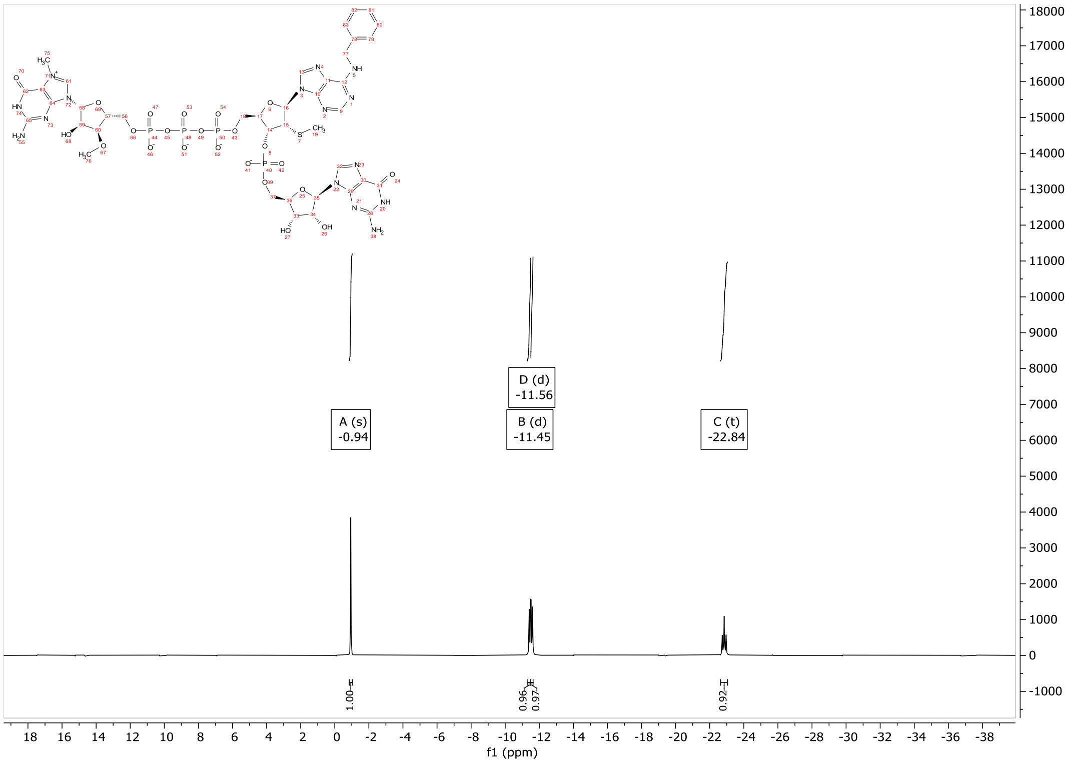
